# Undinarchaeota illuminate the evolution of DPANN archaea

**DOI:** 10.1101/2020.03.05.976373

**Authors:** Nina Dombrowski, Tom A. Williams, Jiarui Sun, Benjamin J. Woodcroft, Jun-Hoe Lee, Bui Quang Minh, Christian Rinke, Anja Spang

**Author notes:** corresponding author. Postal address: Landsdiep 4, 1797 SZ ’t Horntje (Texel).

## Abstract

The evolution and diversification of Archaea is central to the history of life on Earth. Cultivation-independent approaches have revealed the existence of at least ten archaeal lineages whose members have small cell and genome sizes and limited metabolic capabilities and together comprise the tentative DPANN archaea. However, the phylogenetic diversity of DPANN and the placement of the various lineages of this group in the archaeal tree remain debated. Here, we reconstructed additional metagenome assembled genomes (MAGs) of a thus far uncharacterized archaeal phylum-level lineage UAP2 (*Candidatus* Undinarchaeota) affiliating with DPANN archaea. Comparative genome analyses revealed that members of the Undinarchaeota have small estimated genome sizes and, while potentially being able to conserve energy through fermentation, likely depend on partner organisms for the acquisition of vitamins, amino acids and other metabolites. Phylogenomic analyses robustly recovered Undinarchaeota as a major independent lineage between two highly supported clans of DPANN: one clan comprising Micrarchaeota, Altiarchaeota and Diapherotrites, and another encompassing all other DPANN. Our analyses also suggest that DPANN archaea may have exchanged core genes with their hosts by horizontal gene transfer (HGT), adding to the difficulty of placing DPANN in the archaeal tree. Together, our findings provide crucial insights into the origins and evolution of DPANN archaea and their hosts.

## Introduction

Archaea represent one of the two primary domains of life^1–3^ and are thought to have played a major role in the evolution of life and origin of Eukaryotes^4–6^. While most archaea remain uncultivated, cultivation-independent approaches, such as single-cell and metagenomic sequencing, have revealed many previously unknown archaeal lineages in most environments on Earth and have changed our perception of archaeal functional and taxonomic diversity^7–10^. In particular, the Asgard^6^ and DPANN superphyla^11,12^ as well as a multitude of putative phylum-level lineages have been proposed in the archaeal domain over the last two decades but the phylogenetic relatedness and taxonomy of different archaeal lineages remain a matter of debate.

The DPANN radiation^12^, named after the first members of this group (Diapherotrites, Parvarchaeota, Aenigmarchaeota, Nanoarchaeota and Nanohaloarchaeota)^11^, comprises one of these recently proposed archaeal clades and is now thought to be comprised of at least ten and (according to NCBI taxonomy) putative phylum-level lineages^13,14^. Most members of the DPANN archaea are characterized by small cell sizes and reduced genomes, which code for a limited set of metabolic proteins^13^. The few members that have been successfully enriched in co-culture were shown to represent obligate ectosymbionts dependent on archaeal hosts for growth and survival. For instance, members of Nanoarchaeota are ectosymbionts of TACK archaea^15–20^, Micrarchaeota were found in co-culture with Thermoplasmates^21,22^ and Nanohaloarchaeota are dependent on halobacterial hosts^23^. Furthermore, evidence from FISH and co-occurrence analyses have suggested that Huberarchaeota may be ectosymbionts of members of the Altiarchaeota^24,25^. Yet, for most DPANN representatives, the identity of their symbiotic partners remains unclear.

Ever since the discovery of the first DPANN representative - *Nanoarchaeum equitans*, an ectosymbiont of *Ignicoccus hospitalis*^15^ - the phylogenetic placement of putative DPANN clades in the archaeal tree have been uncertain^26^. While various phylogenetic analyses have indicated that DPANN may comprise a monophyletic radiation in the Archaea^10,11,27^, these have been debated^8,28,29^. In particular, analyses focusing on the placement of selected DPANN lineages in isolation, such as Nanoarchaeota and Parvarchaeota, relative to other Archaea, have led to the conclusion that these represent fast-evolving Euryarchaeota^28,29^. Furthermore, it is debated whether the free-living Altiarchaeota belong to the DPANN radiation, form an independent lineage or belong to Euryarchaeota^8,9,14,30,31^. A potential cause for these conflicting topologies is that DPANN are often found on long branches in phylogenetic trees; these long branches might result from compositional biases or fast evolutionary rates^32,33^ (as seen for obligate bacterial endosymbionts^34,35^) or might reflect genomic under sampling of the true diversity of this group^9^. These alternatives are difficult to distinguish because, in the absence of fossils or definitive geochemical traces in the fossil record, we lack a well-constrained timescale for archaeal evolution. Distantly-related long-branching lineages can sometimes artificially group together on trees due to methodological artefacts, a phenomenon called long ranch attraction (LBA)^32^. Ways to ameliorate such artifacts include increased taxonomic sampling^36^, use of phylogenetic models less prone to LBA^37^, and the removal of fast-evolving or compositionally-biased sites from the alignment^38^. Furthermore, it seems possible that horizontal gene transfers (HGT) between symbionts and hosts^39^ could impede correct phylogenetic inferences if not accounted for.

Several recent studies have revealed the presence of a thus far uncharacterized archaeal lineage referred to the Uncultivated Archaeal Phylum 2 (UAP2)^13,40,41^, which seems to affiliate with DPANN archaea and thus may be key in resolving long-standing debates regarding archaeal phylogeny and the evolution of DPANN. In this study, we have used a metagenomics approach to obtain additional genomes of members of the so far uncharacterized UAP2 and provide first insights into their metabolic repertoire and lifestyle. Furthermore, we implemented comprehensive and careful phylogenomic techniques aimed at ameliorating phylogenetic artifacts, shedding new light onto the evolutionary origin and phylogenetic placement of the various DPANN lineages including UAP2 in an updated archaeal phylogeny.

## Results and Discussion

### A thus far uncharacterized putative archaeal phylum-level lineage in metagenome read archives

The generation of a large diversity of metagenome-assembled genomes (MAGs) representing archaeal and bacterial lineages across the tree of life has led to the definition of the tentative archaeal UAP2 phylum^41^. Considering our lack of insights into the biology of members of this lineage as well as its suggested key position in the archaeal tree, we aimed at obtaining a broader genomic representation of the UAP2. In particular, we screened publicly available metagenomes using ribosomal protein sequences of the previously reconstructed UAP2 MAGs and assembled and binned UAP2-positive samples yielding six additional MAGs belonging to the UAP2 lineage (**Table 1, Supplementary Tables 1-2**, see Methods for details). Four of the newly assembled MAGs were recovered from metagenomes of a groundwater aquifer located adjacent to the Colorado River^42^, while the two others as well as six previously reconstructed MAGs, derived from metagenomes of marine waters in the Atlantic^41,43^ and Indian Oceans^44^ as well as the Mediterranean Sea^45^ (**Supplementary Table 2**). UAP2 representatives were detected in samples from various depths in the water column (85-5000 m), fluctuating oxygen conditions (anoxic to oxic) and temperatures (sampling sites had temperatures from 18 up to 106°C) (**Supplementary Figure 1, Supplementary Table 1**). The MAGs, including previously published ones, are on average 78% complete (min: 55%, max: 91%) and show low signs of contamination (<5%) and strain heterogeneity (<2%). In total, they represent 2 high-quality (one from this study) and 10 medium-quality draft genomes according to genome reporting standards for MAGs assessed using a general archaeal marker protein set (**Table 1**, Supplementary Information)^46^. The UAP2 MAGs have small genomes with an average size of 0.66 MB, coding for an average of 750 proteins. They likely represent a distinct archaeal phylum-level lineage based on average amino acid identity (AAI) comparisons with other archaeal taxa (**Supplementary Figure 2, Supplementary Table 3**), phylogenetic analyses including a concatenated 16S-23S rRNA gene tree (**Supplementary Figures 3-5** and see below) as well as classification based on the Genome Taxonomy Database (GTDB) rank normalization (**Table 1, Supplementary Table S2**). Furthermore, the aquifer and marine UAP2 MAGs likely represent different orders as suggested by a relative evolutionary divergence (RED) value of 0.47^47^. Based on two high-quality UAP2 MAGs (Table 1, **Supplementary Table S2**)^46^, we propose two type species; *Candidatus* Undinarchaeum marinum (SRR4028224.bin17) and *Candidatus* Naiadarchaeum limnaeum (SRR2090159.bin1129), representing the marine and aquifer UAP2 clade, respectively (see details below). Undines are water elementals described in the writings of the alchemist Paracelsus, while Naiads were nymphs residing in ponds, springs and other bodies of fresh water in Greek mythology.

**Table 1.**
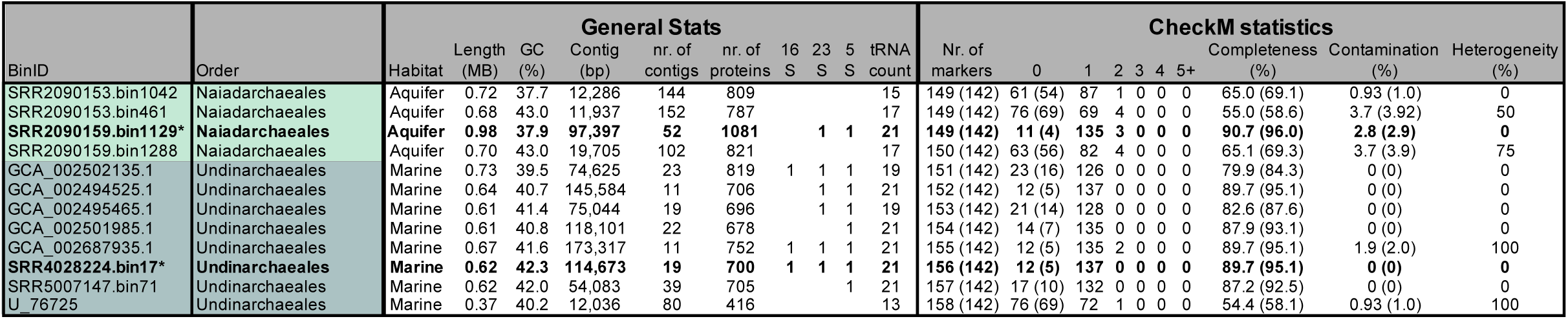
Genome characteristics of Undinarchaeota MAGs. General genome statistics, including genome size, GC content, contig number and other quality characteristics (presence of 16S, 23S and 5S rRNA genes). Additionally, the degree of genome completeness and contamination was estimated using CheckM. The CheckM results were investigated for marker genes commonly absent in DPANN archaea (Supplementary Information) and CheckM was re-run excluding seven marker proteins. The results of this analysis are shown in brackets.

### Undinarchaeota branch between two major DPANN clades

Initial phylogenetic analyses placed Undinarchaeota (formerly UAP2) as a sister lineage to all other DPANN archaea in unrooted trees^13,14,41^. If correct, this placement could give important insights into the timing of DPANN evolution and the nature of the putative last DPANN common ancestor. However, this deep-branching position was poorly supported^13,41^. In order to resolve the phylogenetic relationship of Undinarchaeota and DPANN archaea as well as to test the monophyly of the DPANN radiation, we performed in-depth phylogenetic analyses.

We began by updating the taxon sampling in three published marker protein datasets^10,48,49^ and inferred single-protein trees for each marker to evaluate phylogenetic congruence and detect contaminant sequences and HGTs. Since initial manual inspection revealed extensive incongruence among markers, we developed a marker protein ranking scheme to compare proteins and datasets systematically and without a priori assumptions regarding archaeal phylogeny above the rank of order-, class- or phylum. Briefly, we ranked marker genes according to the extent to which they supported the monophyly of well-established archaeal phylum-, class and order-level lineages (**Supplementary Tables 4-5)** (Methods, Supplementary Information)^6,48,49^. For example, low ranking makers were highly incongruent with individual members of accepted lineages not grouping together and had high numbers of so-called splits indicating HGT events (see Methods for details, **Supplementary Tables 4-5**), while top ranked markers had low indications for potential HGTs between specific DPANN lineages and unrelated taxa. Note that since DPANN monophyly remains actively debated^8,9,11,12,26–29^, we did not penalize marker genes for failing to recover the monophyly of the superphylum as a whole, i.e. the placement of certain DPANN lineages with other archaeal taxa.

These analyses revealed that many universal archaeal genes, including those coding for ribosomal proteins and other core elements of the genetic machinery, may have undergone inter-lineage gene transfers during archaeal diversification (**Supplementary Tables 4-5**). We now have increased power to detect such transfers due to the expanded taxonomic sampling of the archaeal domain compared to previous analyses but the information contained in single protein alignments as well as bootstrap support in corresponding trees is generally low (see Methods, **Supplementary Tables 4-5)**, motivating the use of protein concatenations for phylogenetic reconstructions. However, a large number of gene transfers among markers can mislead phylogenomic analysis because current concatenation and supertree methods assume that all genes evolve on the same underlying tree. To ameliorate the impact of incongruent markers on our inferences, concatenated phylogenies were inferred from the 25% and 50% top-ranked marker proteins which corresponds to those markers with lowest numbers of splits and in turn potential HGTs. We used the best-fitting models of sequence evolution available in both maximum-likelihood using IQ-TREE^50^ and Bayesian frameworks^51^, in combination with alignment recoding, assessment and filtering of compositional biases, as well as statistical topology testing (**Supplementary Table 6, Supplementary Figures 6-56**, Methods, Supplementary Information). Our analyses consistently recovered the clanhood^52^ of the DPANN archaea (including Undinarchaeota) as a whole (that is, all DPANN archaea clustered together on the unrooted tree). Furthermore, our inferences based on curated marker set alignments suggested that Undinarchaeota form a distinct lineage that branches between two other DPANN clans (sequence clusters on the unrooted tree^52^) with maximum statistical support (**Figure 1a,b, Supplementary Figures 6-45, Supplementary Table 6**; Supplementary Information). These clans comprised the Altiarchaeota, Micrarchaeota and Diapherotrites (hereafter referred to as DPANN Cluster 1) and all remaining members of the DPANN (Woesearchaeota, Pacearchaeota, Parvarchaeota, UAP1, Nanoarchaeota, Huberarchaeota, Aenigmarchaeota and Nanohaloarchaeota) (hereafter referred to as DPANN Cluster 2), respectively (**Figure 1a,b**). Finally, in all phylogenies, Undinarchaeota formed two GTDB-level orders, consisting of aquifer-derived and ocean-derived MAGs, i.e. the Naiadarchaeales and Undinarchaeales, respectively (**Figure 1a,b; Supplementary Table 2**).

**Figure 1.**
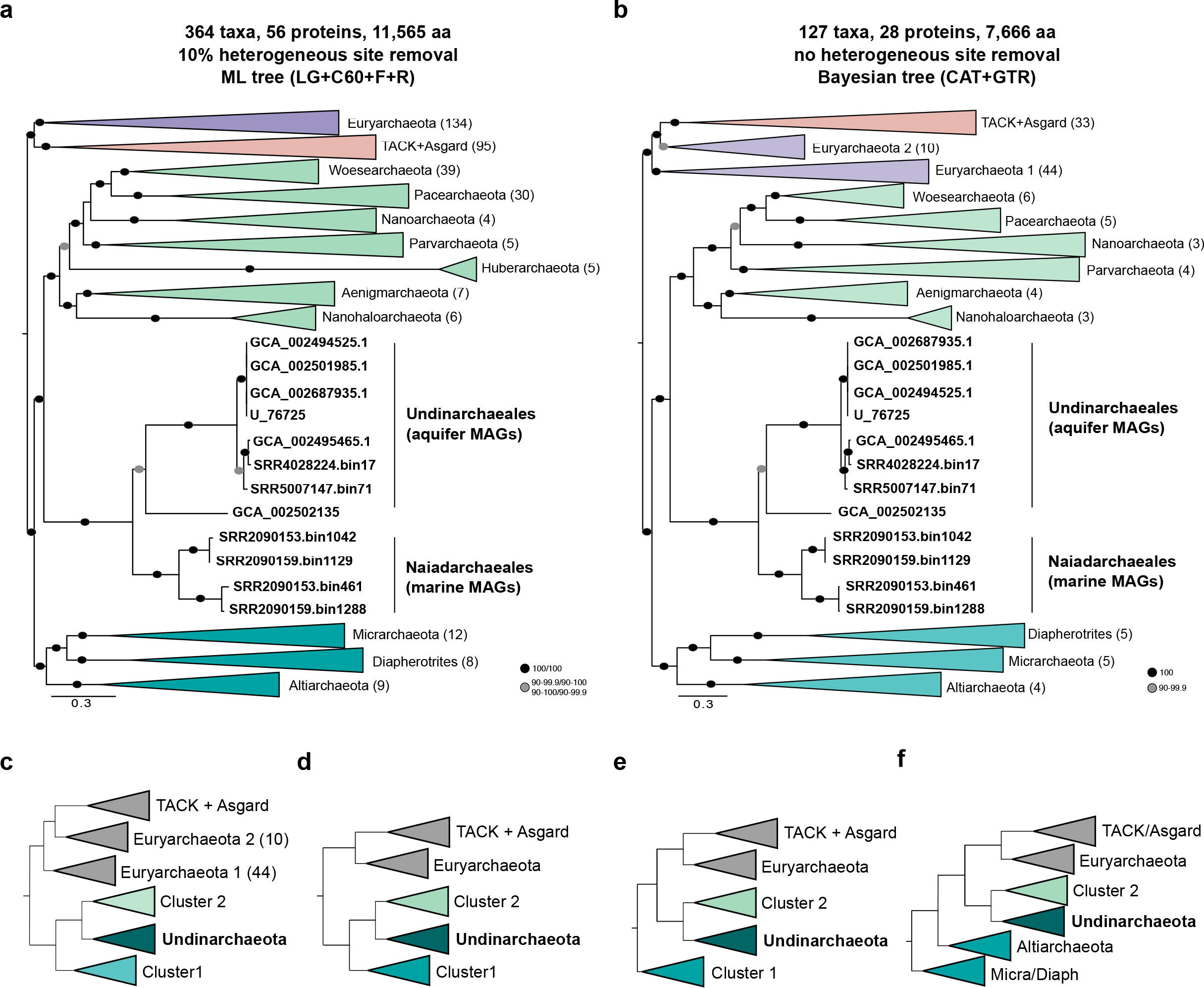
Phylogenetic placement of Undinarchaeota. (a) Maximum likelihood phylogenetic analysis (LG+C60+F+R model) based on an alignment generated with the 50% top ranked marker proteins (n=56) and 364 taxa set. For this alignment 10% of the most heterogeneous sites were removed using chi2 pruning. The full tree is shown in **Supplementary Figure 17**. (b) Bayesian phylogenetic tree (CAT+GTR model) of an alignment generated with the 25% top ranked marker proteins (n=28) and 127 taxa set. For this alignment 40% of the most heterogeneous sites were removed using chi2 pruning. The full tree is shown in **Supplementary Figure 8**. Euryarchaeota 1 includes all Euryarchaeota with the exception of Theionarchaea, Thermococci, Persephonarchaea and Hadesarchaea, which are clustered in Euryarchaeota 2. Scale bar: Average number of substitutions per site. (c-f) Possible positions of the archaeal root inferred using bacteria as an outgroup (c) or using the non-reversible model in IQ-TREE 2 (d-f). All of these methods recover a clade of Undinarchaeota and Cluster 2 DPANN, consistent with synapomorphies including a fused DNA primase and a reduced exosome that lacks Csl4.

Next, we compared these results with phylogenetic inferences based on the 25 and 50% most incongruent markers (**Supplementary Tables 4-5**), which were inferred to have experienced high rates of inter-lineage transfers or were otherwise affected by conflicting phylogenetic signals (**Supplementary Figures 46-49)**. In agreement with our predictions, these analyses yielded phylogenetic trees with various highly supported clusters among unrelated taxa (Supplementary Information). For instance, analyses based on the 25% lowest ranking markers recovered Nanoarchaeota as members of the TACK archaea (**Supplementary Figure 47**) and Nanohaloarchaeota as a sister lineage of Halobacteria (either as a separate cluster (**Supplementary Figure 46**) or with DPANN archaea (**Supplementary Figure 47)**, in agreement with known symbiont-host relationships^16,17,23^. This is particularly notable because we did not a priori penalize trees in which certain DPANN lineages branch with certain other archaeal lineages (**Supplementary Tables 4-5**). In turn, these analyses suggest that conflicting results regarding the placement of certain DPANN lineages, may be due to, at least in part, to the use of a large number of markers affected by host-symbiont HGT. For instance, Nanohaloarchaeota may artificially be drawn towards the Euryarchaeota^28^ when marker sets include too many proteins that were affected by symbiont-host transfers.

Compositional biases in protein sequences can also lead to artefacts in phylogenetic reconstructions^53,54^. To assess the reliability of the inferred placement of Undinarchaeota based on our top-ranked marker protein sets and to ameliorate remaining biases, we subjected the curated alignment to different data treatments including removal of compositionally heterogeneous and fast-evolving sites (see Methods for details, **Supplementary Table 6, Supplementary Figures 13-22 and 30-40**). Removal of compositionally biased sites resulted in notable changes in the tree topology. In particular, the originally inferred sister relationship between Haloarchaea and Methanonatronarchaeia^55^ was supported only in analyses based on the original non-treated alignment; removal of 10% or more of the most biased sites instead supported a close relationship between Methanonatronarchaeia and Archaeoglobales (**Supplementary Figures 13 and 17**), in agreement with more recent work^56^. Importantly, the placement of Undinarchaeota relative to the DPANN Cluster 1 and Cluster 2, as well as the monophyly of each of these clusters, remained stable irrespective of the fraction of heterogeneous sites removed (10% - 40% of sites), suggesting that our inferences are not an artifact of compositional biases.

Finally, we reconstructed phylogenies using a recently-developed non-reversible substitution model that captures asymmetries in the exchange rates between amino acids^57^ to investigate the position of Undinarchaeota relative to the root of the archaeal tree (**Figure 1c, Supplementary Figures 10,11,19,22,24,25,36 and 39, Supplementary Table 6**). This method does not rely on an outgroup and therefore avoids potential LBA artifacts associated with the use of distantly related bacterial sequences to root the archaeal tree^27^. Notably, all our analyses recovered a monophyletic clade of Asgard, TACK and Euryarchaeota with the root being excluded from within this clade with high statistical support (100%). However, the non-reversible model failed to strongly resolve the root position within the DPANN radiation. In particular, the maximum-likelihood root was placed either (a) between all DPANN (including Cluster 1 and 2 as well as Undinarchaeota) on one side and all other Archaea on the other side (**Figure 1c,d, Supplementary Table 6, Supplementary Figures 10-11**), (b) between the Cluster 1 and all other archaea (**Figure 1e, Supplementary Table 6, Supplementary Figures 24-25 and 36**) or(c) between a cluster comprising Micrarchaeota-Diapherotrites and the rest of the Archaea (**Figure 1f, Supplementary Table 6, Supplementary Figures 19, 22 and 39**). However, none of these root positions inferred using non-reversible models received high bootstrap support. Rooting using a bacterial outgroup recovered a root between a monophyletic DPANN clade and the rest of Archaea with moderate to high bootstrap support (94% ultrafast bootstrap^58^, 98% SH-like aLRT support^59^; **Supplementary Figure S56**), consistent with previous results^27^. Thus, our analyses provide strong support for the clanhood of DPANN archaea including Undinarchaeota, but do not confidently resolve the position of the archaeal root either within that clan, or between DPANN and other Archaea^10,14,27^.

### Synapomorphies supporting the monophyly of Undinarchaeota and Cluster 2 DPANN

To further assess the phylogenetic placement of the Undinarchaeota lineage, we surveyed the genomes of DPANN lineages for gene content synapomorphies (shared derived characters) that might enable us to distinguish competing hypotheses for the archaeal root. Similar to other DPANN lineages, Undinarchaeota MAGs encode most proteins involved in replication, transcription and translation and repair (**Supplementary Tables 7-9, Supplementary Figures 58-61**, Supplementary Information). While Undinarchaeota did not share specific features with any of the other archaeal clades, we identified candidate synapomorphies supporting a monophyletic clade comprising Undinarchaeota and Cluster 2 DPANN. Specifically, members of these lineages lack genes encoding the exosome component Csl4, which is present in Cluster 1 DPANN and most other archaea (**Supplementary Figure 61**). The archaeal exosome is thought to consist of four subunits: Rrp41 and Rrp42 form the core ring structure, and Csl4 and Rrp4 constitute the rRNA-binding cap^60^. In spite of the absence of Csl4, Undinarchaeota and Cluster 2 DPANN archaea encode all other subunits of the complex (Rrp4/41/42) suggesting a structural or functional difference of their exosome.

Furthermore, Undinarchaeota and Cluster 2 DPANN share a synapomorphy related to the archaeal DNA primase^61^. Previous work^62^ has suggested that, while DNA primases of most Archaea (including those of the Micrarchaeota) are composed of two subunits encoded by *pri*S and *pri*L, some DPANN lineages (at that time the Nanoarchaeota, Nanohaloarchaeota and Parvarchaeota), were found to possess a *pri*S-*pri*L fusion gene. Our analyses, which includes a larger genomic representation of DPANN archaea, revealed that representatives of the DPANN Cluster 1 consistently encode canonical *pri*S and *pri*L genes, while all Undinarchaeota and DPANN Cluster 2 archaea have a fused version (**Supplementary Table 11**). Note that *pri*S and *pri*L arose from an ancestral duplication and are thus homologous. A phylogenetic analysis of all PriS and PriL subunits (after splitting the fused version), revealed that the fusion likely occurred at the origin of the Undinarchaeota and DPANN Cluster 2 (100/100 and 91.3/99 bootstrap support for PriS and PriL, respectively; **Supplementary Figure 57**).

Consistent with our phylogenetic analyses, these findings support a clade containing Undinarchaeota and DPANN Cluster 2 as sister lineages from which the archaeal root is excluded. It will be interesting to experimentally investigate the functional implication of the identified synapomorphies (exosome component loss, DNA primase fusion) and determine whether they could have played a role in reductive genome evolution in Undinarchaeota and DPANN Cluster 2 archaea.

### Gene repertoires indicate fermentative lifestyle and reliance on a host

#### Catabolism

Comparative genome analyses and inference of the metabolic potential of Undinarchaeota (**Supplementary Tables 7-8 and 12-15**, Supplementary Information for details), suggest that representatives of this clade likely rely on fermentative processes for energy conservation (**Figure 2**). In particular, the presence of the lower Embden-Meyerhof and non-oxidative pentose-phosphate pathway but absence of most genes coding for enzymes of the tricarboxylic acid cycle (TCA) suggest that Undinarchaeota could generate ATP through fermentation of pyruvate to acetate. Simple carbohydrates, such as pyruvate, could perhaps be taken up by passive diffusion^63^. Furthermore, some members of the Naiad- and Undinarchaeales may be able to use extracellular DNA as growth substrates (**Figure 2**). For example, most representatives of the Undinarchaeota encode the complete nucleoside degradation pathway^64–66^(Supplementary Information), including an AMP phosphorylase (DeoA), ribose 1,5-bisphosphate isomerase and Group-III ribulose 1,5-bisphosphate carboxylase (RbcL; RuBisCO) (**Supplementary Figure 59**). In fact, many DPANN representatives have been reported to harbor a RuBisCO homolog and certain members have been suggested to be able to use nucleosides as substrates^66,67^. Undinarchaeota may import DNA via pili encoded by all Undinarchaeota MAGs and subsequently degrade those using their encoded nucleases (Supplementary Information)^68,69^. Intermediates of the nucleoside degradation pathway, such as glycerate-3-phosphate, may subsequently be channeled into the lower glycolytic pathway and contribute to energy conservation by an ATP synthesizing acetate-CoA ligase (acdA). Other products (e.g. glyceraldehyde-3P and fructose-6P produced via gluconeogenesis) may be further metabolized through the non-oxidative pentose-phosphate pathway allowing the synthesis of cellular building blocks such as pyrimidines and purines. It is however notable that group-III-like RuBisCO homologs encoded by Undinarchaeales MAGs have mutations in two positions of the RuBisCO motif^64–67^. In turn, it remains to be determined whether RuBisCO has retained its canonical function in these members of the Undinarchaeales and indeed enables growth on nucleosides (**Supplementary Figure 59**; Supplementary Information). Considering the limited set of predicted proteins involved in central carbon metabolism, experimental verification will be needed to assess whether the encoded pathways provide sufficient ATP to sustain the energy metabolism of the various Undinarchaeota representatives.

**Figure 2.**
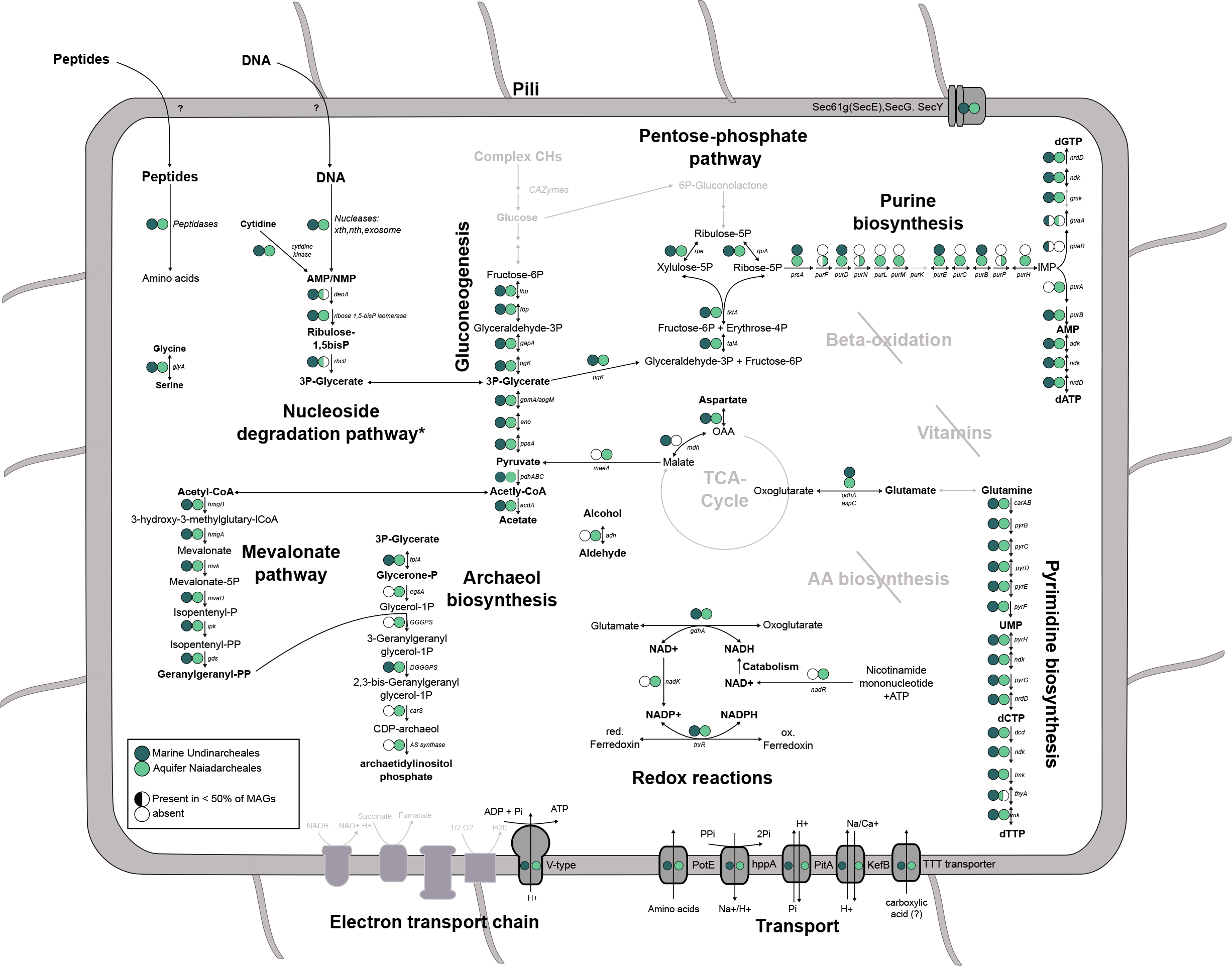
Metabolic characteristics of Undinarchaeota projected on key pathways. Full circles: Gene of interest is present in all or more than 50% of genomes. Half circles: Gene of interest is present in less than 50% but found in at least one Undinarchaeota MAG. Open circle: Gene is absent in all Undinarchaeota MAGs. Dark green: MAGs belonging to the marine Undinarchaeales. Light green: MAGs belonging to the aquifer Naiadarchaeales. \**deo*A was only present in one Naiadarchaeales MAG and genes encoding RuBisCO were only found in two out of four Naiadarchaeales MAGs and it remains to be determined whether this is due to genome incompleteness or a sign of genome streamlining. A detailed list of genes encoded by Undinarchaeota can be found in **Supplementary Tables 7-9**.

#### Anabolism

Even though all representatives of the Undinarchaeota encode a near complete gluconeogenesis pathway (**Figure 2**) including the potentially ancient bifunctional fructose 1,6-bisphosphate (FBP) aldolase/phosphatase, which would allow the synthesis of fructose-6-phosphate^70^, many other biosynthesis pathways are incomplete. For instance, while the Naiadarchaeales MAGs encode all proteins required to synthesize archaeal-type ether lipids, lipid biosynthesis pathways are incomplete in Undinarchaeales MAGs, which lack key genes for the conversion of glycerone-phosphate to archaetidylinosytol-phosphate, in spite of the presence of the archaeal mevalonate pathway^10^ in representatives of both lineages. Incomplete pathways for lipid biosynthesis are particularly common in DPANN Cluster 2 representatives (incl. *N. equitans*) (**Figure 3, Supplementary Figure 62**)^10,13,14,71,72^ and the characterization of the *N. equitans* - *I. hospitalis* symbiotic system has confirmed the exchange of lipids between symbiont and host^73^. Thus, while members of the Naiadarchaeales may synthesize their own lipids, Undinarchaeales representatives may depend on an external source of archaeal or bacterial lipids or intermediates in spite of the presence of the mevalonate pathway and their ability to synthesize geranylgeranyl diphosphate (Supplementary Information). Similarly, the lack of several genes coding for enzymes of the purine biosynthesis pathways in members of the Undinarchaeales but not Naiadarchaeales, indicates that the former are also dependent on the external source of inosine monophosphate (IMP) or other intermediates of the purine biosynthesis pathway (**Figures 2-3, Supplementary Tables 7-8 and 9**, Supplementary Information).

**Figure 3.**
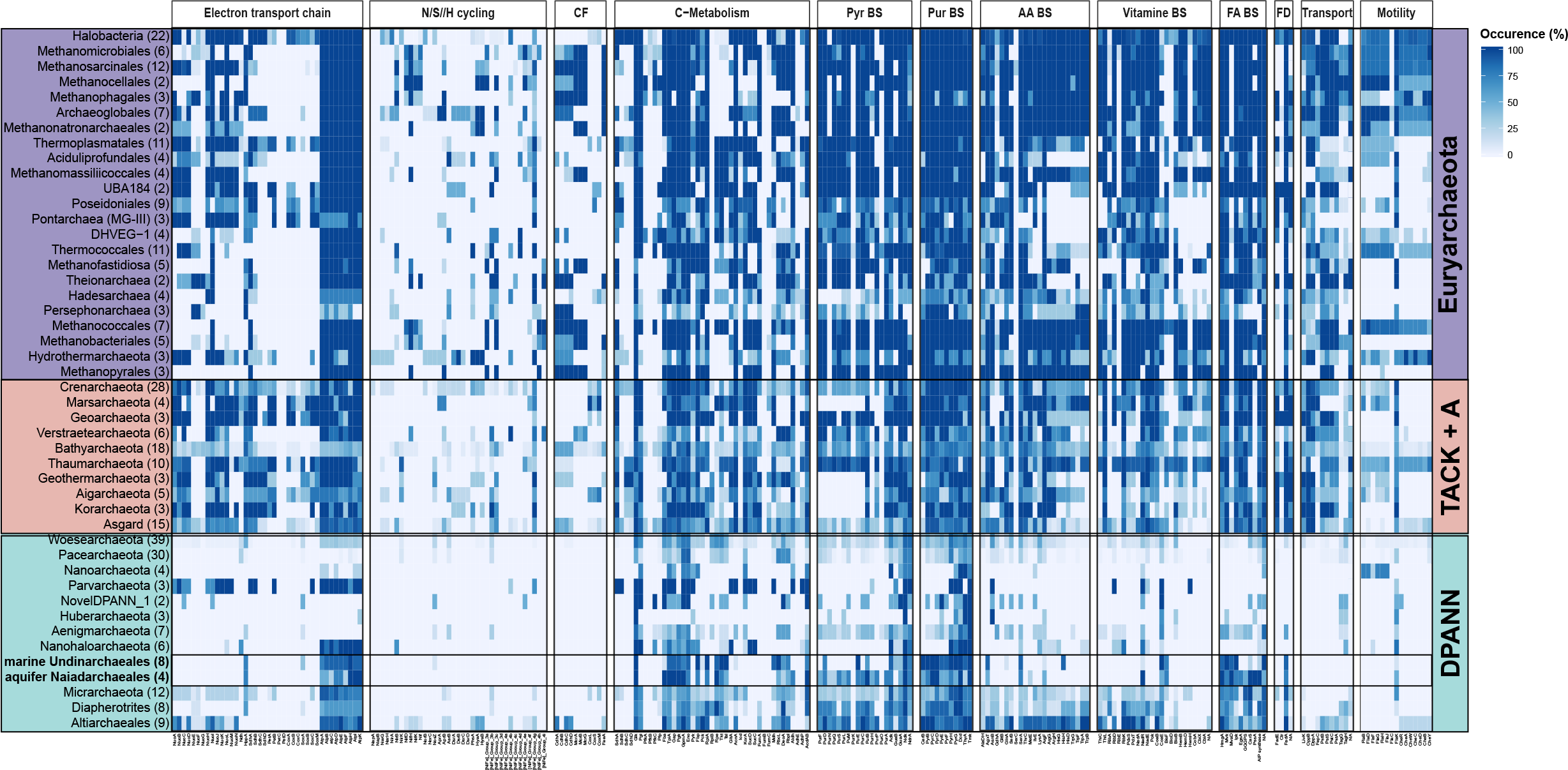
Occurrence of key metabolic proteins across major archaeal lineages. Protein occurrence was calculated by detecting key proteins of interest across 364 archaeal genomes and calculating the occurrence in percent across the total number of genomes included in each phylogenetic cluster based on a presence/absence table. Number in brackets: number of genomes included in each individual phylogenetic cluster. CF = Carbon fixation. FD = Fatty acid degradation. A table listing gene occurrences in archaeal lineages can be found in **Supplementary Table 9** and a list of metabolic genes shown in this plot can be found in **Supplementary Table 19**.

Furthermore, all Undinarchaeota representatives lack genes encoding proteins involved in amino acid and vitamin biosynthesis. Notably, and in agreement with several other potentially symbiotic DPANN archaea^13,14^, all Undinarchaeota representatives seem to contain a limited set of genes for transporters such as amino acid transporter PotE and uncharacterized di/tri-carboxylate transporters (Supplementary Information, **Figures 2-3**), suggesting that they are unable to acquire all essential building blocks directly from the environment. In turn, members of the Undinarchaeota seem to depend on partner organisms to provide compounds that cannot be synthesized or taken up from the environment using transporters. Key differences among the biosynthetic capabilities of members of the Naiad- and Undinarchaeales, may translate into varying substrate requirements and demands from potential host organisms.

#### Cell-cell interactions

Consistent with a host-dependent lifestyle, we detected several proteins with domains known to be involved in cell-cell interactions and common among symbionts^13^. While Undinarchaeota lack genes for ankyrin domain proteins and only encode a small number of beta propeller/WD40 domain proteins, the proteome of members of the Naiadarchaeales comprises diverse proteins with immunoglobulin domains, while Undinarchaeales encode Concanavalins/LamG domain proteins (arCOG07813) (**Supplementary Tables 20-21**). Homology modeling and structure predictions suggested that these proteins might encode potential cell adhesion proteins (**Supplementary Tables 22-23**) and in turn may be involved in attachment or biofilm formation in Undinarchaeota. Notably, the complete absence of LamG domain proteins in the Naiadarchaeales representatives indicates that members of the two Undinarchaeota orders rely on different mechanisms mediating potential symbiont-host interactions.

#### Gene repertoires and reductive genome evolution in DPANN

The presence and absence patterns of genes involved in core metabolic pathways of Undinarchaeota MAGs shows similar trends as seen in DPANN Cluster 2 archaea further supporting the sisterhood of this clade (**Figure 4, Supplementary Tables 7-9,12**). For instance, most DPANN Cluster 2 archaea lack genes involved in core metabolic pathways, such as the electron transport chain, carbon fixation other than the RuBisCO gene, as well as transport and motility genes (**Figure 3**)^10,13–16,23,25,39,74^. While Undinarchaeota seem to have more complete pathways than many of the DPANN Cluster 2 representatives, they appear metabolically less flexible than several members of DPANN Cluster 1 (**Figure 3**)^30,31,75,76^. For instance, members of the DPANN Cluster 1 have more complete nucleoside and lipid biosynthesis pathways and include free-living organisms. In particular, representatives of the Altiarchaeota have been suggested to comprise autotrophic archaea that may use the Wood-Ljungdahl pathway for carbon fixation^24,25,30^ and while this lineage includes symbionts, these do not seem to be obligate^77,78^. In fact, Altiarchaeota have recently been found to include members that likely serve as hosts for Huberarchaeota belonging to DPANN Cluster 2^25^. Furthermore, at least some members of the Diapherotrites have been suggested to be capable of a fermentative free-living lifestyle^76^. However, in spite of overall gene repertoire patterns being consistent with results from our phylogenetic analyses, there is a large variation in gene content and extent of genome reduction within DPANN lineages^13^. Thus, our analyses further support the notion that while reductive genome evolution may have characterized the evolution of Undinarchaeota and DPANN Cluster 2 archaea already at the time of their divergence, the extent of streamlining varies widely and seems to have occurred in parallel in different lineages^13^.

**Figure 4.**
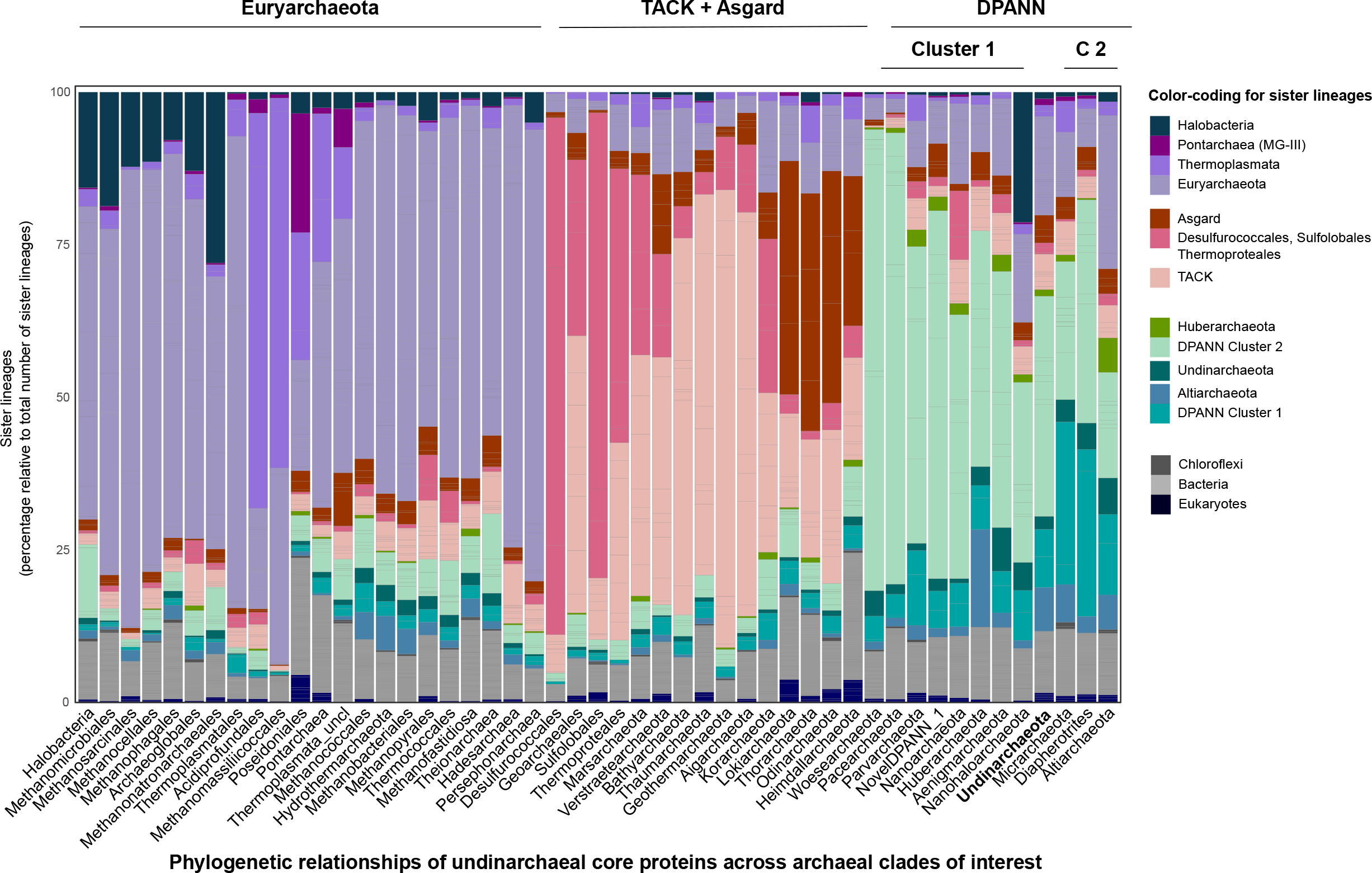
Potential HGTs of core proteins of Undinarchaeota across archaeal lineages. The plot is based on phylogenetic analyses of 520 core proteins (shared across at least three MAGs of the Undinarchaeota), which included a representative set of archaeal, bacterial and eukaryotic genomes (n=3482 taxa). In particular, this plot depicts sisterhood-relationships observed in these phylogenies for all major archaeal lineages including the Undinarchaeota (see **Supplementary Table 18** for the full list of genomes used and **Supplementary Table 16-17** for the accompanying data for the HGT analysis). See custom scripts and Methods for more details.

### Insights into interaction partners of Undinarchaeota

Genomic analyses of the first members of the Nanoarchaeota^39^ as well as our marker protein analyses have indicated that DPANN symbionts may have exchanged genes with their hosts. Furthermore, co-occurrence patterns have recently allowed to pinpoint Altiarchaeota as host for the Huberarchaeota^24,25^. Thus, to shed light onto potential interaction partners of the Naiad- and Undinarchaeales, respectively, we have inferred routes of horizontal gene transfer and generated proportionality networks.

In particular, we reconstructed phylogenies of proteins present in at least three Undinarchaeota genomes (520 genes total) and analyzed sisterhood-relationships among taxonomically distinct lineages including a reference set of 364 archaeal, 3020 bacterial and 100 eukaryotic genomes (**Figure 4, Supplementary Tables 16-17**, Methods, Supplementary Information). Using this approach, we were able to successfully detect signals of HGT among known host-symbiont pairs. For instance, our analysis indicates that Nanoarchaeota and Huberarchaeota have exchanged 11% and 16% of the analyzed proteins with their respective hosts belonging to the Crenarchaeota and Altiarchaeota, respectively. Furthermore, our phylogenies suggest that Nanohaloarchaeota and Halobacteria shared a large amount of analyzed proteins horizontally (12.4% and 7%) (**Figure 4**), which is in agreement with our core gene analyses (**Supplementary Tables 4-5**). However, some of the sisterhood relationships could also be due to compositional biases as a result of protein sequence adaptation to high salinity environments or phylogenetic artifacts. Notably, Undinarchaeota does not show a dominant fraction of genes shared with a specific lineage, i.e. most proteins seem to cluster with taxonomically closely related DPANN lineages (**Figure 4**) or in some cases with potentially deeply branching Euryarchaeota such as Hydrothermarchaeota or group 1 methanogens (**Supplementary Tables 4,16,17**). The few Undinarchaeota genes that may have been shared with distantly related taxa encoded proteins that grouped for instance with Bathyarchaeota and Thermoplasmata (i.e. including Poseidoniales and Pontarchaea, or MG-II and -III) (Supplementary Information, **Supplementary Tables 4 and 16**). For instance, the manual inspection of respective trees revealed that certain Undinarchaeota, in particular marine Undinarchaeales, may have exchanged core genes with members of the Pontarchaea, including a gene encoding ribosomal proteins S19 (arCOG04099, TIGR01025), respectively (**Supplementary Table 4**). Yet, due to the small fraction of putative HGTs with any of these non-DPANN lineages relative to the total proteome of Undinarchaeota, it remains to be assessed whether these patterns are genuine or result from phylogenetic artifacts, lack of partner organisms in current databases or Undinarchaeota forming less intimate interactions than DPANN Cluster 2 archaea.

Finally, we used a read-based co-occurrence analysis to assess whether MAGs of Undinarchaeota are proportional to other archaeal and bacterial genomes^79^ (see Methods for details). Due to the low number of available samples for our inferences of co-proportionality, this analysis did not detect any significant co-occurrence patterns for members of the Naiadarchaeales. However, it did indicate that marine Undinarchaeales may co-vary with three genomes of the Chloroflexi, all belonging to the order Dehalococcoidales (**Supplementary Figure 63**, Supplementary Information).

In turn, further analyses, such as fluorescence *in situ* hybridization, will be needed to identify the organisms engaging in symbiotic interactions with Undinarchaeota. For instance, it will be important to determine their host-specificity and test whether certain members of marine Undinarchaeales indeed interact with members of the Chloroflexi or Pontarchaea. Together with previous findings^80^, our results suggest that certain DPANN symbionts could have non-archaeal partners highlighting the importance of taking into account total community composition when inferring potential co-occurrences between DPANN and other organismal groups.

## Conclusions

We provided an updated view on the evolution of DPANN archaea by investigating the phylogenetic placement and metabolic potential of the Undinarchaeota, a previously uncharacterized archaeal lineage whose members have small genomes and limited metabolic gene repertoires indicative of various auxotrophies. Its members, classified into the two order-level lineages Undinarchaeales and Naiadarchaeales, may lead a symbiotic lifestyle and depend on (a) partner organism(s) for growth and survival similar to other members of the various DPANN representatives^10,13,14^. In addition to insights into the biology of members of the Undinarchaeota, their genomes contributed valuable data to address the longstanding debate about whether DPANN indeed form a monophyletic clade or instead represent an artifactual assemblage of phylogenetically disparate lineages^11,12,14,26–31,81,82^. In our analyses, which were based on a broader taxon sampling, we not only attempted to account for the effect of compositional biases but also developed a new method for detecting potential HGTs of marker proteins. Our results, based on markers least effected by HGT, consistently recovered clan-hood of all DPANN lineages, such that DPANN monophyly (i.e. monophyly of DPANN Cluster 1, Undinarchaeota and DPANN Cluster 2) solely depends on the placement of the root. Notably, in analyses based on phylogenetic markers that had indications for extensive inter-lineage HGT, Nanoarchaeota and Nanohaloarchaeota group with their respective hosts the TACK-archaea and Halobacteria, respectively. This finding indicates that the previous clading of certain DPANN lineages with Euryarchaeota^28,81^ might to a certain degree be a result of conflicting signals among marker proteins concatenated into one supermatrix. Finally, our analyses robustly excluded the root from a clade consisting of Euryarchaeota, TACK and Asgard lineages, strengthening the hypothesis that some or all of the DPANN superphylum diverged early from the archaeal tree^27^. In turn, further exploration of DPANN promises to shed light on their roles in biogeochemical cycles and food webs as well as the early evolution of cellular life and symbiosis.

## Species description

*Description of Candidatus* Undinarchaeum (Un.din.ar.chae’um. L. n. *Undina* female water spirit or nymph (from L. fem. n. *unda* water, wave); N.L. neut. n. *archaeum* (from Gr. adj. *archaios* ancient) archaeon; N.L. neut. n. *Undinarchaeum* an archaeon of water origin). *Candidatus* Undinarchaeum marinum (ma.ri’num. L. neut. adj. *marinum* of the sea, marine)

*Description of Candidatus* Undinarchaeum marinum (ma.ri’num. L. neut. adj. *marinum* of the sea, marine). Type material is the genome designated as SRR2090159.bin1129 representing *Candidatus* Undinarchaeum marinum.

*Description of Candidatus* Naiadarchaeum (Na.iad.ar.cha’eum. L. fem. n. *Naias*, -*adis* a water-nymph of springs and streams, Naiad from Greek mythology; N.L. neut. n. *archaeum* (from Gr. adj. *archaios* ancient) archaeon; N.L. neut. n. *Naiadarchaeum* an archaeon from the freshwater).

*Description of Candidatus* Naiadarchaeum limnaeum (lim.nae’um. N. L. neut. adj. *limnaeum* (from Gr. adj. *limnaios* from the marsh, lake) living in the freshwater). Type material is the genome designated as SRR4028224.bin17 representing *Candidatus* Naiadarchaeum limnaeum.

Based on these genera, we further propose the families *Candidatus* Undinarchaeaceae fam. nov. and *Candidatus* Naiadarchaeaceae fam. nov., the orders *Candidatus* Undinarchaeales ord. nov. and *Candidatus* Naiadarchaeales ord. nov., the class *Candidatus Undinarchaeia* class nov., and the phylum *Candidatus* Undinarchaeota phylum nov.

## Methods

### Genome recovery and reconstructions

#### Reconstructing MAGs of Undinarchaeota

We mined the Sequence Read Archive (SRA) for public metagenomes using SingleM (https://github.com/wwood/singlem). Thereby, we found 37 metagenomes containing reads assigned to Undinarchaeota (**Supplementary Table 1**). Genomes from these 37 metagenomes were recovered by assembly with megahit (v1.1.3)^83^, mapping raw reads from each sample to its corresponding assembly using BamM (v1.7.3) (https://github.com/Ecogenomics/BamM) using BWA ‘mem’ (v0.7.15)^84^ with default parameters. Binning was carried out with MetaBAT (v2.11.1)^85^ using default parameters and allowed the reconstruction of six MAGs. The quality of these six as well as the six previously published Undinarchaeota MAGs was assessed using CheckM (v1.0.7)^86^. Additionally, the CheckM marker set was screened for markers that were absent in all 12 Undinarchaeota MAGs. Thereby, we identified 7 markers (PF01287, PF01667, PF01912, PF01982, PF04127, TIGR00432 and TIGR01213) that were not only absent in Undinarchaeota but often in other DPANN archaea (Supplementary Information). As the inclusion of these markers might underestimate the completeness scores, we also ran CheckM excluding these markers. Both results are summarized in **Table 1**.

#### Contamination Screening

Contigs from all 12 Undinarchaeota MAGs were manually investigated for signs of contamination by looking for an abnormal GC-content (∼10% difference in average GC-content) and/or taxonomic composition. To determine the taxonomic composition, the protein files of each genome (see section below) were searched against the NCBI-non-redundant (nr) database (downloaded in November 2018) using DIAMOND v0.9.22.123 (settings: --more-sensitive --evalue 1e-10 --seq 100 --no-self-hits --taxonmap prot.accession2taxid.gz)^87^. The best hit for each protein was selected based on the highest e-value and bit-score. Subsequently, the obtained NCBI taxonomy identifier was merged with the taxonomic string information provided by NCBI using a mapping file generated in January 2019 using ncbitax2lin (https://github.com/zyxue/ncbitax2lin). Contigs with an “abnormal” taxonomic distribution (>90% of proteins on a contig assigned to a non-archaeal or non-DPANN lineage with a percentage identity above 80%) were either removed or confirmed by phylogenetic analyses of single gene trees (see Supplementary Information).

Notably, several Undinarchaeota MAGs appeared to encode a short genomic region similar to an archaeal fosmid sequence deposited at NCBI and referred to as uncultured marine group I/III euryarchaeote KM3_51_D01 (https://www.ncbi.nlm.nih.gov/nuccore/663520902). In particular, proteins encoded in this region shared about 84% ANI to protein homologs found on this fosmid. To investigate whether our MAGs may be contaminated by contigs of members of the MGII/III archaea (Pontarchaea/Poseidoniales), we performed single protein trees of marker proteins located in this genomic region and queried the ORFs located on fosmid KM3_51_D01 against NCBI-nr Importantly, our results (i) suggested that this fosmid does not belong to the Pontarchaea/Poseidoniales archaea and should be reclassified as “uncultured archaeon KM3” and (ii) verified, that respective genomic regions in our MAGs are not contaminated. Please see Supplementary Information for further details.

### Phylogenetic analyses

#### Generation of a backbone datasets for phylogenetic analyses

##### Archaeal backbones

To generate a representative archaeal reference set all archaeal genomes were downloaded from NCBI (January 2019). Genomes with a completeness >40% and contamination <25% (determined with CheckM lineage_wf v1.0.11)^86^ were used for an initial phylogenetic analysis using the PhyloSift marker set. In brief, homologs of 34 PhyloSift marker proteins were identified in each genome using the PhyloSift v1.0.1 ‘find’ mode (settings: --besthit)^48^. Subsequently, marker proteins were individually aligned using MAFFT L-INS-i v7.407 (settings: --reorder)^88^ and trimmed using BMGE v1.12 (settings: -m BLOSUM30 -h 0.55)^89^. The concatenated protein alignment was used to reconstruct a phylogenetic tree using IQ-TREE v1.6.7 (settings: -m LG -nt AUTO -bb 1000 -alrt 1000)^50,58,59^. Based on this tree, a sub-selection of 364 archaeal reference genomes was generated (comprising 352 reference archaeal and 12 Undinarchaeota genomes, **Supplementary Table 18**), which covers the archaeal phylogenetic diversity while still allows to perform computationally intensive phylogenetic analyses needed to resolve the placement of DPANN in the tree of Archaea. Selected genomes preferentially derived from type strains as well as from known hosts of DPANN archaea. Furthermore, representative MAGs and SAGs were selected based on high completeness and low degree of contamination. Based on an initial maximum likelihood analysis (IQ-TREE, C60+LG model) including these 364 species, *Zestosphaera tikiterensis* NZ3 seems to represent a lineage of the order Sulfolobales rather than the Desulfurococcales^90^ with high statistical support (i.e. bootstrap of >97%, **Supplementary Figure 9**). Furthermore, the placement of “Acidilobales” within the Desulfurococales was highly supported (i.e. bootstrap of 100%, **Supplementary Figure 9**), which indicates that Acidilobaceae represents a family level lineage within the Desulfurococcales rather than an order level lineage^91^. This also corresponds to the taxonomy of the Genome Taxonomy Database (GTDB; https://gtdb.ecogenomic.org). We therefore assigned both *Zestosphaera tikiterensis* NZ3 and Acidilobaceae to the Desulfurococcales in our analyses.

To create a smaller archaeal reference set suitable for Bayesian phylogenies, the 364 archaeal genome set was further subsampled to only include 127 taxa. In particular, at least two members of all major archaeal order-level lineages were selected based on phylogenetic distance, genome quality. For MAGs that were only classified at phylum level (i.e. Woesearchaeota), we selected a representative set of taxa based on the 364 species tree.

##### Bacterial backbones

To generate a bacterial backbone, we downloaded a list of available genomes from NCBI in 2018 (ftp://ftp.ncbi.nlm.nih.gov/genomes/genbank/bacteria/). This list was parsed to only include type strains or representative genomes. From this list we randomly selected a representative strain per genus. In a second step, we selected 15-30 genomes from candidate phyla from the unparsed list. This initial set was screened for completeness and contamination in CheckM^86^ and genomes with less than 50% completeness and more than 10% contamination were excluded. We performed initial phylogenies using alignments generated by PhyloSift^48^ using the PhyloSift search, align and name algorithms to find marker proteins. Individual genes were aligned with MAFFT L-INS-i (settings:--reorder)^88^ and trimmed using BMGE (settings: -m BLOSUM30 -h 0.55)^89^. The concatenated protein alignment was used to reconstruct a phylogenetic tree using IQ-TREE v1.6.7 (settings: -m LG -bb 1000 - alrt 1000)^50^. The initial alignments were screened for long-branches and genomes that did not fall into the expected position in the tree were removed from the taxa selection resulting in a set of 3020 bacterial genomes.

##### Eukaryotic backbones

98 eukaryotic genomes and largely complete transcriptomes were selected so as to obtain even sampling across the known major lineages of eukaryotes. Where possible, we selected free-living representatives of clades with a completely sequenced genome; alternatively, we selected the most complete transcriptome from a free-living member of the group.

#### Selection of marker proteins for phylogenetic analyses

To accurately place Undinarchaeota in the archaeal tree of life, we identified suitable markers among three different but overlapping marker proteins sets: (a) a set of herein identified potential single-copy, near universal archaeal orthologues and the marker protein sets implemented in GTDB (b) and PhyloSift (c).

(a) We inferred a set of markers by running the OMA standalone algorithm^92^ on a set of 149 archaeal genomes chosen to represent known taxonomic diversity. We inferred single protein trees for 97 orthologues that were present in at least 75% of the genomes and manually inspected the resulting trees to identify and remove 15 markers that had undergone gene transfer, resulting in a final set of 82 candidate single-copy orthologs.

(b) The GTDB marker set was downloaded February 2019 (release r86 downloaded from https://gtdb.ecogenomic.org). This set was modified by combining the 122 archaeal GTDB marker set with the complete TIGRFAM database. Therefore, the extra PFAMs defined in the GTDB marker set were added to the TIGRFAM database (TIGRFAMs_15.0_HMM.LIB: downloaded Sept 2018) and the combined database will be referred to as TIGRFAM_GTDB.

(c) The PhyloSift marker set^48^ included all previously reported markers excluding genes for the phenylalanine synthetase. These three marker sets were combined by adding the PhyloSift and OG marker protein set to the TIGRFAM_GTDB database with the final database consisting of 151 non-redundant protein families.

#### Assessment of the 151 marker set for suitability in concatenations

Homologs in the bacterial, eukaryotic and archaeal backbone datasets as well as in Undinarchaeota MAGs were identified using hmmsearch v3.1b2 (settings: -E 1e-20)^93^. The output was parsed to only include the best hit based on the e-value and bit-score. Afterwards, marker proteins were individually aligned using MAFFT L-INS-i (settings: --reorder)^88^ and trimmed using BMGE (settings: -t AA - m BLOSUM30 -h 0.55)^89^. Subsequently, we inferred single protein phylogenies with IQ-tree (settings: -m LG+G+F+R -bb 1000 -alrt 1000)^50^ for each of these 151 marker protein families, including homologs identified in the corresponding reference genomes (i.e. 364 archaea, 3020 bacteria and 98 eukaryotes (see details above). Marker proteins in which archaea and eukaryotes were not monophyletic (i.e. archaeal lineages were paraphyletic with some lineages emerging from within Bacteria), were excluded from any further analysis (see also **Supplementary Tables 4-5**).

Next, we used a custom python script (count_sister_taxa.py; https://github.com/Tancata/phylo/blob/master/count_sister_taxa.py) to rank each marker protein phylogeny based on the extent to which it resolved monophyletic clades of well-established archaeal phylum- or order-level lineages. In brief, the script counts for each tree, how many times a certain taxon does not group with its expected taxonomic clade across all bootstrap trees, thus representing so-called splits. Splits (which may arise through HGT or phylogenetic artifacts) are counted for each taxon across all protein phylogenies and provide the basis for ranking the different markers according to their reliability. Here, we counted both the total number of splits as well as the splits normalized to the total species count. Note, monophyly was assessed on the level of following archaeal clades: Geothermarchaeota, Halobacteria, Methanonatronarchaeales, Methanomicrobiales, Methanosarcinales, Methanocellales, Methanophagales, Archaeoglobales, Thermoplasmatales, Acidiprofundales, Methanomassiliicoccales, Poseidoniales, Thermoplasmata (not assigned at order level), Pontarchaea, Undinarchaeota, Woesearchaeota, Pacearchaeota, NovelDPANN_1 (UAP1), Parvarchaeota, Nanohaloarchaeota, Aenigmarchaeota, Diapherotrites, Huberarchaeota, Micrarchaeota, Altiarchaeota, Methanopyrales, Methanobacteriales, Methanococcales, Desulfurococcales, Sulfolobales, Thermoproteales, Marsarchaeota, Thermococcales, Theionarchaea, Methanofastidiosa, Hadesarchaea, Persephonarchaea, Odinarchaeota, Verstraetearchaeota, Thorarchaeota, Lokiarchaeota, Heimdallarchaeota, Bathyarchaeota, Thaumarchaeota, Korarchaeota, Aigarchaeota, Geoarchaeales, Hydrothermarchaeota and Nanoarchaeota. The number of split phylogenetic clusters (in percentage) as well as the total number of splits normalized by the total number of species within each tree was used as criteria to rank and define the 25%, 50% and 75% top as well as 25% and 50% lowest ranking marker proteins (**Supplementary Tables 4-5**). Finally, the 50% top-ranking single protein trees were manually inspected for signs of contamination or paralogues that were then manually removed from the marker protein sequences. After manual cleaning, the marker proteins were aligned using MAFFT L-INS-i^88^ and trimmed using BMGE^89^ as described above. The single proteins were concatenated using catfasta2phyml.pl (https://github.com/nylander/catfasta2phyml). Note that all single protein phylogenies as well as the individual protein files and alignments have been deposited in a Zenodo repository (10.5281/zenodo.3672835).

#### Phylogenetic analyses for the species trees

To confirm the placement of Undinarchaeota several different phylogenetic trees were generated (all trees are summarized in **Supplementary Table 6** and shown in full in **Supplementary Figures 6-56**):

##### (I) Using different subsets of marker proteins for maximum-likelihood phylogenies

The alignment for the concatenated 25%, 50% and 75% top as well as 25% and 50% lowest ranking marker proteins were generated for the 364 as well as 127 taxa set and phylogenetic trees were generated using IQ-TREE (v1.6.7, settings: -m LG+C60+F+R -bb 1000 -alrt 1000)^50^.

##### (II) Stationary trimming for maximum-likelihood phylogenies

The concatenated alignment for the 50% top ranking marker proteins was aligned with MAFFT L-INS-i^88^ and trimmed with BMGE^89^ using stationary trimming (options: -s FAST -h 0:1 -g 1) to remove compositional heterogenous sites from both the 127 and 364 taxa set. Trees were generated using IQ-TREE^50^(settings: -m LG+C60+F+R -bb 1000 -alrt 1000).

##### (III) Data recoding for maximum-likelihood phylogenies

The concatenated alignment for the 50% top ranking marker proteins (alignment: MAFFT L-INS-i^4^, trimming: BMGE^89^) was re-coded into 4 character-states (SR4 re-coding^94^; i.e. data simplification from 20 to four character states) to reduce compositional heterogeneity both for the 127 and 364 taxa set using a custom script (Recode_aa.py). The states used were the following: A,G,N,P,S,T = A; C,H,W,Y = C; D,E,K,Q,R = G and F,I,L,M,V = T. Phylogenetic trees were generated using IQ-TREE (settings: -m C60SR4 - bb 1000 -alrt 1000).

##### (IV) Removing fast-evolving sites for maximum-likelihood phylogenies

SlowFaster^95^ was used on the concatenated alignment for the 50% top ranking marker proteins (alignment: MAFFT L-INS-i^88^, trimming: BMGE^89^; settings see above) to remove fast-evolving sites both from the 127 and 364 taxa set. Sites were removed in a stepwise manner removing 10%, 20%, 30% and 40% of the fastest evolving sites. Phylogenetic trees were generated using IQ-TREE^50^ (settings: -m LG+C60+F+R -bb 1000 -alrt 1000).

##### (V) Removing heterogenous sites for maximum-likelihood phylogenies

Alignment_pruner.pl (https://github.com/novigit/davinciCode/blob/master/perl) was used to remove heterogenous sites in a stepwise manner from the concatenated alignment for the 50% top ranking marker proteins (alignment: MAFFT L-INS-i^88^, trimming: BMGE^89^; settings see above) both from the 127 and 364 taxa set. Sites were removed in a stepwise manner removing 10%, 20%, 30% and 40% of the most heterogeneous sites and the trees were generated using IQ-TREE^50^ (settings: -m LG+C60+F+R - bb 1000 -alrt 1000).

##### (VI) Using published marker gene sets for maximum-likelihood phylogenies

Several alternative marker sets were used to confirm the tree topologies. These include the PhyloSift marker set^48^ using the default alignment with hmmalign^96^ or MAFFT L-INS-i^88^, the 122 archaeal GTDB marker set^49^ using FastTree v2.1.10^97^(settings: WAG, LG) or IQ-TREE^50^ (settings: LG+C60+F+R), the RP14 marker set^10^ and the universal 48 marker set^6^. Please note that the PhyloSift^48^, 122 GTDB marker gene set^49^ and RP14 marker protein^10^ set were subjected to phylogenetic analyses before defining the 364 taxa set and include slightly less taxa (356).

##### (VII) Using different subsets of marker proteins for Bayesian phylogenies

The concatenated alignment for the 25% and 50% top ranking marker proteins (alignment: MAFFT L-INS-i^88^, trimming: BMGE^89^ and the 127 taxa set were used for Bayesian inferences using PhyloBayes-MPI v1.8^51^ (settings: -cat -gtr -x 10 -1 -dgam 4). In particular, for each marker protein family, four parallel chains were run until convergence was reached, unless stated otherwise (maxdiff < 0.3; settings: bpcomp -x 25%_burnin chain1 chain2 chain3 chain4). Additionally, we checked for the minimum effective size using tracecomp (minimum effective size > 50; settings: -x 25%_burnin chain1 chain2 chain3 chain4). Subsequently, readpb_mpi of the pb_mpi package was used for posterior predictive testing (settings: -- allpred).

##### (VIII) Rooting maximum-likelihood phylogenetic trees using non-reversible models

To root our phylogenetic trees, we analyzed the concatenated alignment based on the 50% top ranking marker proteins using a single partition model and a multiple partition non-reversible model. These analyses were performed on the 127 and 364 taxa set using the full alignments and the recoded alignments in which 20% or 40% of the most heterogeneous sites were removed. Trees were generated with an updated version of IQ-TREE v2^57^ (while all previously mentioned phylogenies were generated with IQ-TREE v1)^50^. The single partition models were run with the following settings: -mset LG -bb 1000 -alrt 1000 -m MFP+MERGE, followed by --model-joint NONREV --min-freq 0.001 -nparam 10 -optfromgiven - bb 1000 -alrt 1000. The multiple partition model was run with these settings: -model-joint NONREV --min-freq 0.001 -nparam 10 -optfromgiven -bb 1000 -alrt 1000. Additionally, we used minimal ancestor deviation (MAD) rooting to determine the root position^98^.

##### (IX) Rooting maximum-likelihood phylogenetic trees using bacteria as an outgroup

To root our phylogenetic trees with bacteria as an outgroup we added 88 bacterial genomes to our 127 archaeal taxa selection. These 88 bacterial genomes were selected from the bacterial backbone tree by covering a broad diversity and including at least one to three genomes per phylum. However, to minimize LBA artifacts, we did not select members of the CPR^99^ as well as other symbionts or thermophiles, which often emerge on long branches in phylogenetic trees^100,101^. Next, we identified homologs of the universal 48 markers^6^ in each genome, individually aligned proteins using MAFFT L-INS-i^88^ (settings: --reorder) and trimmed using BMGE (settings: -t AA -m BLOSUM30 -h 0.55)^89^. Single proteins were concatenated using catfasta2phyml.pl (https://github.com/nylander/catfasta2phyml) and a phylogenetic tree was generated using IQ-TREE (settings: -m LG+C60+F+R -bb 1000 -alrt 1000).

### Phylogenetic analyses of the core Undinarchaeota protein set

A set of 520 proteins for Undinarchaeota were selected based on arCOGs detected in at least three or more Undinarchaeota genomes with an e-value greater than 1e-20 (see below). Next, these 520 arCOGs were queried against a protein database of 364 archaeal, 3020 bacterial and 98 eukaryotic genomes using PSI-BLAST (settings: -evalue 1e-20 -show_gis -outfmt 6 -max_target_seqs 1000 -dbsize 100000000 -comp_based_stats F -seg no) against the arCOG database (version from 2014)^102^. Protein alignments were generated for each individual protein family using MAFFT L-INS-i^88^ (protein alignments with <1000, settings: --reorder) or MAFFT (alignments with >1000 sequences, settings: --reorder). Afterwards, all alignments were trimmed using BMGE^89^ (settings: -t AA -m BLOSUM30 -h 0.55). Phylogenetic trees were generated using IQ-TREE^50^ (settings: -m LG+G -wbtl -bb 1000 -bnni). To search for HGT events, trees were analyzed using the script count_sister_taxa.py (https://github.com/Tancata/phylo/blob/master/count_sister_taxa.py), which allows to determine the closest sister lineage of any taxon of interest in an unrooted phylogeny, i.e. the sister lineage is defined as the lineage in the smallest sister group while all other taxa are the outgroup. Cases in which homologs of Undinarchaeota formed a sister lineage to non-DPANN, were visually confirmed and all hits for Undinarchaeota are listed in **Supplementary Tables 16-17**.

### Phylogenetic analyses of 16S and 23S rRNA genes

16S and 23S rRNA gene sequences were identified in all archaeal genomes using Barrnap v0.9^103^ (settings: --kingdom arc --evalue 1e-20; https://github.com/tseemann/barrnap). Partial sequences and sequences shorter than 400 bp were removed prior to extracting the sequences using bedtools v2.26.0^104^. Subsequently, sequences were individually aligned using MAFFT L-INS-i^88^ and trimmed using TrimAL^105^(v1.2rev59, -automated1 setting) or BMGE^89^ (settings: -t DNA -m DNAPAM100:2 -h 0.55) and concatenated (https://github.com/nylander/catfasta2phyml). 16S and 23S rRNA gene homologs located on the same contig were concatenated whenever more than one copy of the 16S or 23S rRNA gene was encoded by a genome. Alignment_pruner.pl (https://github.com/novigit/davinciCode/blob/master/perl) was used to remove the 10% most heterogenous sites from the alignment. After running an initial tree with all 16S-23S rRNA gene sequences using IQ-TREE, we confirmed that MAGs with multiple copies clustered together and removed all but one representative copy for the following tree. Additionally, we removed 16S and 23S rRNA homologs of the genome-reduced Huberarchaeota to avoid potential long-branch attraction artifacts. A phylogenetic tree was generated with IQ-TREE for the full-length alignment trimmed with TrimAL^105^ (setting: -automated1), BMGE^89^ (settings: -t DNA -m DNAPAM100:2 -h 0.55) as well as the 10% pruned TrimAL alignment (IQ-TREE; settings: -m GTR+G -bb 1000 -alrt 1000)^50^. The trees are provided in **Supplementary Figures 3-5**.

### Phylogenetic analyses of the RuBisCO gene

RuBisCO protein sequences were identified by their arCOG ID (arCOG04443), which was identified in all Undinarchaeota as described in section ‘Gene calling and annotation’. Subsequently, we extracted the sequences for the RuBisCO protein from Undinarchaeota and combined them with an unmasked alignment generated in a previous study^67^ using mafft_align^88^ with the --add option. The alignment was trimmed using BMGE (-g 0.5 -b 1 -m BLOSUM30 -h 0.55)^89^ and a tree generated using IQ-TREE (-m LG+G - wbtl -bb 1000 -bnni). The conservation of the catalytic site was compared in reference to *Synechococcus elongatus* PCC_6301 as described previously^67^. The tree is provided in **Supplementary Figure 59**. The logo for the catalytic site was generated with logomaker v0.8 in python v2.7.15^106^.

### Phylogenetic analyses of the archaeal DNA primase

To compare the phylogenetic history of the canonical and the fused PriL and PriS subunits of the primase we extracted arCOGs corresponding to the two subunits (arCOG04110 for PriS and arCOG03013 for PriL, respectively) from the 364 reference archaeal genomes. To determine the conserved domains for PriS and PriL in both the fused and canonical proteins we uploaded the sequences to the Conserved Domain Search Service (CD-Search) web-based tool using default settings (**Supplementary Table 11**). The start and end positions provided by CD-Search were used to split the fused version of the primase into “unfused” PriS and PriL subunits using bedtools getfasta. The sequences for the canonical PriS and PriL as well as split PriS and PriL were combined and aligned using MAFFT L-INS-i^88^(option --reorder) and trimmed using TrimAL^105^ (-gappyout). A phylogenetic tree was generated with IQ-TREE (-m LG+F+C10 -bb 1000 -alrt 1000)^50^. The tree is provided in **Supplementary Figure 57**.

### Gene calling and annotation

We annotated the 12 MAGs of Undinarchaeota (six from this study) as well as our set of 352 archaeal reference genomes. Gene calling was performed using Prokka^107^ (v1.14, settings: --kingdom Archaea --addgenes --increment 10 --compliant --centre UU --norrna --notrna). For further functional annotation, the generated protein files were compared against several databases, including the arCOGs (version from 2014)^102^, the KO profiles from the KEGG Automatic Annotation Server^108^ (KAAS; downloaded April 2019), the Pfam database^109^ (Release 31.0), the TIGRFAM database^110^ (Release 15.0), the Carbohydrate-Active enZymes (CAZy) database^111^ (downloaded from dbCAN2 in September 2019), the MEROPs database^112^ (Release 12.0), the Transporter Classification Database^113^ (TCDB; downloaded in November 2018), the hydrogenase database^114^ (HydDB; downloaded in November 2018) and NCBI_nr (downloaded in November 2018). Additionally, all proteins were scanned for protein domains using InterProScan (v5.29-68.0; settings: --iprlookup --goterms)^115^.

Individual database searches were conducted as described in the following section. ArCOGs were assigned using PSI-BLAST v2.7.1+ (settings: -evalue 1e-4 -show_gis -outfmt 6 -max_target_seqs 1000 - dbsize 100000000 -comp_based_stats F -seg no)^116^. KOs as well as PFAMs, TIGRFAMs and CAZymes were identified in all archaeal genomes using hmmsearch v3.1b2^96^ (settings: -E 1e-4). The Merops database was searched using BLASTp v2.7.1 (settings: -outfmt 6, -evalue 1e-20). For all database searches the best hit for each protein was selected based on the highest e-value and bitscore and all results are summarized for Undinarchaeota in **Supplementary Table 7**. For InterProScan we report multiple hits corresponding to the individual domains of a protein using a custom script (parse_IPRdomains_vs2_GO_2.py). Additionally, tRNA genes were identified from contigs of Undinarchaeota and all archaeal reference genomes using tRNAscan-SE v2.0^117^ and the results are summarized in **Supplementary Table 10**.

Proteins potentially involved in cell-cell interactions (**Supplementary Table 20**, some of which were reported in Castelle et al., 2018^13^) were separately screened with HHpred^118^ and Phyre2^119^. HHpred was run on a local server by first running hhblits the uniclust30_2018_08 database (settings: -E 1E-03, - oa3m). The alignment file in a3m format was used as an input in hhsearch against the pdb70 database (settings: -p 20 -Z 250 -loc -z 1 -b 1 -B 250 -ssm 2 -sc 1 -seq 1 -dbstrlen 10000 -norealign -maxres 32000 -contxt context_data.crf -blasttab). Protein homology was investigated using Phyre2 by using the batch upload function of the web-version.

### Metabolic comparisons

Results from the gene calling and annotation described above were used as basis for comparative genome analyses and metabolic comparisons. For simplicity we reported gene presence/absence patterns on class level whenever possible, while DPANN and most taxa without cultured representatives were defined at the phylum level. First, the occurrence of each individual gene found across each MAG/reference genome was counted in R (v3.5.0). This count table provided the basis for summary tables generated using the ddply function of the plyr package (v 1.8.4). The results of these analyses are summarized in **Supplementary Tables 9 and 12**. To plot the heatmap shown in **Figure 3**, the count table was first transferred to a presence/absence matrix and the ddply function was used to summarize the counts across each phylogenetic cluster. The data was visualized as a heatmap using the ggplot function with geom_tile and facet_wrap of the ggplot2 package v3.0.0. A table summarizing the gene IDs used can be found in **Supplementary Table 19**. The heatmap was manually merged with the collapsed tree in Inkscape v0.91.

### Average amino acid identity

The average amino acid identity (AAI) across all archaeal reference genomes and the 12 Undinarchaeota MAGs was calculated using comparem v0.0.23 (settings: aai_wf)^120^. The output was summarized in R (v3.5.0) using the packages reshape2 (v1.4.3), plyr (v1.8.4) and dplyr (v0.7.6) generating the **Supplementary Table 3** and ggplot2 (v3.0.0) and ggpubr (v0.2) to plot the data.

### Undinarchaeota co-occurrence analysis

37 SRA metagenomes (**Supplementary Table 1**) containing reads assigned to Undinarchaeota were pseudo-aligned to a genome dataset consisting of 6,890 GTDB r89 archaeal and bacterial genus-dereplicated genomes (GTDB; https://gtdb.ecogenomic.org) and 12 Undinarchaeota MAGs using the software Kallisto v.0.44.0 with the default k-mer size (31 bp)^121^. To reduce the number of false positive genomes, the pseudo-alignment results were subjected to three filtering criteria published previously^121^. The subset of genomes that fulfilled the three criteria was used to create a reference database for read mapping of the 37 SRA metagenomes via bamM using default parameters (https://github.com/Ecogenomics/BamM). Relative abundances (Ar) were normalized based on the total read counts, i.e. the maximum number of reads of all metagenomes (Nm) divided by the sample read count (Ns), multiplied by number of reads of that particular sample that mapped to the genome (r), and the average read length (l), divided by the genome size (g) (Ar=Nm/Ns * (r*l/g))^25,79^. Proportionality was analyzed using the R package propR^79^ based on normalized relative abundances, followed by the centered log-ratio transformation. Only metagenomes containing maximum UAP2 normalized relative abundance above 1.0 were included in this step (that resulted in 27 metagenomes involved in calculating proportionality). Finally, genomes that were proportional (ρ ≥ 0.9) to more than three Undinarchaeota MAG were identified as Undinarchaeota co-correlated.

## Data and Code availability

The datasets analyzed during the current study are available in the NCBI repository at https://www.ncbi.nlm.nih.gov/ and the accession numbers can be found in Supplementary Table 1. New MAGs generated from the current study have been deposited at NCBI GenBank and can be accessed under BioProject **PRJNA609027** as well as via our repository at Zenodo. Furthermore, additional supplementary files including: (a) genome files including the contigs and protein files for all 12 Udinarcheota MAGs as well as the 352 reference genomes, (b) phylogenies for the species and single gene tree analyses including protein files, alignments and treefiles (Supplementary Figures 6-56) and (c) workflows for the annotations and phylogenies as well as custom python and R-scripts (to analyze tree files), perl code (to parse data) and R codes (to parse the data and generate figures) have been deposited at Zenodo with the **doi: 10.5281/zenodo.3672835**.

## Acknowledgements

This work was supported by a grant of the Swedish Research Council (VR starting grant 2016-03559 to AS), the NWO-I foundation of the Netherlands Organization for Scientific Research (WISE fellowship to AS), and an Australian Research Council (ARC) Future Fellowship (FT170100213, awarded to CR). TW was supported by a Royal Society University Research Fellowship. BW was supported by the Australian Research Council Discovery Early Career Research Awards #DE160100248.

We want to thank Damien de Vienne for helpful discussion regarding testing of protein tree congruency as well as Celine Petitjean for sharing genomic and transcriptomic data of a selection of representative eukaryotes. We are also grateful to those that made their metagenomic datasets publicly accessible, including Pierre Offre for the MOCA dataset, Jill Banfield for the aquifer metagenomes, Peer Bork for the Indian Ocean metagenomes and Francisco Rodriguez-Valera for the Mediterranean metagenomes.

