## Supplementary-Information for "Undinarchaeota illuminate the evolution of DPANN archaea"

### Table of Contents


Table of Contents 2

General 3

- Evaluating CheckM completeness estimates 3
- Screening for contaminants 3

Phylogenetic analyses 4

Informational processing and repair systems 7

- Replication and cell division 7
- Transcription 7
- Translation 8
- DNA-repair and modification 9
- Stress tolerance 9

Metabolic features 10

- Central carbon and energy metabolism 10
- Anabolism 13
- Purine and pyrimidine biosynthesis 13
- Amino acid degradation and biosynthesis 14
- Lipid biosynthesis 15
- Vitamin and cofactor biosynthesis 16

Host-symbiont interactions 16

- Genes potentially involved in host-symbiont interactions 16
- Horizontal gene transfer among Undinarchaeota and other microbial lineages 17
- Co-occurrence analyses 18

References 19

Supplementary Figures 1-63 27-89

### General

#### Evaluating CheckM completeness estimates

Out of 147 archaeal marker genes used by CheckM[1] for estimating genome completeness, seven were absent in all twelve Undinarchaeota metagenome-assembled genomes (MAGs) and in some cases were also absent from Cluster 1 and/or Cluster 2 DPANN archaea (see Main Text for definition of the clusters). Here, we briefly want to discuss these marker proteins and the potential consequences for estimating genome completeness. (1) PF01287 encodes translation initiation factor 5A and is absent in all Undinarchaeota. However, when searching for the corresponding arCOG (arCOG04277; **Supplementary Table 9**) it seems that all archaea, including Undinarchaeota, encode this protein suggesting that the PFAM is not ideal to search for the presence of this protein in at least some archaea. (2) Rps27e homologs (arCOG04108, PF01667) are almost uniquely lacking in Undinarchaeota (see discussion below) and thus do not seem to be universal for this lineage. (3) Similarly, *eif*6 homologs (arCOG04176, PF01912) are absent in Undinarchaeota though present in all other archaeal lineages. (4) PF01982/arCOG01904 homologs appear to be absent in Undinarchaeota and most Cluster 2 DPANN archaea with the exception of Nanohaloarchaeota. This protein encodes a CTP-dependent riboflavin kinase (RFK) commonly found in archaea. However, as we discuss below vitamin biosynthesis genes are commonly absent in DPANN and thus proteins involved in vitamin biosynthesis are less ideal to determine genome completeness for this clade. (5) PF04127 homologs are absent in almost all DPANN except Altiarchaeota and encode the coenzyme A biosynthesis bifunctional protein (CoaBC), which is involved in vitamin biosynthesis and thus also expected to be absent in most DPANN archaea. (6) TIGR00432/arCOG00989 homologs are absent in all DPANN archaea with the exception of Nanoarchaeota and Altiarchaeota. The gene encodes the tRNA-guanine(15) transglycosylase, which is involved in a unique archaeal pathway for archaeosine-tRNA biosynthesis. (7) TIGR01213/arCOG01015 is absent in most DPANN with the exception of Aenigmarchaeota and Nanoarchaeota. The gene encodes a tRNA-pseudouridine synthase responsible for synthesis of pseudouridine from uracil-54 and uracil-55. Altogether, these findings suggest that the marker gene set used by CheckM includes a subset of genes, which is absent in a large proportion of DPANN archaea and thus may underestimate the completeness of DPANN archaeal genomes. Therefore, we additionally assessed genome-completeness with a set excluding these seven markers and provide the alternative and perhaps more accurate completeness and contamination estimates in parenthesis in **Table 1** and **Supplementary Table S2**.

#### Screening for contaminants

Contigs were manually investigated for signs of contamination by screening for an abnormal GC-content (~10% difference of average GC-content) and/or taxonomic affiliation based on a BLASTp search against ncbi_nr (**Supplementary Table 7**; details described in the Methods). We noticed that some Undinarchaeota MAGs (e.g. contig GCA_002494525_5) have a region of ~30 proteins that, based on a BLASTp search, show high similarity to proteins encoded on a fosmid classified as uncultured marine group II/II euryarchaeote_KM3_51_D01[2]. However, phylogenetic analyses of marker proteins encoded in this region, revealed that they clustered with homologs of Undinarchaeota rather than with homologs of Marine Group II/III Archaea. Furthermore, we compared the average amino acid (AAI) of the fosmid of KM3_51_D01 with Undinarchaeota, which showed 84% AAI to Undinarchaeales and 59% AAI to Naiadarchaeales, suggesting that this fosmid was incorrectly assigned to Euryarchaeota and should rather be classified as uncultivated archaeal fosmid.

### Phylogenetic analyses

In published phylogenetic analyses, Undinarchaeota (originally named UAP2) branched sister to all other DPANN archaea[3,4]. To evaluate this placement and assess DPANN monophyly, we performed in-depth phylogenetic analyses using different sets of representative archaeal taxa (364 and 127 taxa set) and marker proteins as well as using concatenated 16S and 23S rRNA gene sequences (see below). In brief, the initial protein set was based on a selection of 151 markers used in previous studies, such as ribosomal protein marker sets, the Genome Taxonomy Database (GTDB) and the PhyloSift marker set[4–7]. Notably, initial phylogenetic analyses of single gene trees, including both archaeal, bacterial and eukaryotic homologs of these 151 commonly used markers, revealed that 39 protein did not recover archaeal monophyly suggesting that they are unsuited for concatenated marker protein analyses. In turn, we excluded these 39 markers as well as translation elongation factor aEF-2 (TIGR00490; arCOG01559), which has two paralogs in some archaeal lineages[8], from our initial marker set (**Supplementary Tables 4-5**).

To further assess the suitability of marker proteins for concatenated gene trees, we scored the remaining markers based on the recovery of well accepted monophyletic archaeal taxa defined at the order to phylum level (see Methods), i.e. we penalized markers, whenever any of these clades were paraphyletic: Geothermarchaeota, Halobacteria, Methanonatronarchaeales, Methanomicrobiales, Methanosarcinales, Methanocellales, Methanophagales, Archaeoglobales, Thermoplasmatales, Acidiprofundales, Methanomassiliicoccales, Poseidoniales, Thermoplasmata (unassigned at order level), Pontarchaea (MG-III), Undinarchaeota, Woesearchaeota, Pacearchaeota, NovelDPANN_1 (UAP1), Parvarchaeota, Nanohaloarchaeota, Aenigmarchaeota, Diapherotrites, Huberarchaeota, Micrarchaeota, Altiarchaeota, Methanopyrales, Methanobacteriales, Methanococcales, Desulfurococcales, Sulfolobales, Thermoproteales, Marsarchaeota, Thermococcales, Theionarchaea, Methanofastidiosa, Hadesarchaea, Persephonarchaea, Odinarchaeota, Verstraetearchaeota, Thorarchaeota, Lokiarchaeota, Heimdallarchaeota, Bathyarchaeota, Thaumarchaeota, Korarchaeota, Aigarchaeota, Geoarchaeales, Hydrothermarchaeota and Nanoarchaeota. Violation of monophyly was counted as splits - as described in the methods using a script that we make available in git-bub (count_sister_taxa.py; https://github.com/Tancata/phylo/blob/master/count_sister_taxa.py) - which provided a mean to rank the marker proteins based on congruency and potential events of horizontal gene transfer (HGT). Please note, that we did not make any a priori assumptions regarding the relationship of any of these clades with each other, i.e. our markers did not require that certain clans such as the DPANN are monophyletic. This is important because DPANN monophyly remains debated and we did not want to bias our marker protein assessment. Subsequently, concatenated alignments were created by combining the 25%, 50% and 75% highest (least amount of potential HGTs) - as well as 25% and 50% lowest-scoring (highest amount of potential HGTs) marker proteins (**Supplementary Tables 4-5**). These datasets were subjected to a variety of Bayesian and Maximum-likelihood-based phylogenetic analyses that differed with respect to model as well as data treatment, such as removal of fast-evolving or compositionally heterogeneous sites, and the results are shown in **Supplementary Table 6** and **Supplementary Figs. 6-56**.

All our inferences based on the curated marker protein sets recovered a monophyletic DPANN clan and consistently placed Undinarchaeota as an independent lineage branching between two monophyletic DPANN clans, here referred to as Cluster 1 DPANN archaea including Altiarchaeota, Micrarchaeota and Diapherotrites and Cluster 2 DPANN archaea including Nanoarchaeota, Pacearchaeota, Woesearchaeota, Huberarchaeota, Parvarchaeota and Nanohaloarchaeota (**Supplementary Figs. 6-41, Supplementary Table 6**). In particular, the consistent placement of Altiarchaeota within Cluster 1 is notable, since the evolutionary history of this archaeal lineage remains an ongoing matter of debate[9–13].

Subsequently and as detailed in the main text, we addressed the effect of using highly incongruent markers with a high degree of splits (i.e. non-monophyletic archaeal taxa, see above) on phylogenetic analyses by inferring phylogenetic trees using 25% and 50% of the lowest ranking markers (**Supplementary Tables 4-5)**. Notably, resulting phylogenetic trees recovered topologies inconsistent with the accepted archaeal taxonomy and confirming the unsuitability of these markers due to conflicting evolutionary signals (**Supplementary Figs. 46-49**). For instance, in trees inferred using the 25% lowest ranking marker set, known symbionts, such as Nanoarchaeota and Nanohaloarchaeota, clustered with their crenarchaeal or halobacterial hosts, respectively (**Supplementary Fig. 46-47**). Based on our investigation of horizontal gene transfer events, ~7% and ~12.4% of the proteome of Halobacteria and Nanohaloarchaeota, respectively, have been exchanged horizontally or cluster together due to similar compositional biases (Figure 4a, **Supplementary Table S17**). Importantly, this suggests that conflicting signals regarding the placement of certain archaeal clades, in particular members of the DPANN archaea, may in part be due to the use of unsuitable markers in phylogenetic reconstructions based on protein concatenations.

Considering that previous analyses[3,4] suggested that the Undinarchaeota lineage represents an outgroup of all DPANN lineages, we further addressed the reliability of our inferences by accounting for effects imposed by fast-evolving[14,15] or compositionally heterogeneous sites[16]. In particular, and to ensure that the placement of Undinarchaeota is not affected by such artefacts, fast-evolving and compositionally heterogeneous sites were removed using the SlowFaster method[14] and chi2 testing[17], respectively. In brief, 10%, 20%, 30% and 40% of the most biased sites were removed from alignments generated using the 25% and 50% highest ranked marker proteins both from the 127 and 364 taxa set (**Supplementary Table 6**, **Supplementary Figs. 13-22 and 30-40**). All analyses confirmed the placement of Undinarchaeota as an independent lineage emerging between DPANN Cluster 1 and Cluster 2 DPANN archaea. Furthermore, these analyses confirmed the placement of Altiarchaeota within Cluster 1 DPANN. The most notable difference between the treated and untreated alignments was the placement of Methanonatronarchaeia[18]. In agreement with a recent study addressing the effect of compositional biases and/or fast-evolving sites[19], our analyses indicated that Methanonatronarchaeia form a sister lineage of Halobacteria in full alignments, while forming an early branching Methanotecta lineage when biased sites are removed (**Supplementary Figs. 13-22, 31-35 and 37-38**). In turn, it seems likely that the sisterhood of Halobacteria and Methanonatronarchaeia is due to a phylogenetic artifact perhaps resulting from convergent sequence adaptations to high salinity.

Next, we investigated tree topologies inferred from several non-curated alignments as well as marker protein sets such as a ribosomal marker set[4], the GTDB marker set[5] and the PhyloSift markers[6] as well as 16S/23S rRNA genes. Unexpectedly, these trees not only showed inconsistent placements for the Undinarchaeota lineage but also for the Altiarchaeota. First of all, an analysis based on an alignment of 34 concatenated marker proteins of the PhyloSift marker protein set (alignment length of 5,353 amino acids, **Supplementary Fig. 50**), placed Altiarchaeota as a separate archaeal lineage emerging in-between DPANN (including Undinarchaeota) as a whole and all other Archaea, respectively (support values of 87.7/89). In this case, the Undinarchaeota lineage represented the first diverging DPANN lineage. Secondly, an alignment consisting of 14 concatenated ribosomal proteins (alignment lengths of 1,974 (trimmed with BMGE) and 2,406 (trimmed with trimAL)) recovered Undinarchaeota as an independent lineage emerging in-between all DPANN (including Altiarchaeota) and all other Archaea (support values of 99.7/100 and 99.2/99), respectively (**Supplementary Figs. 53-54**) as has been assumed previously[3,4]. Thirdly, in phylogenetic trees recovered from the complete set of 122 archaeal marker proteins used by GTDB (**Supplementary Figs. 51-52**), Altiarchaeota formed a separate branch forming a sister group to all DPANN Archaea (bootstrap supports of 85.3/98 and; using IQ-tree and bootstrap support 1; using FastTree). Here, Undinarchaeota branched as a sister lineage of Cluster 2 DPANN (**Supplementary Table 6**). Finally, we performed a concatenated analysis of the 16S and 23S rRNA genes using the 364 taxa set. In contrast to protein phylogenies, these analyses (depending on trimming method and alignment filtering) recovered Undinarchaeota in between Cluster 1 and Cluster 2 DPANN archaea (**Supplementary Fig. 4**; 93.7/80) or as a sister lineage of the Aenigmarchaeota (**Supplementary Fig. 3**; 95.7/86), while Altiarchaeota formed a monophyletic cluster with Micrarchaeota and Diapherotrites (88.1/90 bootstrap support) (**Supplementary Figs. 3-5**). However, these latter analyses recovered several unexpected groupings and trees were characterized by overall low support in deeper branches and were likely affected by long-branch attraction artifacts (LBA)[20]. In addition, alignments of the concatenated 16S/23S rRNA were relatively short (3,128 nucleotides) and likely harbored insufficient information to resolve deeper branches in the tree as indicated by low support values. Finally, 16S/23S rRNA genes have previously been shown to be good molecular thermometers that reflect the optimal growth temperature of organisms[21,22]. In turn, the sequence composition of rRNA genes can be biased and impair the accurate assessment of phylogenetic relationships.

These analyses highlight the importance of carefully assessing the suitability of marker protein sets as well as of determining the effect of compositional biases on phylogenetic inferences aiming to address archaeal evolution[19,23]. In particular, many commonly used marker protein sets seem to comprise protein families unsuitable for concatenation: for instance, single-protein tree analyses of the PhyloSift and 122 GTDP markers not only revealed that some markers were exchanged horizontally with bacteria (violation of archaeal monophyly) but also indicated that several of these markers failed to recover monophyly of well-accepted archaeal order- to phylum-level lineages indicating horizontal exchange (**Supplementary Tables 4-5**).

Altogether, our extensive phylogenetic reconstructions provide strong support for a clan consisting of Undinarchaeota as sister lineage of DPANN Cluster 2 archaea, as well as the placement of Altiarchaeota as part of a clan comprising Cluster 1 DPANN archaea. Monophyly of the DPANN clan in turn solely depends on the placement of the root (see Main Text).

### Informational processing and repair systems

A common feature of symbionts, particularly bacterial endosymbionts, is the loss of genes related to energy production, general biosynthetic pathways, DNA repair mechanisms and to a lesser degree informational processing[24]. To elucidate whether similar patterns characterize the evolution of Undinarchaeota and DPANN archaea in general, we investigated the presence and absence of core genes involved in replication, transcription, translation and DNA and repair processing.

#### Replication and cell division

Undinarchaeota representatives encode most genes related to replication processes (**Supplementary Fig. 58**, **Supplementary Tables 7-10**). More specifically, Undinarchaeota MAGs encode two DNA polymerases: DNA polymerase B1 (PolB1, arCOG00328) and the two subunits of the DNA Polymerase D (arCOG04447 and arCOG04455). Additionally, two of the four aquifer representatives encode a DNA polymerase IV (Dpo4, arCOG04582). Furthermore, Undinarchaeota MAGs encode all replication-related proteins commonly found in archaea including ORC1-type DNA replication protein 1 (Orc1, arCOG00467) and several helicases including the potential replicative helicase Mcm2 (arCOG00439) and a DNA ligase (Lig, arCOG01347). Notably and in agreement with all other analyses and previously published results[25], Cluster 2 DPANN archaea and Undinarchaeota encode a fused version of the DNA primase, i.e. PriS and PriL are encoded by one gene (**Supplementary Fig. 57**, **Supplementary Table 11**). Finally, they encode DnaG (arCOG04281) and two topoisomerases: type 1 (TopA; arCOG01527) and type 2 topoisomerase 6 (Top6AB; arCOG04143 and arCOG01165), while lacking genes for gyrase or reverse gyrase. However, this is not unexpected since gyrases are more sparsely distributed in archaea (**Supplementary Figure 58**)[26,27]. In particular, reverse gyrase seems to be uniquely found in hyperthermophiles (with an optimal growth temperature above 80°C) and thus represents a genetic marker for the adaptation to life at high temperatures[13,28–32]. In turn, the absence of reverse gyrase homologs in the herein analyzed MAGs suggests that they may not comprise hyperthermophiles.

Each Undinarchaeota MAG encodes two paralogs for FtsZ cell division proteins (arCOG02201) as well as a septum site-determining protein MinD (K03609). Additionally, Undinarchaeota representatives encode histones (arCOG02144), chromosome segregation and condensation protein SpcA (K05896)[33] as well as the chromosomal protein MC1 (arCOG04743). MC1 is only found in a small number of archaeal lineages and plays a role in DNA bending and compaction in certain Euryarchaeota[34–36].

#### Transcription

With few exceptions, Undinarchaeota MAGs encode most proteins involved in transcription (**Supplementary Fig. 60**, **Supplementary Tables 7-9**), including all core subunits of the RNA polymerase (RpoA1; arCOG04257, RpoA2; arCOG04256, RpoB; arCOG01762, RpoD; arCOG04241, RpoK; arCOG01268, RpoL; arCOG04111, RpoF; arCOG01016, RpoH; arCOG04258, RpoE; arCOG00675, RpoN; arCOG04244 and RpoP; arCOG04341). Similar to many other archaea, Undinarchaeota representatives lack genes for RpoG (arCOG04271) and Rpo13 (arCOG05938)[37], which do not seem to be required to form a functional RNA polymerase[38]. All transcription factors common to archaea, such as the transcription initiation factor IIB (Tfb, arCOG01981), transcription factor S (Tfs; arCOG00579) and transcription factor E (Tfe, arCOG04270), are present in Undinarchaeota. Interestingly, Tfs is found in Undinarchaeota and Cluster 2 but not Cluster 1 DPANN archaea (**Supplementary Fig. 60**). Few non-essential genes related to transcription appear to be absent in Undinarchaeota MAGs: these include genes coding for DNA/RNA-binding protein Alba1 (arCOG01753) (only present in two aquifer Naiadarchaeales MAGs), as well as the complete absence of Rad3-related DNA helicase (DinG, arCOG00770) in Undinarchaeta, which is involved in nucleotide excision repair[39].

#### Translation

Undinarchaeota MAGs seem to encode most proteins involved in translation (**Supplementary Fig. 61**, **Supplementary Tables 7-10**). For example, Undinarchaeota MAGs encode most ribosomal proteins and all archaeal tRNA synthetases. Two notable exceptions are Rpl30e (arCOG01752), which is absent in marine Undinarchaeales, and Rps27e (arCOG04108), which is lacking in all Undinarchaeota representatives. Rpl30e seems also absent in other archaea, such as Halobacteria, Nanohaloarchaeota and most Thermoplasmatales, and may therefore not be essential for building a functional ribosome[40,41]. On the other hand, Rps27e is found in all analyzed cultivated archaeal taxa and absent in only a subset of so-far uncultivated archaeal lineages such as all Undinarchaeota, Persephonarchaea (MSBL1), Poseidoniales (MG-II) and Pontarchaea (MG-III). Rps27e is likely involved in rRNA processing and is thought to be universally present in archaea and eukaryotes but absent in bacteria[40,42]. It remains to be determined to what extent the absence of Rps27e in Undinarchaeota impairs the functioning of their ribosomes.

Undinarchaeota MAGs also encode most translation-related proteins such as the initiation factors EifA (arCOG01179), Eif2 (arCOG04107), Eif5 (arCOG04277), the elongation factors Tuf (arCOG01561), EF1b (arCOG01988) and FusA (arCOG01559) as well as the potential terminating factor eRF1 (arCOG01742). Other proteins involved in translation that are present in Undinarchaeota MAGs include the ribonuclease P complex (Rnp1-4), ribonuclease Z (Rnz, arCOG00501), ribonuclease J (RnjA, arCOG00546) and endoribonuclease (Nob1, arCOG00721). However, and in contrast to most other archaea, Undinarchaeota seem to lack the translation factor Eif6 (arCOG04176), which has a ribosomal anti-association activity[43], and the potential translation initiation ATPase Rli1 (K06174), which plays a role in the dissociation of the two ribosomal subunits[44,45]. HflX (arCOG00353), which potentially plays a role in ribosome recycling in bacteria and some archaea[46], is also absent in Undinarchaeota representatives and it is unclear whether any other proteins can complement the function of these proteins.

Undinarchaeota MAGs encode the exosome subunits Rrp4/41/42 (arCOG00678, arCOG01575 and arCOG01574) but lack Csl4 (arCOG00676) as well as any Csl4-domain containing proteins (i.e. IPR039771, IPR030850). Csl4 plays a role in specificity and regulation of RNA processing[47,48] and interacts with DNA primase DnaG (arCOG04281)[49]. Experimental evidence suggests that a Csl4-Rrp4/41/42 complex (Csl4-exosome) degrades an oligo-A tail more effectively than a Rrp4-Rrp4/41/42 complex (Rrp4-exosome), suggesting that a complex without Csl4 might be still functional but possibly less efficient than the full complex[47]. Notably, the presence of a putative Rrp4-exosome without Csl4, seems to be a distinctive feature of Undinarchaeota and Cluster 2 DPANN archaea (**Supplementary Fig. 61**, **Supplementary Table 9**) and indicates a structural difference of their exosomes as compared to Cluster 1 DPANN and other archaea.

Undinarchaeota MAGs encode the necessary proteins to synthesize translationally modified residues including diphthamide (via Dph2/5/6; arCOG04112, arCOG04161 and arCOG00035)[8]. However, Undinarchaeota lack most genes related to the pathway for wyosine derivatives. Wyosine is important for post-translational modifications at position 37 of the phenylalanine-specific transfer RNA (tRNAPhe) that is common in archaea and eukaryotes[50]. Specifically, while Undinarchaeota MAGs encode a potential Trm5 methyltransferase (arCOG00033), they lack the three other biosynthesis genes required for this pathway (Taw1/2/3; arCOG04174, arCOG10124 and arCOG04156). This finding suggests that members of this group can only methylate the N1 position of guanosine-37 and form m1G37 but no other derivatives.

Finally, Undinarchaeota MAGs encode several proteins involved in post-translational processes that include the two subunits required to assemble a potential proteasome (PsmAB, arCOG00971 and arCOG00970) as well as a chaperon of the HSP20-family (arCOG01832) and the thermosome chaperonin (ThsA; arCOG01257).

#### DNA-repair and modification

A common feature of bacterial organisms with reduced genomes is the absence of key proteins involved in DNA-repair[24]. However, Undinarchaeota MAGs appear to harbor most proteins related to DNA repair that are typically found in archaea (**Supplementary Fig. 58**, **Supplementary Tables 7-10**). Proteins related to recombination and repair that were detected in Undinarchaeota representatives include a holliday junction resolvase (Hjc, arCOG00919), a 5'-3'_exonuclease (Fen1, arCOG04050), type III and V endonucleases (Nth; arCOG00459 and Nfi; arCOG00929), the DNA repair and recombination proteins RadAB (arCOG00415, arCOG00417), the double-strand break repair protein Rad50/Mre11 complex (SbcCD; arCOG00368 and arCOG00397) and the GroEL chaperonin (arCOG01257). However, Undinarchaeota MAGs seem to lack genes for a single-stranded-DNA-specific exonuclease (RecJ; arCOG00427), the Chaperone DnaK (arCOG03060) and the ATP-dependent DNA helicase (DinG, arCOG00770), which are otherwise found in most Archaea including most DPANN archaea. It remains to be assessed, whether other DNA helicases encoded by Undinarchaeota MAGs, such as the DNA double-strand break repair helicase (HerA, arCOG00280) or the uncharacterized ATP-dependent helicase (Lhr, arcog00557), may function in DNA repair and functionally substitute for the lacking enzymes mentioned above.

A few proteins related to DNA repair show lineage-specific distributions across the Undinarchaeota representatives. For example, only marine Undinarchaeales but not aquifer Naiadarchaeales MAGs encode the alkylation repair enzyme AlkD (arCOG05122) and the exodeoxyribonuclease III (Xth, arCOG02207) as well as a type 4 uracil-DNA glycosylase (Ugd1m; arCOG00905), which is likely involved in base excision repair[51]. Naiadarchaeales MAGs on the other hand encode a methylated-DNA--protein-cysteine methyltransferase (Ogt, arCOG02724), involved in the repair of alkylation damage[52] and an 8-oxoguanine DNA glycosylase (Ogg, arCOG04357) that is involved in repairing oxidative DNA damage[53].

#### Stress tolerance

Since it is likely that Undinarchaeota encounter fluctuating environmental conditions in marine and aquifer habitats, we next investigated each MAG for the presence of proteins involved in stress tolerance other than repair-related proteins (**Supplementary Tables 7-10**). Marine Undinarchaeales representatives encode two heat-shock proteins of the HSP20 family[54] (arCOG01832 and arCOG01833), two potential glutaredoxins (arCOG02607, arCOG02608) and a Fe/Mn-containing superoxide dismutase (SodA, arCOG04147). Superoxide reductases, which are often found in anaerobic and microaerophilic microorganisms[55], appear to be lacking. Two of the aquifer Naiadarchaeales MAGs encode peroxiredoxins (arCOG00310, arCOG00312), which might be involved in oxidative stress tolerance[56] and most Undinarchaeota MAGs encode a thioredoxin system (TrxAB; arCOG01972 and arCOG01296) that may play a role in counteracting fluctuations in nutrient availability and oxygen status[57]. Overall, this suggests that Undinarchaeota utilize several systems to encounter oxidative stress.

### Metabolic features

#### Central carbon and energy metabolism

**Carbon metabolism.** Next, we investigated key pathways of representatives of Undinarchaeota regarding carbon metabolism and potential modes of energy conservation. All Undinarchaeota MAGs have a low number of genes encoding carbohydrate and peptide transporters as well as carbohydrate degradation enzymes (CAZymes). The only putative transporters were PotE (arCOG00009), involved in amino acid transport, an uncharacterized solute transporter (arCOG00238; IPR001898), which might take up sulfate and/or dicarboxylate with the concomitant uptake of sodium ions[58], a phosphate transporter (PitA, arCOG02267) and a potential tripartite tricarboxylate transporter (TTT; arCOG04469, PF01970; **Supplementary Tables 7, 9, 13**). According to the Transporter Classification DataBase (TCDB), the TTT transporter is homologous to TctA (TCDB ID 2.A.80.2.1), which is predicted to be a putative citrate transporter[59]. Additionally, all marine Undinarchaeales MAGs encode cation anti-/symporters, such as the Na(+)/H(+) antiporter (KefB; arCOG01953) and the sodium:calcium antiporter (arCOG02881), which may be used to maintain osmic balance in marine environments[60,61]. Consistently, KefB is only encoded by two aquifer MAGs and arCOG02881 was completely absent in Naiadarchaeales.

While Undinarchaeota MAGs encode a small number of peptidases, these may be involved in anabolism rather than catabolism (see below) (**Fig. 3, Supplementary Tables 7-9 and 15**). In fact, we did not detect any signal peptide in any of the predicted peptidases (as determined by InterProScan including a SignalP search) suggesting that peptidases are located intracellularly. In turn, the most likely substrates used for central metabolism and energy conservation seem to be simple carbohydrates, such as pyruvate or acetate, or nucleic acids. While Undinarchaeota representatives lack specific sugar transporters, simple sugars could perhaps be taken up by passive diffusion[62]. Nucleic acids might be taken up via pili (encoded by Undinarchaeota MAGs, see below) and degraded into nucleosides via nucleases, such as ribonuclease J (RnjA, arCOG00546), ribonuclease HII (RhnAl; arCOG02942 and RhnB; arCOG04121)[63], exonuclease III (XthA, arCOG02207)[64], ATP-dependent RNA helicase (DeaD, arCOG00558), or endonuclease YncB (arCOG03192)[65]. Nucleoside triphosphates might be fed into the nucleoside degradation pathway via the AMP phosphorylase (DeoA, arCOG02013), ribose 1,5-bisphosphate isomerase (arCOG01124) and ribulose 1,5-bisphosphate carboxylase (RbcL; RuBisCO, arCOG04443), which would yield 3-phosphoglycerate[66–68]. This pathway has been discussed to be relevant for a range of DPANN archaea in previous studies[68,69]. Two Naiadarchaeales MAGs seem to encode a functional group III-b RuBisCO with the same key catalytic residues as found in the oxygen-sensitive RuBisCO found in *Methanocaldococcus jannaschii*[70] (**Supplementary Figure S59a-c**). MAGs from marine Undinarchaeales on the other hand encode a group-III-like RuBisCO homolog that, with the exception of the homolog of MAG GCA_002502135, shows substitutions at the catalytic site in position 195 (aspartic acid (D) to glutamic acid (E)) and 196 (phenylalanine/leucine/tyrosine (F/L/Y) to glycine (G)) (**Supplementary Fig. 59c**). The substitution of D by E does not change the property of the side chains, which are both acidic and in turn have a negative charge. The main difference between these two amino acids is the length of the side chain, with E having one methyl-group more than D. On the other hand, even though both F/L/Y and G (Phenylalanine/ Leucine/ Tyrosine and Glycine) are neutral, the side chains of F and Y differ from those of L and G with respect to class (aliphatic versus aromatic). In turn, it remains to be determined, whether the substitution of D by E at position 195 could be compensated by the change of F/L/Y to G at position 196 and whether the RuBisCO-like proteins of marine Undinarchaeales have retained their canonical function. However, the presence of a conserved group-III RuBisCO and other major key genes of the AMP degradation pathway suggests that at least some aquifer Naiadarchaeales might use this pathway to produce 3-phosphoglycerate that can enter the central carbon metabolism. However, it has to be noted that DeoA was only found in one and the RuBisCO only in two out of four Naiadarchaeales genomes. With the exception of MAG SRR2090159.bin1129 all had a relatively low completeness, such that the absence of these genes in several Naiadarchaeales could be either due to genome completeness or be signs of genome streamlining.

The gene repertoire of Undinarchaeota suggests that 3-phosphoglycerate produced by the AMP degradation pathway could enter the lower glycolytic pathway via their phosphoglycerate mutase (GpmA; arCOG01993 and ApgM; arCOG01696), enolase (Eno, arCOG01169) and phosphoenolpyruvate synthase (PpsA, arCOG01111), which would yield pyruvate and ATP. Pyruvate kinase (Pk, arCOG04120), which in most archaea converts phosphoenolpyruvate to pyruvate, is absent from Undinarchaeota representatives. The only enzyme that might fulfill this role in Undinarchaeota is PpsA, which is encoded by these MAGs and may be reversible, as reported in certain archaea[71–73]. Additionally, 9 out of 12 Undinarchaeota representatives encode a putative pyruvate dehydrogenase complex (PdhABC; arCOG01054, arCOG01052 and arCOG01706) (found also in some other DPANN archaea (**Supplementary Table S9**)[74]), while lacking genes for oxoacid-ferredoxin oxidoreductase complexes (e.g. OorABDG; K00174 to K00177, arCOG01599 to arCOG01608), which are common in other archaeal lineages[75]. Though the functional annotation of pyruvate dehydrogenases is challenging based on sequence information alone, it seems possible that Undinarchaeota members use this complex for the conversion of pyruvate to acetyl-CoA.

Acetyl-CoA could be further metabolized to acetate via the ADP-forming acetyl-CoA synthetase (AcdAB; arCOG01340 and arCOG01338), a reaction that would allow the production of ATP. Additionally, three MAGs of the aquifer Naiadarchaeales encode a putative aldehyde as well as alcohol dehydrogenase (PF00171 as well as PF08240 and arCOG01455, respectively). While none of the Undinarchaeota homologs are closely related to experimentally characterized enzymes, it seems possible that these Undinarchaeota representatives are able to ferment 3-phosphoglycerate to pyruvate (generating ATP via the lower glycolytic pathway) and ethanol (to remain redox balance).

Key enzymes of the glycolytic pathway, the 6-phosphofructokinase (Pfk, arCOG03370) and pyruvate kinase (Pk, arCOG04120), appear to be absent in Undinarchaeota MAGs (**Fig. 3**, **Supplementary**

**Tables 7-9 and 12**). Additionally, representatives of the Undinarchaeota lack genes for key proteins of the classical and modified versions of the Entner–Doudoroff (ED) pathway such as the gluconate dehydratase (Gad, arCOG01168) or 2-dehydro-3-deoxy-D-gluconate/2-dehydro-3-deoxy-phosphogluconate aldolase (K11395). The absence of the upper glycolytic pathway and ED-pathway coincides with the absence of carbohydrate-active enzymes belonging to the glycoside hydrolase (GH) family (**Supplementary Table 14**), indicating that members of Undinarchaeota are unable to utilize complex carbohydrates as carbon or energy sources. Additionally, Undinarchaeota MAGs lack most genes encoding enzymes linked to the TCA cycle. While representatives of the marine Undinarchaeales might be able to convert oxaloacetate to malate using malate dehydrogenase (Mdh, arCOG00246), MAGs of the aquifer Naiadarchaeales encode a malic enzyme (MaeA, arCOG00853) that might convert malate to pyruvate.

Undinarchaeota MAGs encode genes for the initial steps of gluconeogenesis, i.e. the gluconeogenic enzymes that allow the conversion of pyruvate to fructose-6-phosphate, including the phosphoenolpyruvate synthase (PpsA, arCOG01111) and the bifunctional fructose-1,6-bisphosphate aldolase/phosphatase (Fbp, arCOG04180). While Fbp was originally suggested to be present in most archaea[76], based on our analyses Fbp is detected in only a few representatives of the DPANN archaea other than Undinarchaeota and ~40% of genomes analyzed from Altiarchaeota (**Supplementary Table 9**). The presence of Fbp in Undinarchaeota MAGs suggests that members of this group are able to synthesize cellular building blocks via gluconeogenesis using 3-phosphoglycerate. Glyceraldehyde-3-phosphate and fructose-6-phosphate produced during gluconeogenesis might be fed into the non-oxidative pentose-phosphate pathway via a putative transaldolase (TalA, arCOG05061) and transketolase (TktA, arCOG01053) to produce pentoses, such as ribose-5-phosphate. These could subsequently enter into anabolic pathways including the purine biosynthetic pathway.

**Redox balance.** The absence of genes encoding enzymes involved in the oxidative phase of the pentose pathway as well as the lack of isocitrate dehydrogenase (Icd, arCOG01164), NADH dehydrogenases or hydrogenases (**Fig. 3**, **Supplementary Tables 7-9**) suggests that Undinarchaeota may use alternative enzymes to reduce NAD(P)+ to NAD(P)H. One possible candidate enzyme for this conversion is the thioredoxin reductase (arCOG01296, TrxB)[57], genes for which are present in most Undinarchaeota MAGs. Furthermore, Undinarchaeota MAGs encode glyceraldehyde-3-phosphate dehydrogenase (Gap, arCOG00493), and, in the case of the aquifer Naiadarchaeales representatives, also malic enzyme (MaeA, arCOG00853) and a NAD(P)-dependent glyceraldehyde-3-phosphate dehydrogenase (GapN, arCOG01252). All of these proteins could couple NADPH-generation to central carbon metabolism[77]. Another enzyme that plays a role in providing organisms with *de novo* NADP is the NAD kinase (NadK, arCOG01348)[76], which is encoded by all aquifer MAGs. The absence of NadK in marine Undinarchaeota is interesting as most bacteria and archaea encode this enzyme[78]. One of the few characterized organisms that lack NadK is the obligate intracellular bacterium *Chlamydia trachomatis* that seems to rely on its host for NADP maintenance[79]. However, based on our analysis homologs of this enzymes seem also to be lacking in certain free-living archaea such as Methanococcales (**Supplementary Table S9**).

**Energy metabolism.** Undinarchaeota MAGs lack genes encoding membrane-bound complexes belonging to the electron transport chain including NADH dehydrogenases, hydrogenases, cytochromes and terminal oxidases (**Supplementary Table 9**). Additionally, we could not detect any genes for terminal reductases, such as nitrate/nitrite reductase, sulfite reductase or fumarate reductase in the

Undinarchaeota MAGs. However, an archaeal-type V-type ATP synthase is encoded by all Undinarchaeota representatives (AtpABCDEFHI; arCOG00868, arCOG00865, arCOG02459, arCOG04101, arCOG00869, arCOG04102, arCOG03363 and arCOG04138). Furthermore, most Undinarchaeota encoded a pyrophosphate-driven (instead of ATP-driven) sodium pump (HppA, arCOG04949), that may use energy conserved by pyrophosphate hydrolysis for proton movement across the membrane[80–82].

Altogether, our analyses of the central carbon and energy metabolism of the herein reconstructed Undinarchaeota representatives indicate that members of this group are restricted to energy conservation using substrate-level phosphorylation. In fact, considering the limited substrate range and absence of various central carbon metabolic pathways as well as membrane-bound complexes, it seems possible that members of the Undinarchaeota rely on other organisms to sustain their living (see below) - a lifestyle common in members of the DPANN[4,83].

#### Anabolism

##### Purine and pyrimidine biosynthesis

Most aquifer Naiadarchaeales MAGs encode enzymes needed to convert ribose-5-phosphate into inosine monophosphate (IMP) (**Fig. 3**, **Supplementary Tables 7-9 and 12**), while MAGs from marine Undinarchaeales appear to lack key genes encoding proteins involved in purine biosynthesis. For example, phosphoribosylformylglycinamidine cyclo-ligase (PurM, arCOG00639), phosphoribosylaminoimidazole-succinocarboxamide synthase (PurC, arCOG04421) and phosphoribosylaminoimidazole-succinocarboxamide formyltransferase (PurH, arCOG02824) were only found in either two or three marine Undinarchaeales MAGs while being present in most aquifer representatives. However, it has to be noted that not all steps of the purine biosynthesis pathway have been elucidated in archaea. For instance, it is currently unclear which enzyme catalyzes the conversion of 5-aminoimidazole ribonucleotide to N5-carboxyaminoimidazole ribonucleotide[84], which in bacteria is mediated by the N5-CAIR synthase (PurK, arCOG01597). Similarly, genes encoding the guanylate kinase (Gmk, K00942) seem to be absent not only from all Undinarchaeota MAGs but from archaeal genomes in general[85] (**Supplementary Table 9**) and it remains to be determined, which enzyme substitutes this reaction in Archaea.

The genes required to convert IMP to purines have a slightly inconsistent occurrence across marine and aquifer Undinarchaeota MAGs. For example, genes for IMP dehydrogenase (GuaB, arCOG00612), which catalyzes the conversion of IMP to xanthosine 5'-phosphate), were only found in three marine Undinarchaeales representatives (GCA_002495465, SRR4028224.bin17 and SRR5007147.bin71) and one aquifer MAG (SRR2090153.bin1042), while genes for GMP synthase (GuaA, K01951), which catalyzes the second step, were present in two additional aquifer MAGs (SRR2090159.bin1129 and SRR2090159.bin1288). Genes encoding the nucleoside diphosphate kinase (Ndk, arCOG04313) and ribonucleoside-triphosphate reductase (NrdD, arCOG04889), required for the formation of dGTP, could be identified in most Undinarchaeota MAGs. Finally, adenylosuccinate synthetase (PurA, arCOG04387), which is required to convert IMP to adenylosuccinate, was only present in three aquifer MAGs, but the genes encoding the proteins involved in the production of dATP (PurB;

arCOG01747, AdkA; arCOG01039, Ndk; arCOG04313 and NrdD; arCOG04889) were found in most Undinarchaeota MAGs. The lack of GuaB in most aquifer representatives is puzzling, since they harbor genes for most other steps of this pathway. In turn, it remains to be determined whether the absence of this gene is due to genome incompleteness or represent a true biological signal.

Next, our analysis of the pyrimidine biosynthesis pathway revealed that Undinarchaeota MAGs encoded most of the proteins necessary to convert carbamoyl phosphate to CTP and UTP[85]. These proteins include the carbamoyl-phosphate synthase (CarA, arCOG00064) catalyzing the first step in the pathway. Glutamine required by CarA likely cannot be synthesized by Undinarchaeota themselves but must be taken up from the environment (see also discussion on amino acid biosynthesis pathways below). All enzymes required for the conversion of glutamine to uridine monophosphate (UMP), which are encoded by the PyrBCDEF (arCOG00911, arCOG00689, arCOG00603, arCOG00029, arCOG00081) gene cluster, were found in the majority of Undinarchaeota MAGs (**Fig. 4, Supplementary Tables 7-9**). Furthermore, almost all MAGs encode the enzymes that convert UMP further into dCTP (PyrH; arCOG00858, PyrG; arCOG00063, Ndk; arCOG04313 and NdrD; arCOG04889), dUTP (Dcd, arCOG04048) and dTTP (ThyA; arCOG03214 and Tmk; arCOG01891). Notably, we could not identify genes for thymidylate synthase (ThyA; arCOG03214) or flavin-dependent thymidylate synthase (ThyX; arCOG01883) in most aquifer Naiadarchaeales MAGs while these were present in Undinarchaeales representatives. Considering the presence of all other enzymes of this pathway, it seems possible that the lack of *thy*A/X genes is due to genome incompleteness.

##### Amino acid degradation and biosynthesis

Our analyses of the amino acid metabolism suggested that Undinarchaeota representatives lack most genes encoding enzymes required for amino acid biosynthesis and interconversion (**Supplementary Tables 7-9 and 12**). The identified genes code for aminopeptidases that might be involved in the turnover of intracellular proteins or general protein processing. For instance, we found genes for leucyl aminopeptidase (PepA; arCOG04322), methionine aminopeptidase (Map; arCOG01001) and a potential membrane-associated serine protease of the S54 family (GlpG, arCOG01768; IPR022764 and IPR035952)[86]. Aminopeptidases potentially involved in protein hydrolysis include a Xaa-Pro aminopeptidase (PepQ, arCOG01000) and a putative metallopeptidase (arCOG04217). Other enzymes related to amino acid metabolism, which were predicted to be present in Undinarchaeota representatives, could be involved in the interconversion of amino acids and respective organic acids. For example, the putative aspartate aminotransferases encoded by Undinarchaeota MAGs (AspC, arCOG01130) might convert L-aspartate and 2-oxoglutarate to glutamate and oxaloacetate, their serine hydroxymethyltransferase (GlyA, arCOG00070) could be involved in the interconversion of serine and glycine, a glutamate dehydrogenase (GdhA, arCOG01352) might produce glutamate without ammonia assimilation[87] and a cysteine desulfurase might interconvert cysteine and alanine (SufS, arCOG00065). The SufS of Undinarchaeota representatives may also function in iron-sulfur cluster assembly alongside with SufBCD[88] (arCOG01715, arCOG04236, TIGR01981), which are encoded by 10 out of 12 MAGs. Additionally, Undinarchaeota MAGs encode a potential serine-pyruvate aminotransferase (PucG, arCOG00082), which might transaminate L-serine and pyruvate to 3-hydroxypyruvate and alanine. While Undinarchaeota MAGs encode a 3-phosphoglycerate dehydrogenase (SerA, arCOG01754), which catalyzes the first step in serine biosynthesis, genes required to mediate the other two steps of this pathway, generally encoded by *ser*B (arCOG00083) and *ser*C (arCOG01158), were absent. However, it seems possible that an uncharacterized aminotransferase might complement the function of SerC. For example, in *Methanocaldococcus jannaschii* a broad-spectrum class V aminotransferase is sufficient for phosphoserine production[89]. The gene in the cited study belongs to arCOG00082, a homolog of which is present in Undinarchaeota. Interestingly, this protein has an aminotransferase class V domain (IPR000192) and thus might indeed be involved in serine biosynthesis in Undinarchaeota. Similarly, the function of phosphoserine phosphatase SerB might be mediated by another, so far uncharacterized phosphatase.

Altogether, the lack of many genes involved in amino acid metabolism suggests that Undinarchaeota representatives need to acquire amino acids from the environment or a host, for instance using the amino acid transporter PotE (arCOG00009) that is present in all MAGs (**Supplementary Table 7**). PotE has an amino acid/polyamine transporter domain (IPR002293), but lacks any additional domain that would allow to make more specific predictions regarding the identity of amino acids that could be taken up by this transporter. The lack of other amino acid transport systems could suggest that Undinarchaeota representatives require a host to obtain certain amino acids and other necessary metabolites (see also below) directly.

##### Lipid biosynthesis

Previous analyses have revealed that many DPANN archaea lack lipid biosynthesis genes[74,90] and it was shown that several cultivated representatives such as *N. equitans* and potentially *Nanohaloarchaeaum antarcticus*, acquire their lipids from their respective hosts[91–93]. Interestingly, especially aquifer Naiadarchaeales MAGs encode for a far more complete gene set for lipid biosynthesis than *N. equitans* and many other DPANN archaea (**Fig. 3**, **Supplementary Tables 7-9 and 12, Supplementary Figure S62**). First of all, all Undinarchaeota MAGs encode the key genes for proteins involved in the mevalonate pathway, which allows the conversion of acetyl-CoA to isopentenyl-diphosphate (IPP). These proteins include HmgB (arCOG01767), HmgA (arCOG04260), Mvk (arCOG01028), MvaD (arCOG02937, IPR029765) and Ipk (arCOG00860). Furthermore, the presence of genes for geranylgeranyl diphosphate synthase (GGPS, arCOG01726), suggests that Undinarchaeota representatives have the ability to convert IPP further to geranylgeranyl diphosphate (GGPP), a precursor of ether-linked lipids[94]. Undinarchaeota MAGs also encode an undecaprenyl-diphosphate synthase (UppS, arCOG01532) that could convert GGPP to undecaprenyl diphosphate, which is a potential precursor of glycosyl carrier lipids[95]. Intriguingly, genes for enzymes synthesizing archaeol via the glycerophospholipid pathway seem to be solely encoded by aquifer but absent from marine Undinarchaeota MAGs. For instance, aquifer Naiadarchaeales code for glycerol-1-phosphate dehydrogenase (EgsA, arCOG00982) that transforms glycerone-1-phosphate to glycerol-1-phosphate and is essential to build the backbone of phospholipids[96]. Furthermore, two aquifer Naiadarchaeales MAGs (SRR2090159.bin1129 and SRR2090159.bin1288) encode phosphoglycerol geranylgeranyltransferase (GGGPS, arCOG01085), which could convert glycerol 1-phosphate and GGPP to geranylgeranyl)glycerol 1-phosphate, catalyzing the first step in archaeal lipid biosynthesis[97]. While all Undinarchaeota representatives encode a protein assigned to the arCOG00476 family comprising putative digeranylgeranylglyceryl phosphate synthases (DGGGP synthase), only the homologs identified in the aquifer MAGs harbor the characteristic DGGGP synthase domain (IPR023547). Finally, all aquifer MAGs encode enzymes for the last steps of the archaeal lipid biosynthesis: these include CDP-archaeol synthase (CarS, arCOG04106)[98], as well as archaetidylinositol phosphate synthase (AIP synthase, arCOG00670), archaetidylserine synthase (AS synthase, arCOG00671) and the putative archaetidylserine decarboxylase (Psd, arCOG04470)[99]. Thus, while aquifer Naiadarchaeales seem to be able to synthesize their own lipids, marine Undinarchaeales representatives instead may rely on lipids or certain intermediates from potential interaction partners.

##### Vitamin and cofactor biosynthesis

Undinarchaeota MAGs encode diverse enzymes whose function is dependent on the presence of vitamins and cofactors, such as the thiamine-domain (IPR029061) containing proteins, such as the pyruvate dehydrogenase (PdhA, arCOG01054) and transketolase (Tk, arCOG01051 and arCOG01053). Yet, Undinarchaeota MAGs seem to lack most genes coding for enzymes involved in vitamin biosynthesis pathways (**Supplementary Tables 7-9 and 12**). The few enzymes present include the nicotinamide-nucleotide adenylyltransferase (NadR, arCOG00972) encoded by all aquifer Naiadarchaeales MAGs and dihydrofolate reductase (FolA, arCOG01490) found in marine Undinarchaeales representatives. Additionally, most Undinarchaeota representatives contain genes coding for a putative dephospho-CoA kinase (CoaE, arCOG01045) and a protein assigned to the arCOG04076 family, which comprises candidate enzymes for GTP-dependent dephospho-CoA kinases[100]. However, we could not detect genes for other enzymes involved in coenzyme A biosynthesis in any of the Undinarchaeota genomes such as coenzyme A biosynthesis bifunctional protein CoaB (arCOG01704) or phosphopantetheine adenylyltransferase (CoaD, arCOG01223). Additionally, Undinarchaeota MAGs seem to lack genes for transporters specific to coenzymes and vitamins, which would allow the uptake of these compounds (**Supplementary Table 13**). In turn, this further suggests that the herein analyzed Undinarchaeota representatives may depend on direct contact with a partner organism to acquire vitamins and cofactors.

#### Host-symbiont interactions

Comparative genome analyses revealed a limited set of central carbon metabolism related proteins as well as the low number of genes encoding transporters and enzymes involved in vitamin and amino acid biosynthesis, raising the possibility that Undinarchaeota representatives depend on partner organisms for growth. To shed more light onto potential interaction partners, we have analyzed proteins that may be involved in species-species interactions and inferred routes of horizontal gene transfer and generated proportionality networks (see Main Text).

##### Genes potentially involved in host-symbiont interactions

Cellular appendages, such as pili and the archaellum, and other surface proteins (i.e. LamG-domain containing proteins) represent mechanisms reported to mediate cell-cell interactions[4,101]. While we did not detect genes encoding subunits of the archaellum[102], it has previously been suggested that certain DPANN use pili to interact with their hosts[103]. Undinarchaeota MAGs have gene clusters encoding several proteins potentially involved in pili formation (VirB11; arCOG01818, TadC; arCOG01808, EppA;

arCOG02300) as well as an uncharacterized protein with archaeal pilin domains (i.e. arCOG03871/IPR013373) (**Supplementary Tables 7-9**). The potential VirB11 protein contains a P-loop NTPase domain (IPR027417) and might generate energy from NTP hydrolysis[104] and TadC might provide an assembly platform for the assembly of pili[105]. Prepilins, which are encoded by 10 out of 12 Undinarchaeota MAGs, could be transported through the membrane via the sec-transport system (secDEFGY encoded by arCOG03055, arCOG02204, arCOG03054, arCOG02957 and arCOG04169) and modified via potential prepilin peptidases (arCOG02298, arCOG02300 and arCOG02300). Some pili-related proteins seem to be present in aquifer Naiadarchaeales representatives but are not encoded by marine Undinarchaeales MAGs: these include CpaF (an uncharacterized protein with a type II/IV secretion system protein domain: arCOG01819; IPR001482) and uncharacterized proteins in the same genetic region that may functionally be related to pili formation such as a potential surface binding proteins (arCOG05787, arCOG03512). A potential VirB4 ATPase (arCOG04035) on the other hand is only encoded by marine Undinarchaeales MAGs.

Surface modification proteins, involved in the modification of S-layers and construction of extracellular matrices, represent additional means that may enable host-symbiont interactions[106]. While Undinarchaeota MAGs seem to lack S-layer proteins SlaA (arCOG06039) and SlaB (arCOG07272), they encode an uncharacterized S-layer protein (arCOG03418; IPR006454) as well archaeal glycosylation proteins, such as the glycosyltransferase AglA (arCOG01410) and the protein glycotransferase AglB (arCOG02044), that might be involved in S-layer protein N-glycosylation[107]. AglA and B are encoded in the same genetic region as other archaeal glycosylation proteins (arCOG00899, arCOG03199) and potential membrane-binding proteins (arCOG05092, arCOG00395, arCOG07813 and arCOG02080). Overall, this suggests the presence of an S-layer or the potential of Undinarchaeota to generate an extracellular matrix that might play a role in cell-cell interactions. In support of the latter, we found that some of the longest proteins (~1400 amino acids) present in marine but not aquifer Undinarchaeota representatives encode LamG-like protein domains (arCOG07813; IPR006558), which might be involved in the formation of an extracellular matrix (**Supplementary Table S20-23**)[108]. LamG-like proteins in Undinarchaeota are often encoded in the same genetic region as potential S-layer proteins, glycosyltransferases (AglA) or pilus-assembly proteins (TadA). Furthermore, our investigation of a suite of proteins discussed to be involved in cell-interactions[74] and present in the Undinarchaeota MAGs shares similarity to proteins involved in cell adhesion( **Supplementary Table 22-23**). We also identified a hypothetical protein with TSP type-3 repeat domains (IPR028974, arCOG07561) in marine Undinarchaeales MAGs, which may represent another putative extracellular matrix protein. Finally, it is worth mentioning that similar to many other DPANN archaea[74,109], Undinarchaeota MAGs do not seem to encode CRISPR-Cas systems, which, among others, are involved in viral defense[110].

#### Horizontal gene transfer among Undinarchaeota and other microbial lineages

It has previously been shown that intimately interacting organisms can share genes through horizontal gene transfer (HGT). For example, *N. equitans,* the first cultivated representative of the DPANN, and its host *I. hospitalis* seem to have exchanged several genes horizontally[111,112]. To investigate the possibility whether Undinarchaeota representatives have exchanged genes with potential hosts and to pinpoint routes of HGT, we reconstructed protein trees of all proteins present in at least three or more

Undinarchaeota genomes (520 genes total) and analyzed sisterhood-relationships among taxonomically distinct lineages. In brief, we identified homologs of these 520 Undinarchaeota protein families in a reference set of 364 archaeal, 3020 bacterial and 100 eukaryotic genomes and generated single protein trees. Subsequently, HGT events were identified using a custom script (count_sister_taxa.py; https://github.com/Tancata/phylo/blob/master/count_sister_taxa.py), that allows to determine the next closest sister lineage of any lineage of interest (see Methods for details). Notably, this approach revealed significant fractions of HGT among known DPANN symbiont-host systems (**Fig. 4a, b**, **Supplementary Tables 16-17**). Undinarchaeota did not show a dominant fraction of genes shared with a specific lineage, i.e. most genes seemed to be shared with taxonomically closely related DPANN Archaea. The largest number of proteins, that did not cluster with DPANN homologs, seemed to be related to homologs of Asgard archaea (16 proteins to Heimdall-, Loki- or Thorarchaeota), Bathyarchaeota (11 proteins), and Thermoplasmata (11 proteins to Thermoplasmatales, Pontarchaea or Poseidoniales) (**Supplementary Tables 16-17**). Notably, the protein families potentially transferred with these members of these archaeal lineage comprise components of informational processing machineries such as a potential tRNA pseudouridine synthase (arCOG04252; one Bathyarchaeota clustering inside Undinarchaeota), RNA 3'-terminal phosphate cyclase (arCOG04125, transfer to Heimdallarchaeota) and ribosomal protein S19 (arCOG04099; transfer to Pontarchaea). Considering that genes for information processing are thought to evolve predominantly vertically but may be exchanged between known symbiont-hosts systems, this opens the possibility that marine Undinarchaeales engage in symbiotic interactions with one of these lineages.

#### Co-occurrence analyses

Next, we used a read-based co-occurrence analysis to assess whether MAGs of Undinarchaeota are proportional to other archaeal and bacterial genomes[113] (see Methods for details). Unfortunately, we only detected Undinarchaeota in a low number of metagenome datasets, such that this analysis does not have sufficient statistical power to resolve co-proportionality with high support. In turn, we did not detect any significant co-occurrence patterns for members of the Naiadarchaeales and any other taxonomic lineage. In fact, the majority of genomes co-varying with Undinarchaeota MAGs belongs to other DPANN and Patescibacteria/CPR lineages, which are all characterized by small cell sizes and reduced genomes such that these co-occurrence patterns could be due to an artifact resulting from the enrichment of small cells for some of the samples (though interactions among members of these lineages cannot be excluded). The main observation was that marine Undinarchaeales appeared to co-vary with three genomes of the Chloroflexi, all belonging to the order Dehalococcoidales (**Supplementary Figure 63)**. Most members of the Dehalococcoidales have small genomes (i.e. ~1.5 Mb for *Dehalococcoides mccartyi* 195, which however has a rather large cell size of 0.3-1 µm) and represent free-living heterotrophic bacteria that can use chlorinated compounds as electron acceptors[114]. While challenging to grow in isolation, Dehalococcoides can be maintained in enrichment cultures, in which they rely on acetate and hydrogen from other community members[114]. Based on metabolic gene repertoires of members of the Undinarchaeales, which are characterized by the absence of any of the known genes for the various hydrogenase protein families[115] (**Supplementary Table 7**), hydrogen-dependent syntrophy seems unlikely to support interactions with Dehalococcoides. Yet, a symbiotic relationship could be based on exchange of acetate or be of parasitic nature as observed in currently known host-symbiont systems[93,116–119]. It has to be noted, however, that the UAP2-positive metagenomes (**see Supplementary Table S1**) used for the proportionality analyses differ in respect to sampling method and filtering steps and we cannot exclude that correlation patterns are due to methodological artefacts. Therefore, prospective analyses of a larger number of metagenomes generated with consistent methodology and without filtering steps will be needed to further assess co-proportionality of Undinarchaeota with other organism groups.

Yet, since co-occurrence analyses predicted a potential association of marine Undinarchaeales with three Chloroflexi of the order Dehalococcoidales (**Supplementary Figure 63**) we manually investigated single-gene trees for potential transfers between these groups but could only identify a small fraction of candidate HGTs. For example, a potentially transferred gene encodes a Fe-S cluster assembly ATPase SufC (arCOG04236). In the corresponding phylogeny, two Chloroflexi (*Thermogemmatispora carboxidivorans* and *Ktedonobacter* sp.) branch as a sister group of marine Undinarchaeales with low bootstrap support of 41%. Another potential transfer involves a gene for mevalonate kinase (arCOG01028). In particular, the undinarchaeal sequence from GCA_002502135 emerges from within a cluster of Chloroflexi (70% bootstrap support). However, the number of putative HGTs among Chloroflexi and Undinarchaeota are significantly lower than the HGTs detected in other DPANN-host symbiont systems, such as Nanoarchaeota and Crenarchaeota, and do therefore not provide support for the association Undinarchaeota with members of this bacterial lineage.

In turn, further analyses including fluorescence *in situ* hybridization will be needed to shed further light onto potential interaction partners of Undinarchaeota and test whether certain members of this group indeed interact with members of the Pontarchaea or Chloroflexi.

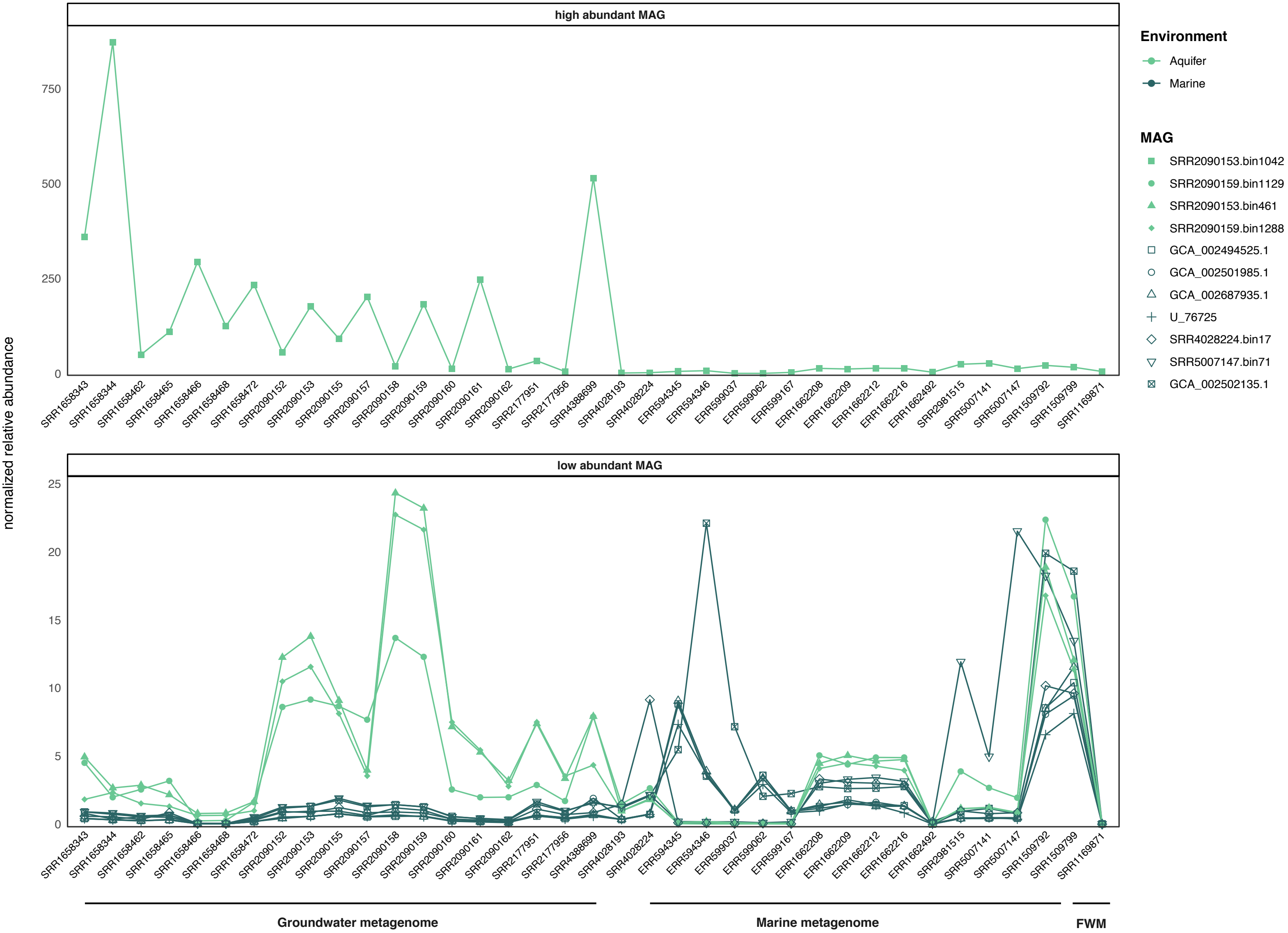


**Supplementary Figure 1 | Abundance of Undinarchaota (UAP2) across 37 metagenomic datasets.** Normalized relative abundance of aquifer Naiadarchaeales (light green) and marine Undinarchaeales (dark green) across 37 metagenomes. Relative abundances were normalized by the total read counts. Read mapping was done to metagenomes belonging to different environmental types including groundwater, marine and freshwater metagenomes (FWM). Due to the difference in abundance, MAGs with a relative abundance >50 were plotted separately. Details on the metagenomes can be found in Supplementary Table S1.

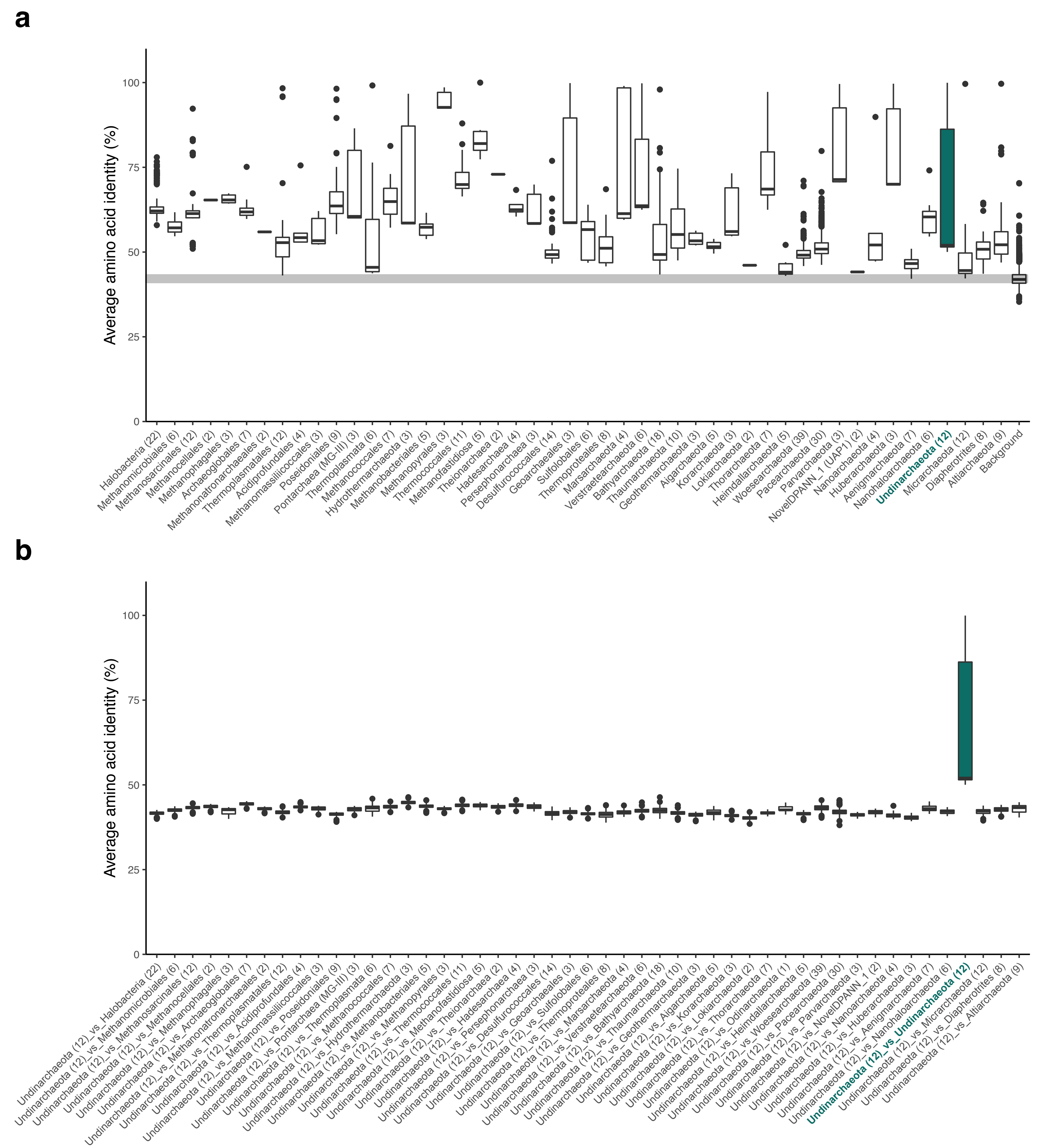


**Supplementary Figure 2 | Comparing the amino acid identity (AAI) of major archaeal lineages**. **a**, Shared AAI across archaeal lineages. Background: Comparing all archaeal lineages included in the analyses but excluding archaea belonging to the same lineages in order to determine the lowest AAI that defines a cluster. **b**, AAI of Undinarchaeota compared to all other archaeal lineages to show that the highest identity is when comparing Undinarchaeota to themselves. The lower and upper hinges of the boxplot correspond to the first and third quartiles. The upper/lower whiskers extend from the hinge to the largest/smallest value no further than 1.5* of the inter-quartile range. Data beyond the whiskers are shown as individual data points. Number in brackets: Number of genomes included in each cluster. Raw values are listed in Supplementary Table 3.

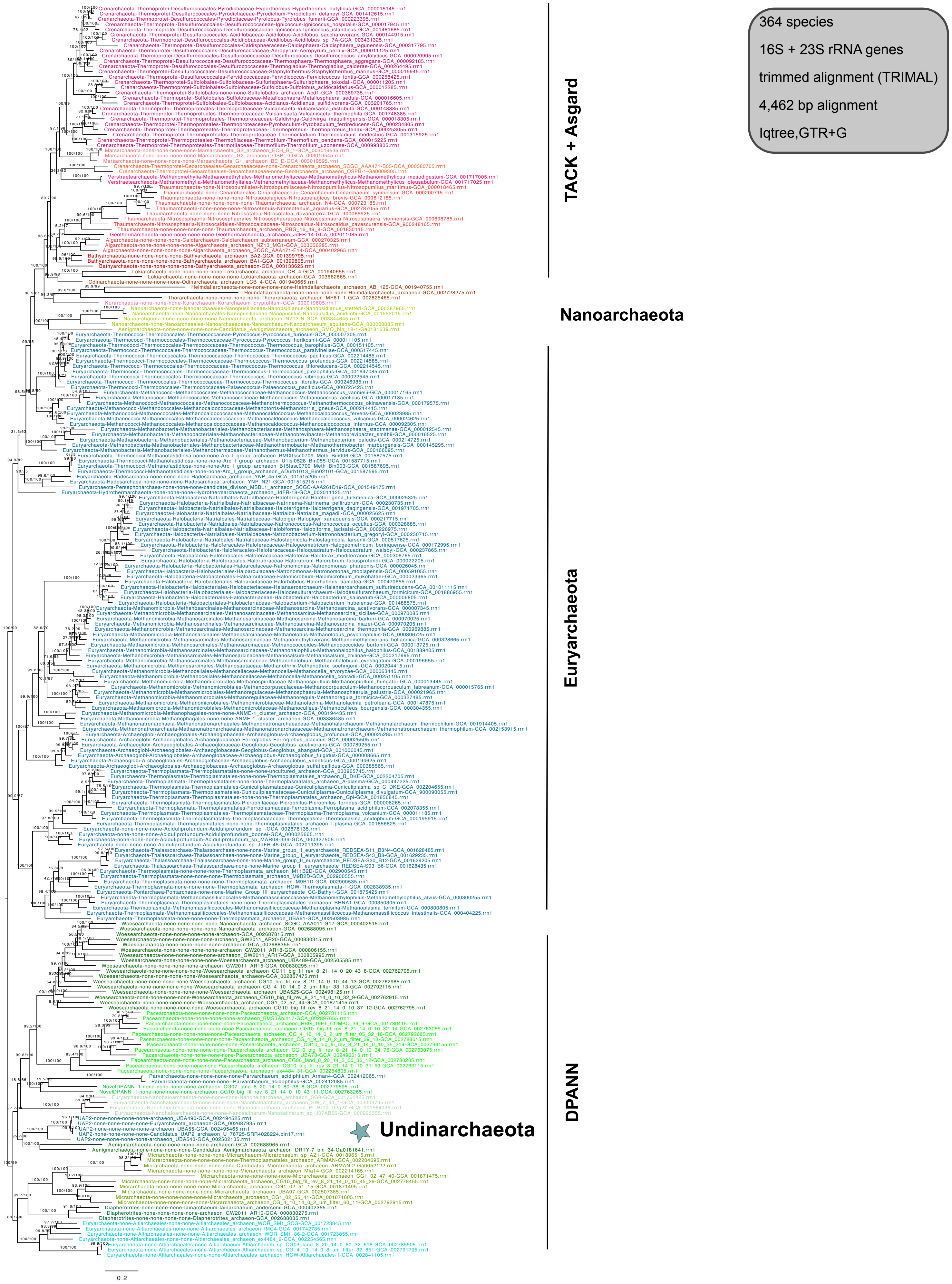


**Supplementary Figure 3 | Phylogenetic placement of Undinarchaeota based on a concatenated alignment of the 16S and 23S rRNA gene sequences.** Sequences were extracted from the 364 species set (of these 238 species encoded 16S and/or 23 rRNA genes). The alignment was trimmed using TRIMAL (alignment length = 4,462 bp) and a maximum-likelihood phylogenetic tree was inferred with the GTR+G model with an ultrafast bootstrap approximation (left) and SH-like approximate likelihood tests (right), each run with 1000 replicates. Scale bar: Average number of substitutions per site. The tree was artificially rooted using DPANN archaea.

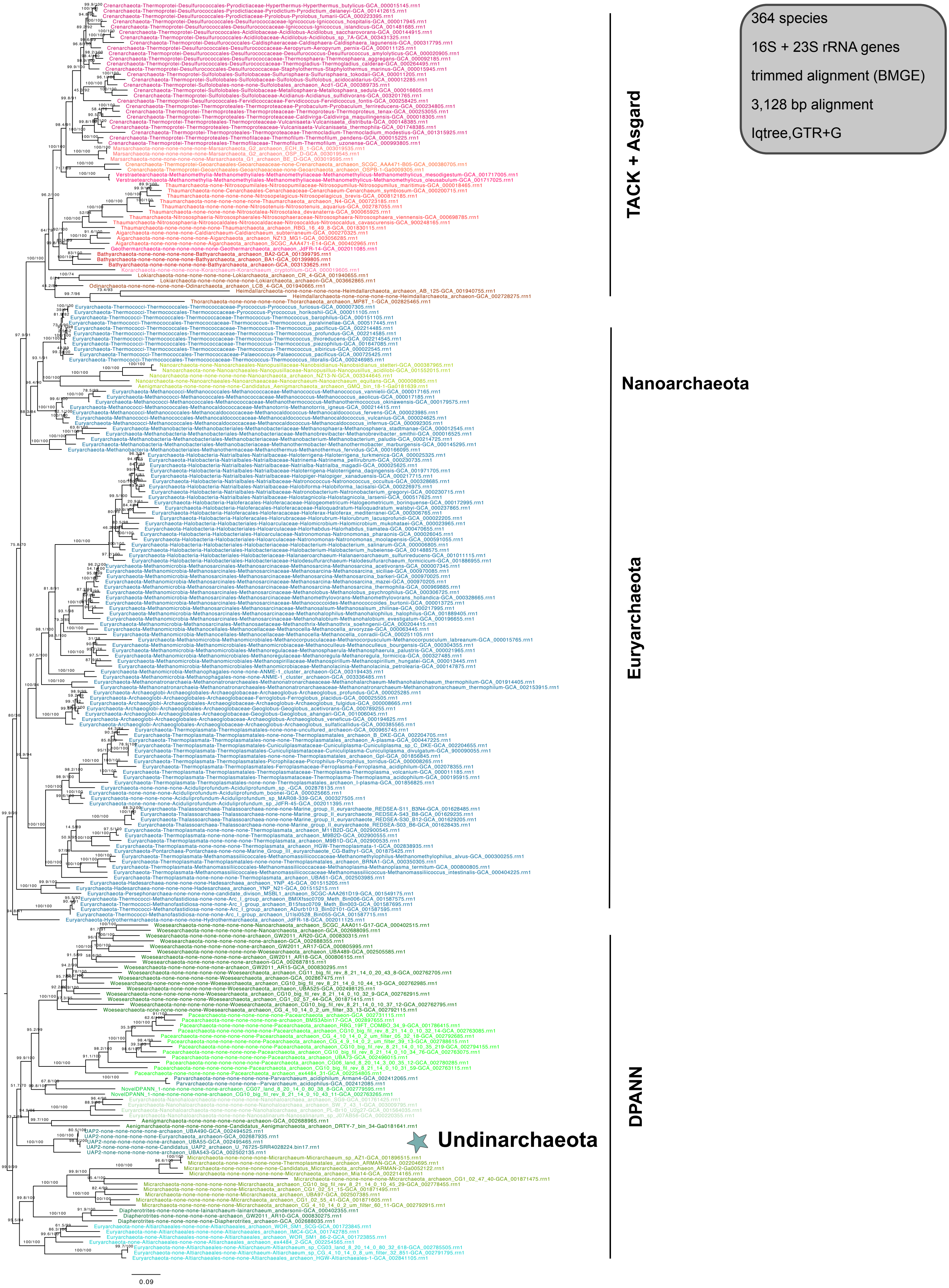


**Supplementary Figure 4 | Phylogenetic placement of Undinarchaeota based on a concatenated alignment of the 16S and 23S rRNA gene sequences**. Sequences were extracted from the 364 species set (of these 238 species encoded 16S and/or 23 rRNA genes). The alignment was trimmed using BMGE (alignment length = 3,128 bp) and a maximum-likelihood phylogenetic tree was inferred with the GTR+G model with an ultrafast bootstrap approximation (left) and SH-like approximate likelihood tests (right), each run with 1000 replicates. Scale bar: Average number of substitutions per site. The tree was artificially rooted using DPANN archaea.

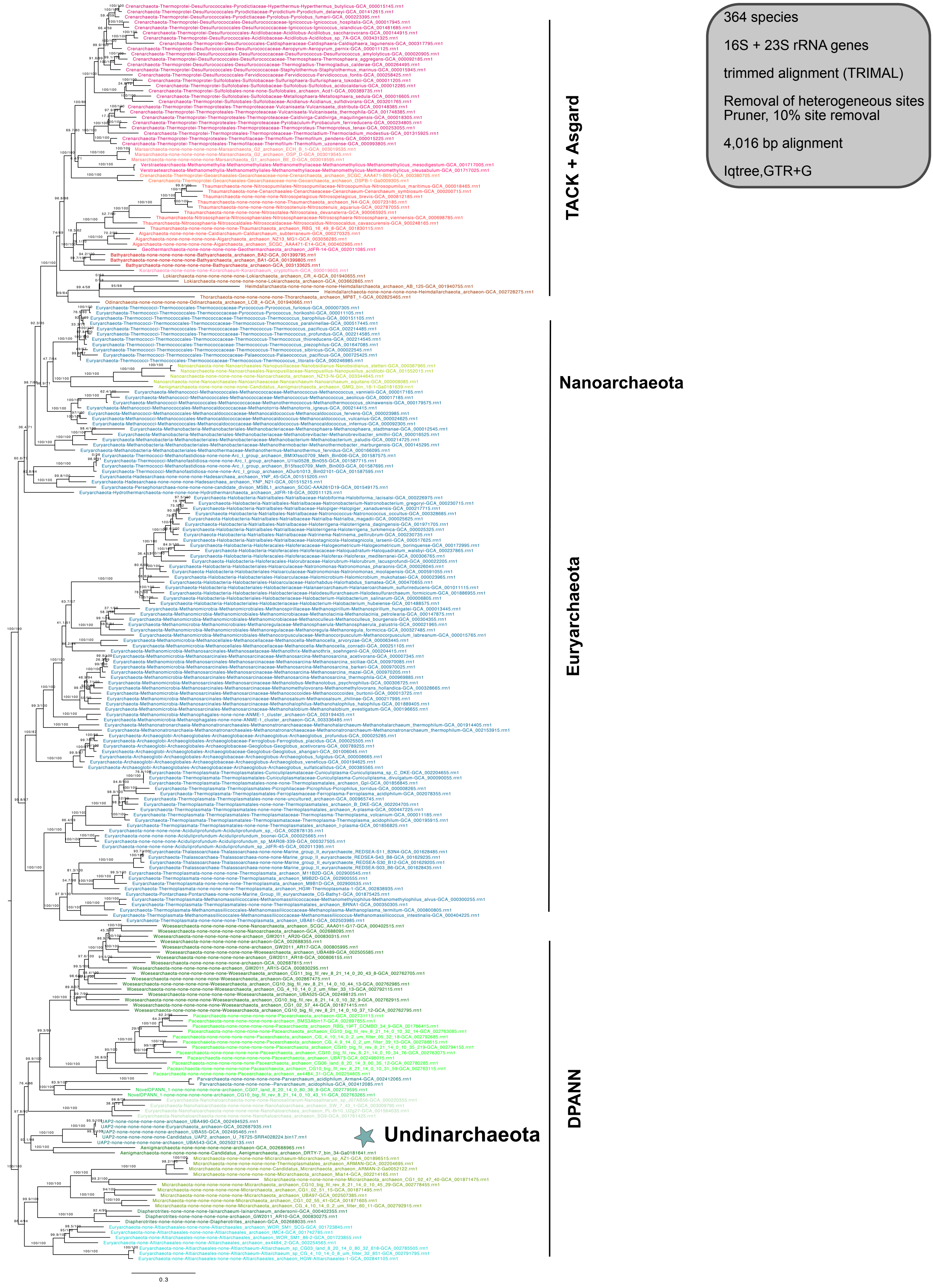


**Supplementary Figure 5 | Phylogenetic placement of Undinarchaeota based on a concatenated alignment of the 16S and 23S rRNA gene sequences**. Sequences were extracted from the 364 species set (of these 238 species encoded 16S and/or 23 rRNA genes). The alignment was trimmed using TRIMAL and 10% of the most heterogenous sites were removed using an alignment pruner (alignment length = 4,016 bp). A maximum likelihood phylogenetic tree was inferred with the GTR+G model with an ultrafast bootstrap approximation (left) and SH-like approximate likelihood tests (right), each run with 1000 replicates. Scale bar: Average number of substitutions per site. The tree was artificially rooted using DPANN archaea.

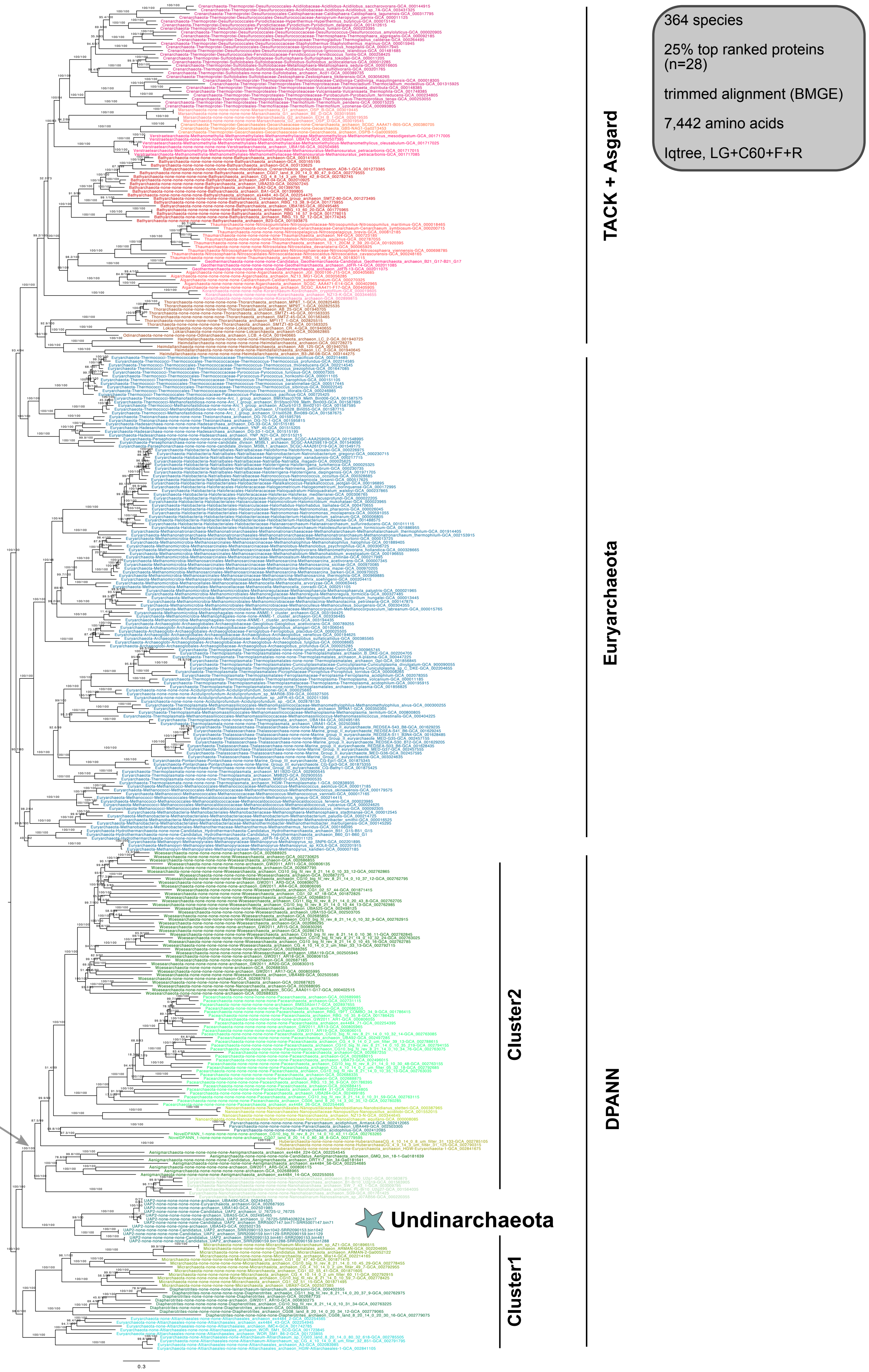


**Supplementary Figure 6 | Phylogenetic placement of Undinarchaeota based on an alignment generated with the 25% top ranked proteins (n=28) and the 364 species set.** The alignment was trimmed with BMGE (alignment length = 7,422 aa). A ML phylogenetic tree was inferred with the LG+C60+F+R model with an ultrafast bootstrap approximation (left) and SH-like approximate likelihood tests (right), each run with 1000 replicates. The tree was artificially rooted with DPANN archaea and the grey arrow shows the root position inferred with minimal ancestor deviation rooting (Tria et al., 2017). Scale bar: Average number of substitutions per site. Tree statistics for tree number 1 can be found in Supplementary Table 6.

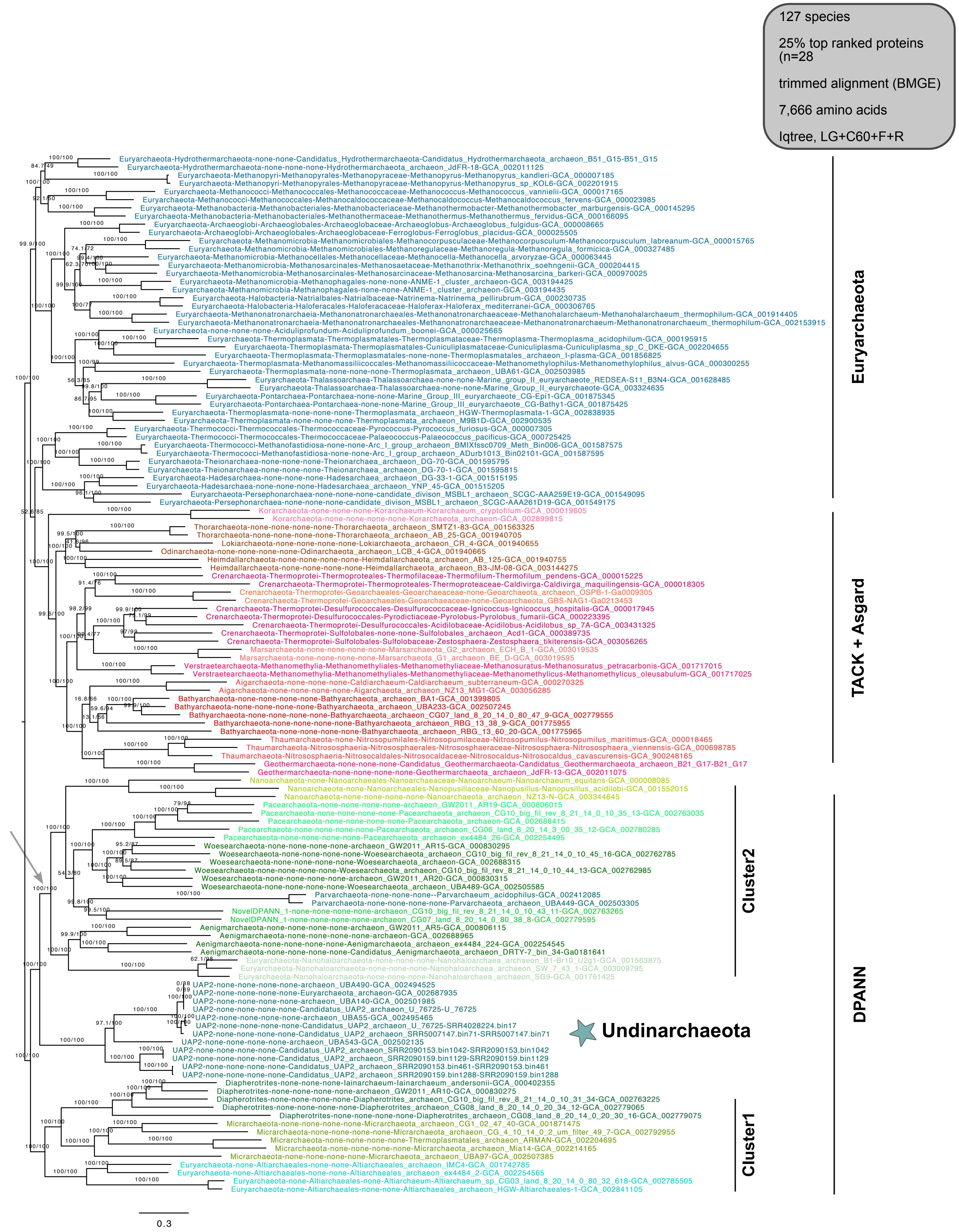


**Supplementary Figure 7 | Phylogenetic placement of Undinarchaeota based on an alignment generated with the 25% top ranked proteins (n=28) and the 127 species set.** The alignment was trimmed with BMGE (alignment length= 7,666 aa). A ML phylogenetic tree was inferred with the LG+C60+F+R model with an ultrafast bootstrap approximation (left) and SH-like approximate likelihood tests (right), each run with 1000 replicates. The tree was artificially rooted with the DPANN archaea and the grey arrow shows the root position inferred with minimal ancestor deviation rooting (Tria et al., 2017). Scale bar: Average number of substitutions per site. Tree statistics for tree number 2 can be found in Supplementary Table 6.

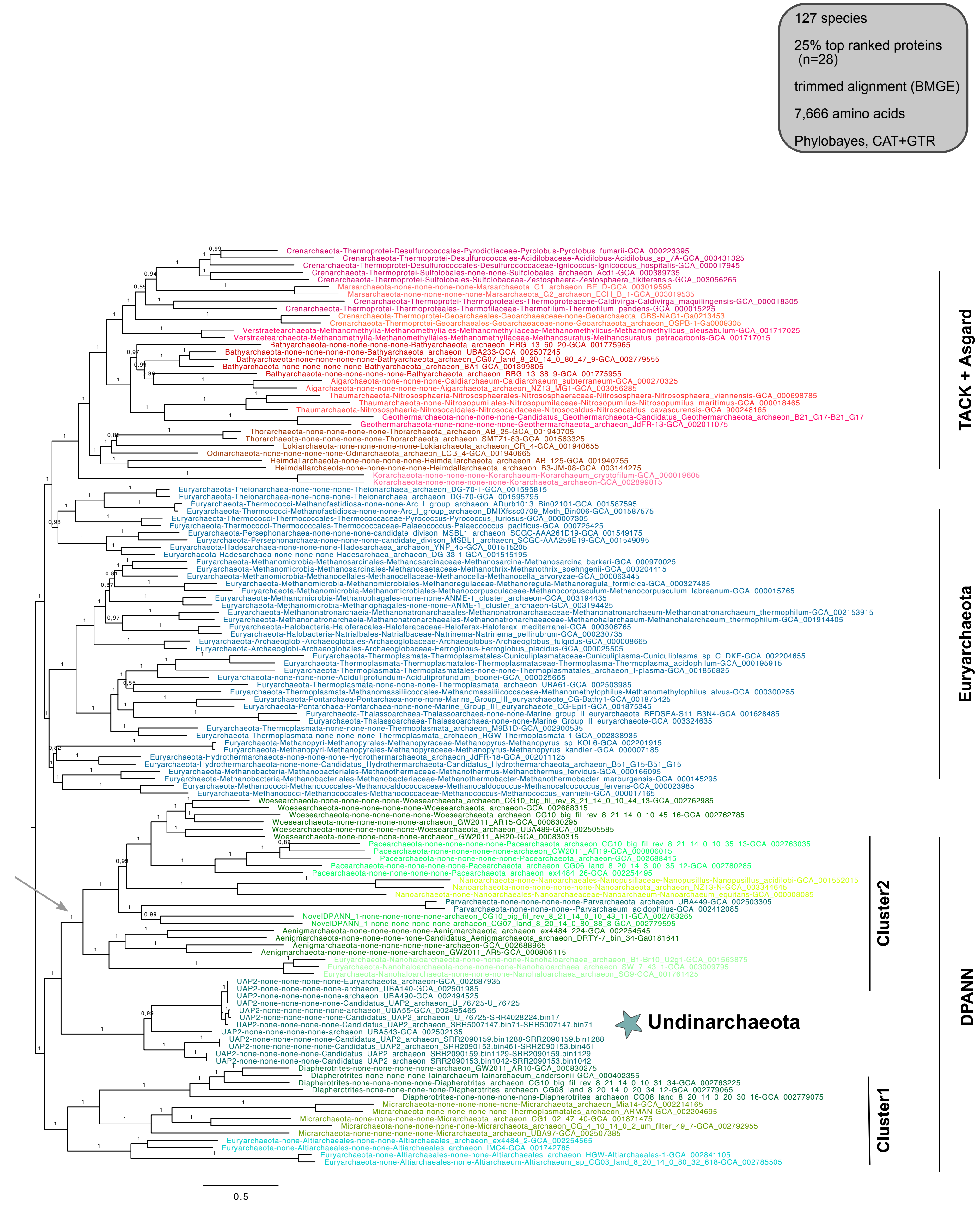


**Supplementary Figure 8 | Phylogenetic placement of Undinarchaeota based on an alignment generated with the 25% top ranked proteins (n=28) and the 127 species set.** The alignment was trimmed with BMGE (alignment length = 7,666 a). A Bayesian phylogenetic tree was inferred with the CAT+GTR model, run with four chains for a total of 34,260 cycles (25% burn-in). The tree was artificially rooted with the DPANN archaea and the grey arrow shows the root position inferred with minimal ancestor deviation rooting (Tria et al., 2017). Scale bar: Average number of substitutions per site. Tree statistics for tree number 3 can be found in Supplementary Table 6.

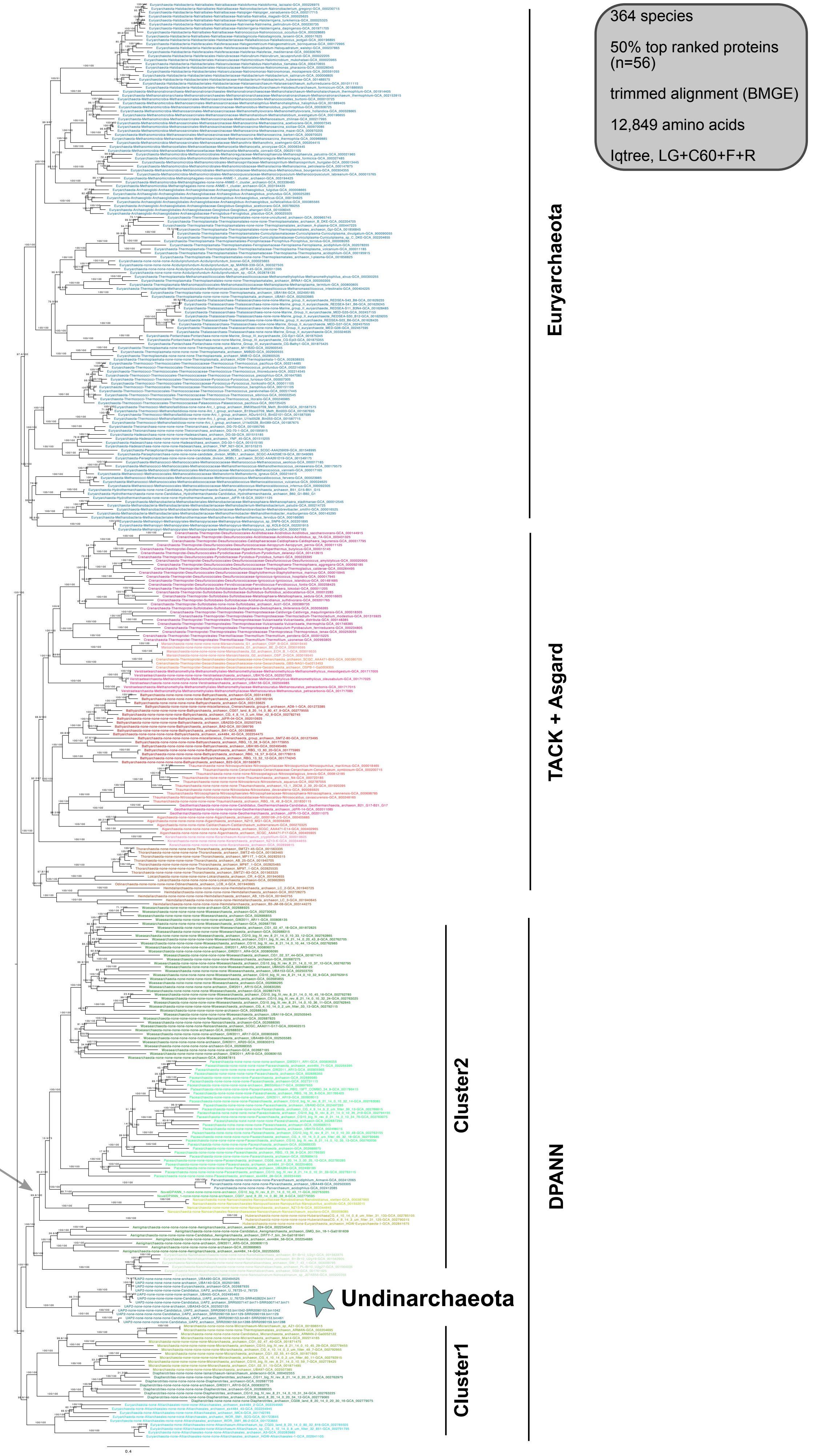


**Supplementary Figure 9 | Phylogenetic placement of Undinarchaeota based on an alignment generated with the 50% top ranked proteins (n=56) and the 364 species set**. The alignment was trimmed with BMGE (alignment length = 12,849 aa). A ML phylogenetic tree was inferred with the LG+C60+F+R model with an ultrafast bootstrap approximation (left) and SH-like approximate likelihood tests (right), each run with 1000 replicates. The tree was artificially rooted with the DPANN archaea and the grey arrow shows the root position inferred with minimal ancestor deviation rooting (Tria et al., 2017). Scale bar: Average number of substitutions per site. Tree statistics for tree number 4 can be found in Supplementary Table 6.

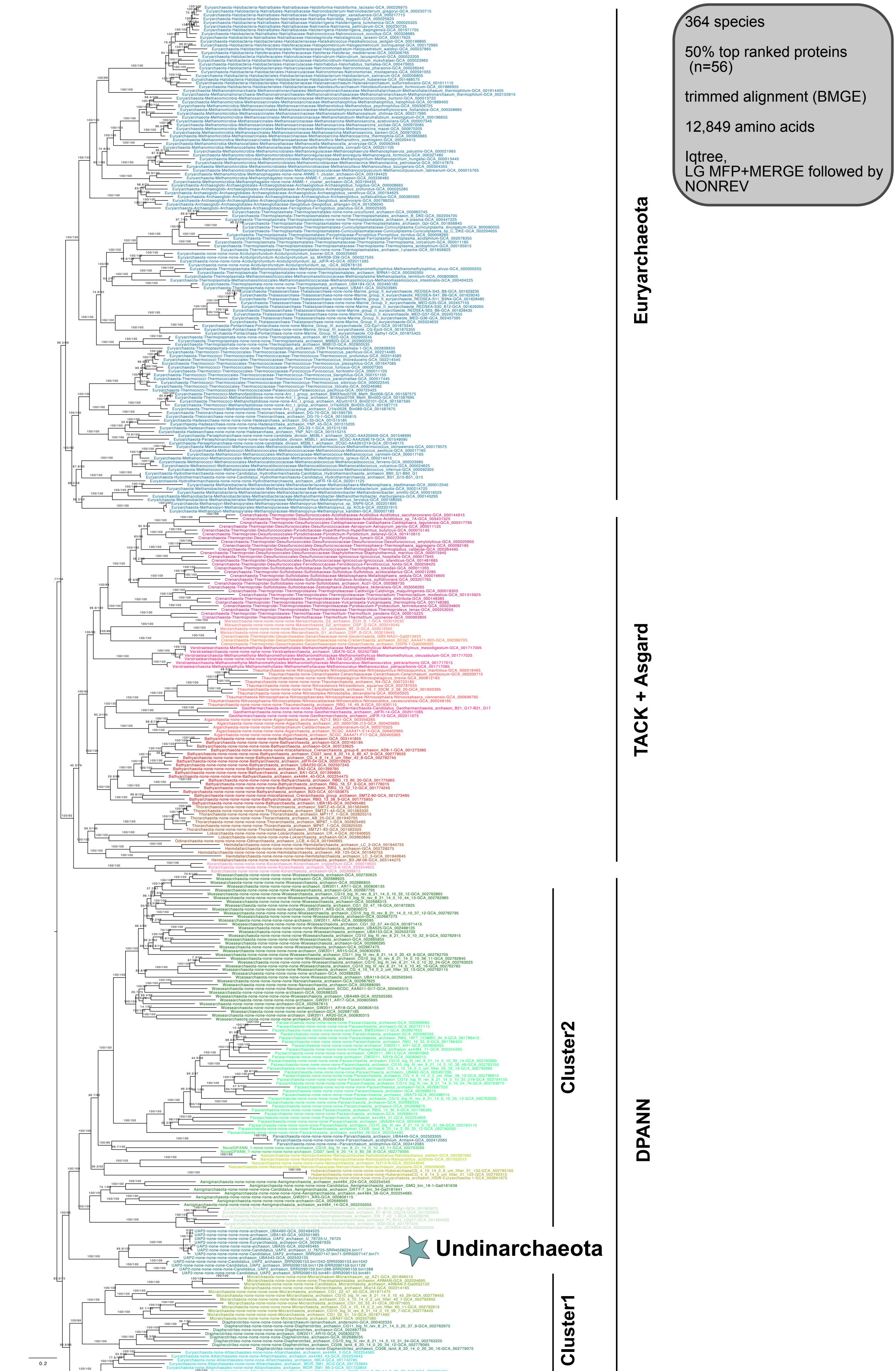


**Supplementary Figure 10 | Phylogenetic placement of Undinarchaeota based on an alignment generated with the 50% top ranked (n=56) and the 364 species set**. The alignment was trimmed with BMGE (alignment length = 12,849 aa). An initial ML phylogenetic tree was inferred with the LG model (-m MFP +MERGE) followed by a tree generated with a non-reversible model with an ultrafast bootstrap approximation (left) and SH-like approximate likelihood tests (right), each run with 1000 replicates. The tree was rooted using the non-reversible model. Scale bar: Average number of substitutions per site.Tree statistics for tree number 5 can be found in Supplementary Table 6.

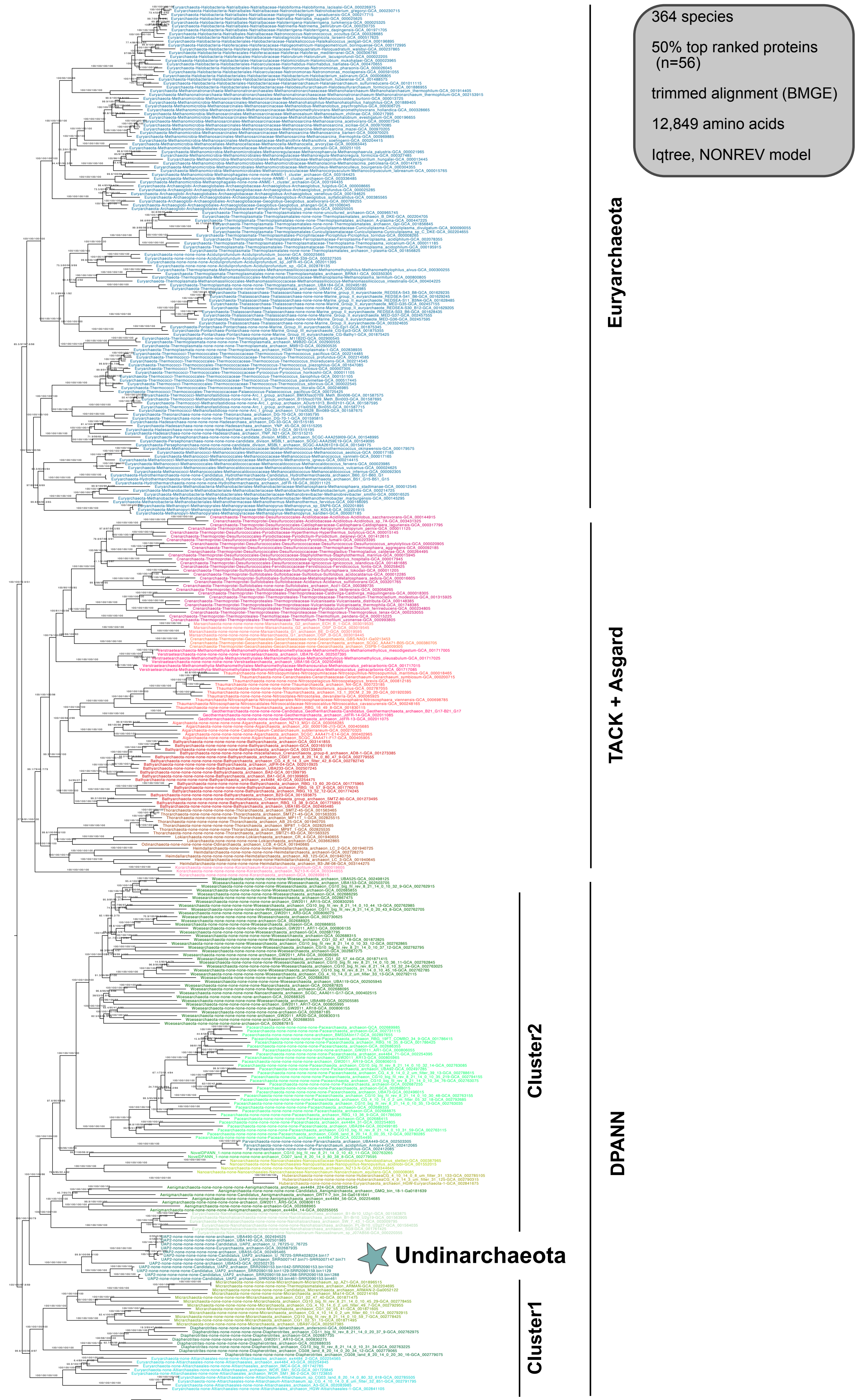


**Supplementary Figure 11 | Phylogenetic placement of Undinarchaeota based on an alignment generated with the 50% top ranked proteins (n=56) and the 364 species set**. The alignment was trimmed with BMGE (alignment length = 12,849 aa). A ML phylogenetic tree was inferred with the NONREV model. The first two values show the support for the reversible and the second two for the non-reversible model. Values 1 and 3 were generated with an ultrafast bootstrap approximation and 2 and 4 with an SH-like approximate likelihood tests, each run with 1000 replicates. The tree was rooted using the non-reversible model in iqtree v2. Scale bar: Average number of substitutions per site. Tree statistics for tree number 6 can be found in Supplementary Table 6.

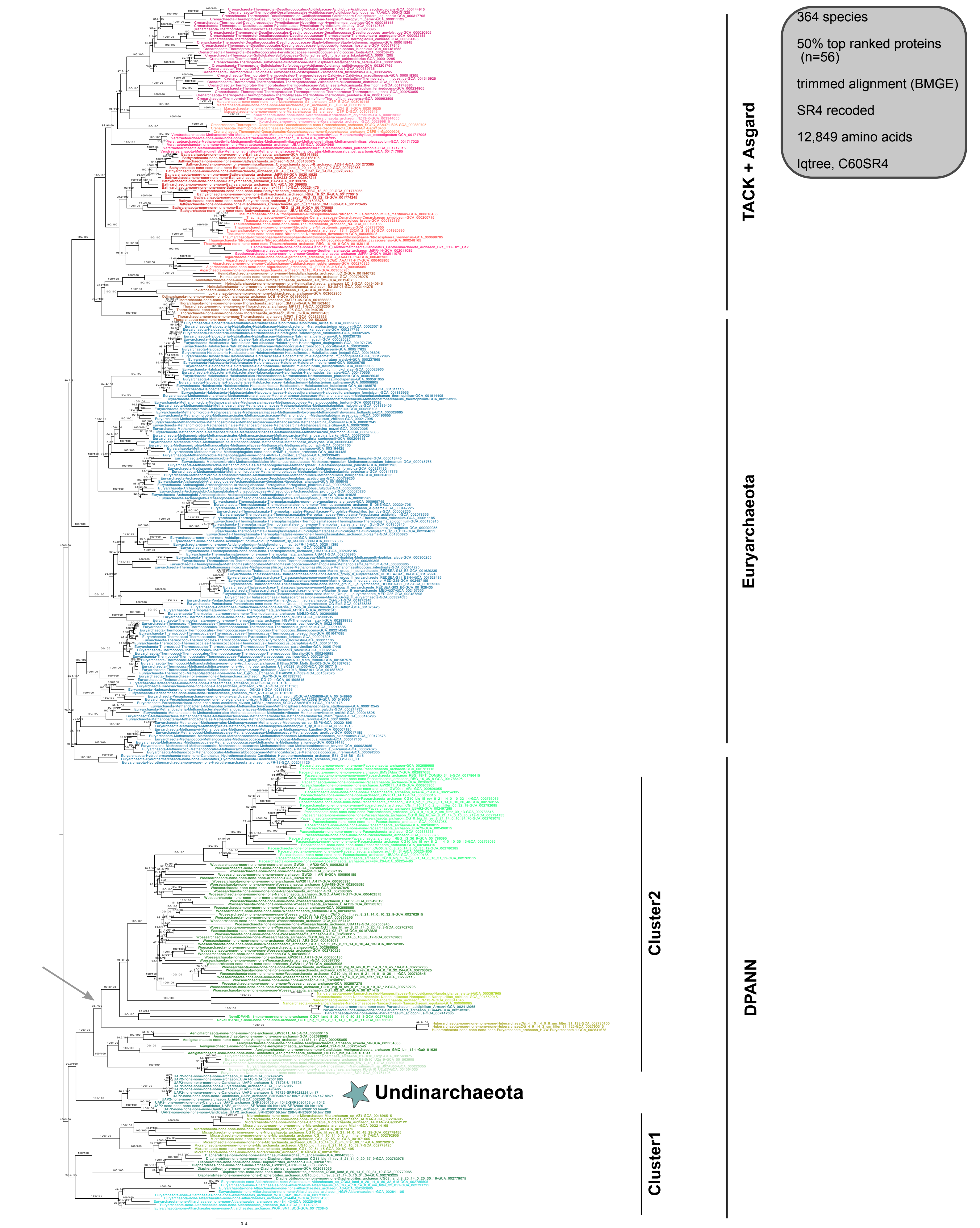


**Supplementary Figure 12 | Phylogenetic placement of Undinarchaeota based on an alignment generated with the 50% top ranked proteins (n=56) and the 364 species set.** The alignment was trimmed with BMGE and recoded into 4 character states (SR4 decoding; alignment length = 12,849 characters). A ML phylogenetic tree was inferred with the C60-SR4 model with an ultrafast bootstrap approximation (left) and SH-like approximate likelihood tests (right), each run with 1000 replicates. The tree was artificially rooted with the DPANN archaea and the grey arrow shows the root position inferred with minimal ancestor deviation rooting (Tria et al., 2017). Scale bar: Average number of substitutions per site. Tree statistics for tree number 7 can be found in Supplementary Table 6.

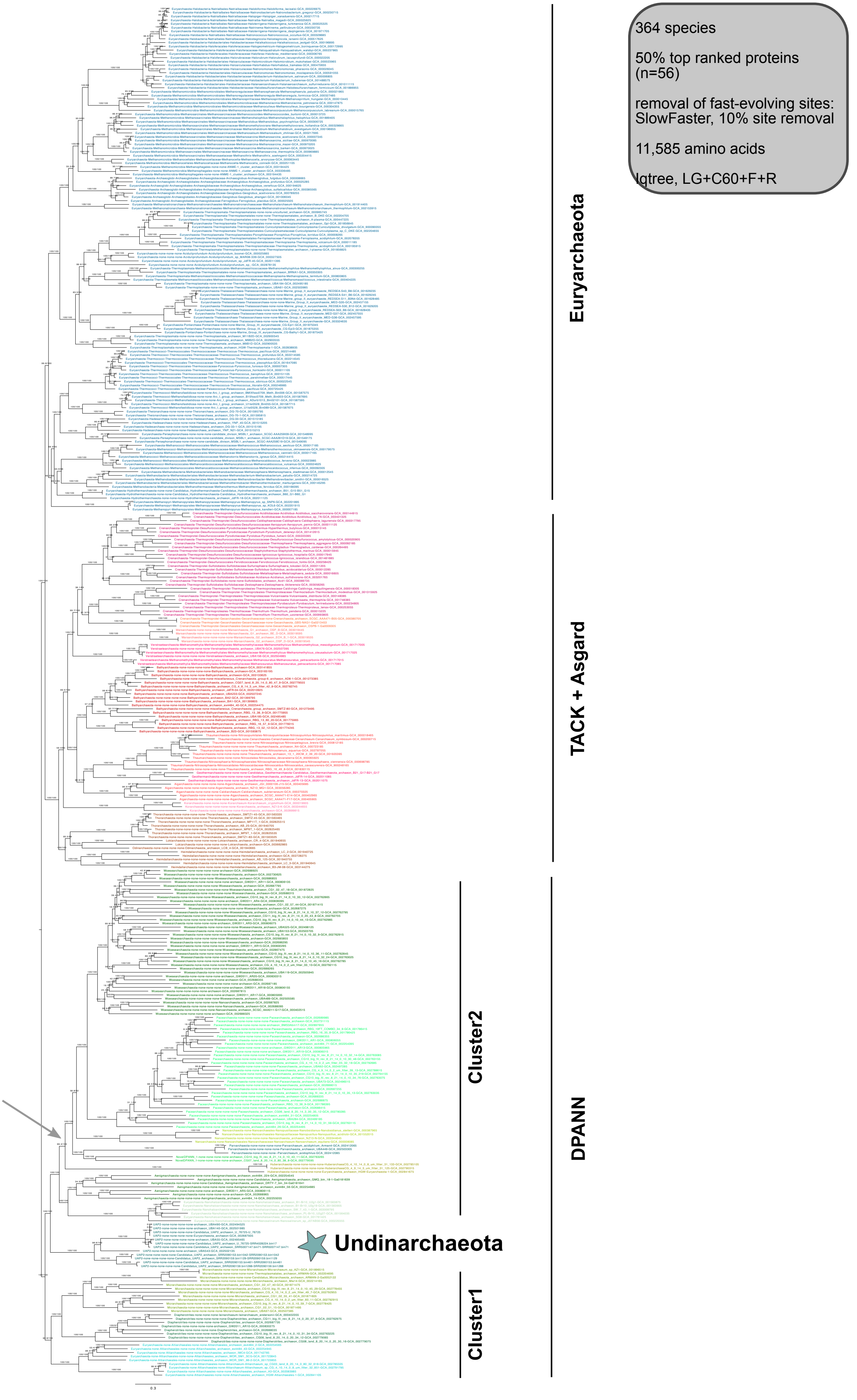


**Supplementary Figure 13 | Phylogenetic placement of Undinarchaeota based on an alignment generated with the 50% top ranked proteins (n=56) and the 364 species set.** 10% of fast-evolving sites were removed from the alignment with SlowFaster (alignment length = 11,585 aa). A ML phylogenetic tree was inferred with the LG+C60+F+R model with an ultrafast bootstrap approximation (left) and SH-like approximate likelihood tests (right), each run with 1000 replicates. The tree was artificially rooted with the DPANN archaea and the grey arrow shows the root position inferred with minimal ancestor deviation rooting (Tria et al., 2017). Scale bar: Average number of substitutions per site. Tree statistics for tree number 8 can be found in Supplementary Table 6.

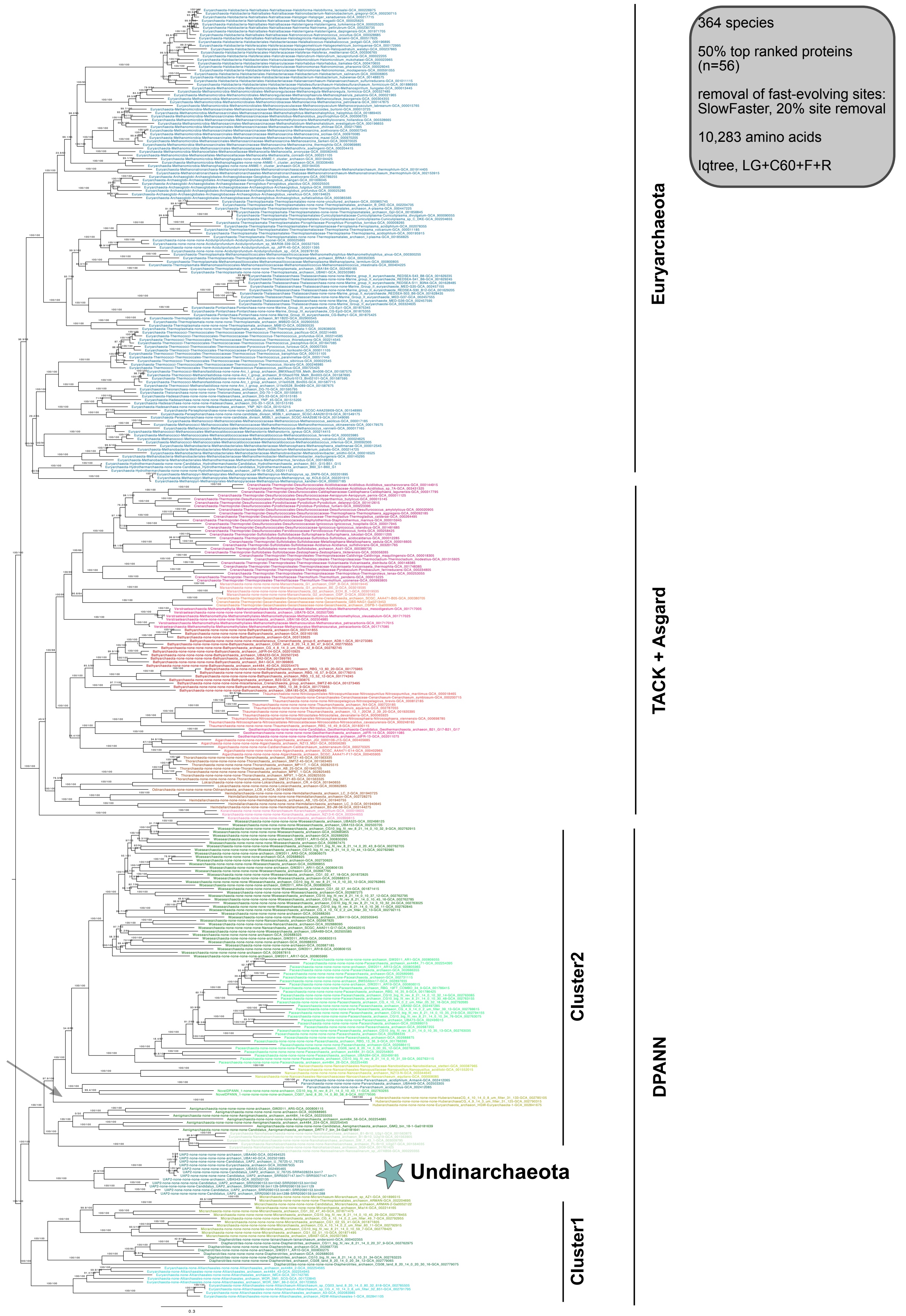


**Supplementary Figure 14 | Phylogenetic placement Undinarchaeota based on an alignment generated with the 50% top ranked proteins (n=56) and the 364 species set.** 20% of fast-evolving sites were removed from the alignment with SlowFaster (alignment length = 10,283 aa). A ML phylogenetic tree was inferred with the LG+C60+F+R model with an ultrafast bootstrap approximation (left) and SH-like approximate likelihood tests (right), each run with 1000 replicates. The tree was artificially rooted with the DPANN archaea and the grey arrow shows the root inferred with minimal ancestor deviation (Tria et al., 2017). Scale bar: Average number of substitutions per site. Tree statistics for tree number 9 can be found in Supplementary Table 6.

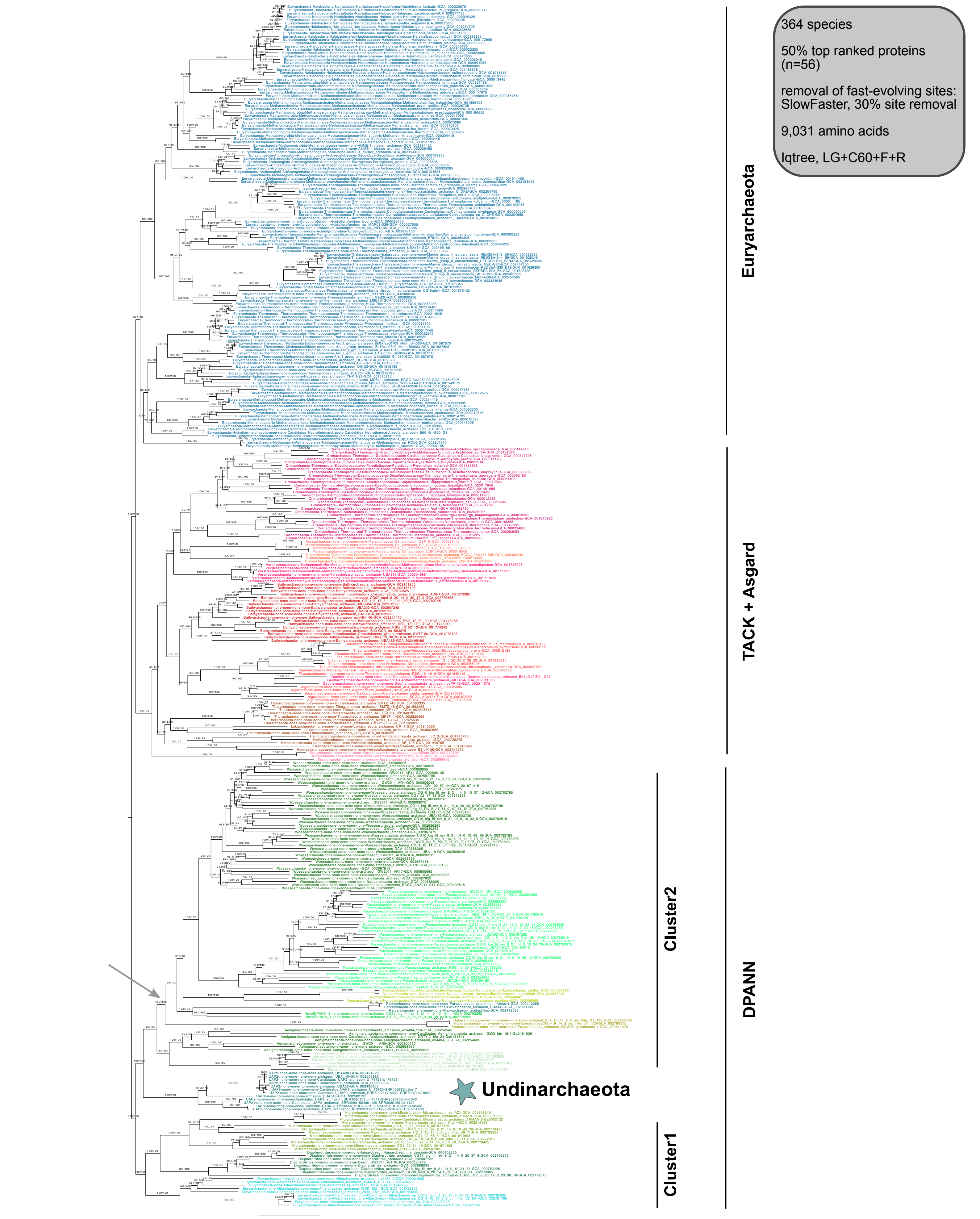


**Supplementary Figure 15 | Phylogenetic placement of Undinarchaeota based on an alignment generated with the 50% top ranked proteins (n=56) and the 364 species set.** 30% of fast-evolving sites were removed from the alignment with SlowFaster (alignment length = 9,031 aa). A ML phylogenetic tree was inferred with the LG+C60+F+R model with an ultrafast bootstrap approximation (left) and SH-like approximate likelihood tests (right), each run with 1000 replicates. The tree was artificially rooted with the DPANN archaea and the grey arrow shows the root position inferred with minimal ancestor deviation (Tria et al., 2017). Scale bar: Average number of substitutions per site. Tree statistics for tree number 10 can be found in Supplementary Table 6.

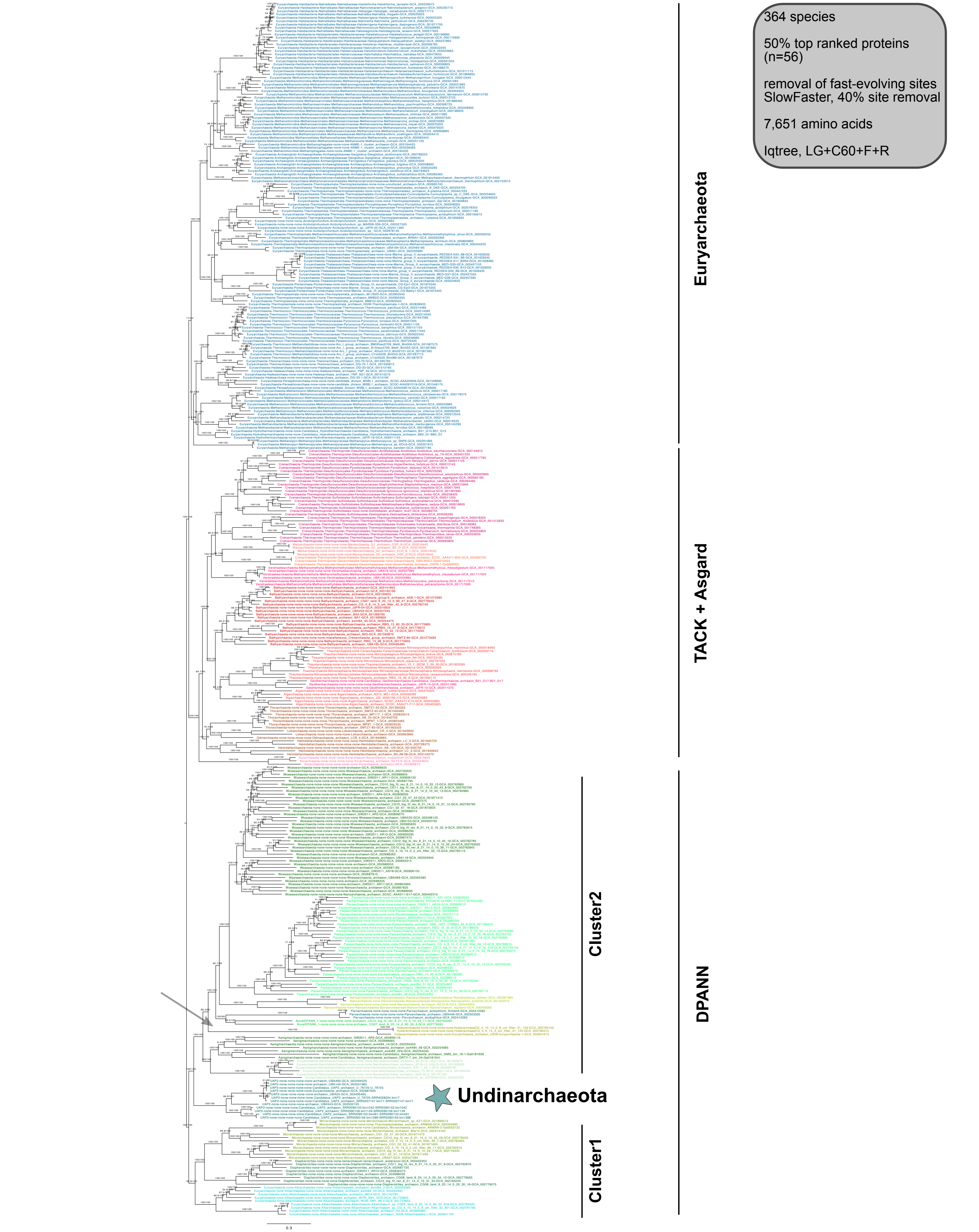


**Supplementary Figure 16 | Phylogenetic placement of Undinarchaeota based on an alignment generated with the 50% top ranked proteins (n=56) and the 364 species set.** 40% of fast-evolving sites were removed from the alignment with SlowFaster (alignment length = 7,651 aa). A ML phylogenetic tree was inferred with the LG+C60+F+R model with an ultrafast bootstrap approximation (left) and SH-like approximate likelihood tests (right), each run with 1000 replicates. The tree was artificially rooted with the DPANN archaea and the grey arrow shows the root position inferred with minimal ancestor deviation rooting (Tria et al., 2017). Scale bar: Average number of substitutions per site. Tree statistics for tree number 11 can be found in Supplementary Table 6.

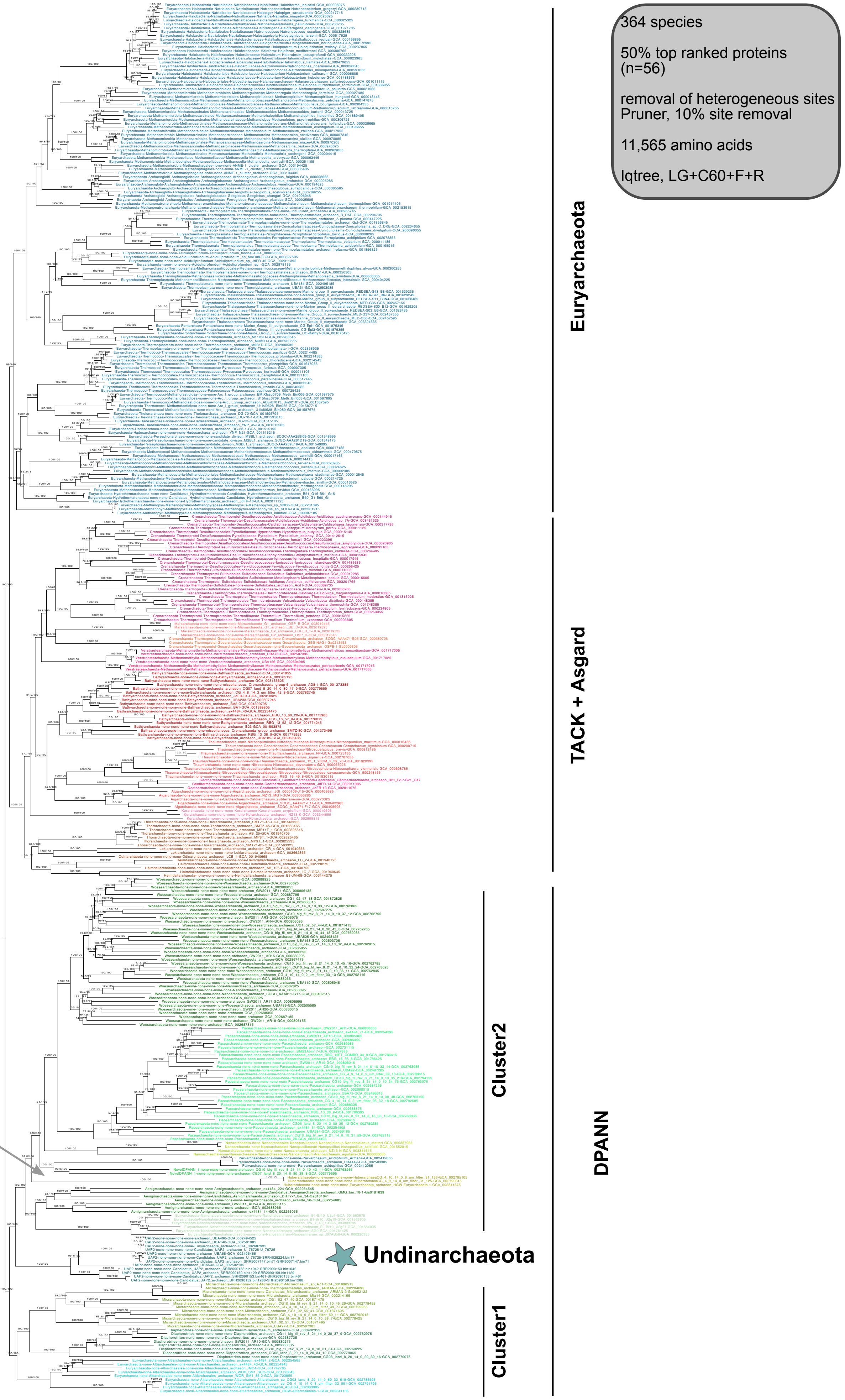


**Supplementary Figure 17 | Phylogenetic placement of Undinarchaeota based on an alignment generated with the 50% top ranked proteins (n=56) and the 364 species set.** 10% of heterogeneous sites were removed from the alignment using the chi2 test (alignment length = 11,565 aa). A ML phylogenetic tree was inferred with the LG+C60+F+R model with an ultrafast bootstrap approximation (left) and SH-like approximate likelihood tests (right), each run with 1000 replicates. The tree was artificially rooted with the DPANN archaea and the grey arrow shows the root position inferred with minimal ancestor deviation rooting (Tria et al., 2017). Scale bar: Average number of substitutions per site. Tree statistics for tree number 12 can be found in Supplementary Table 6.

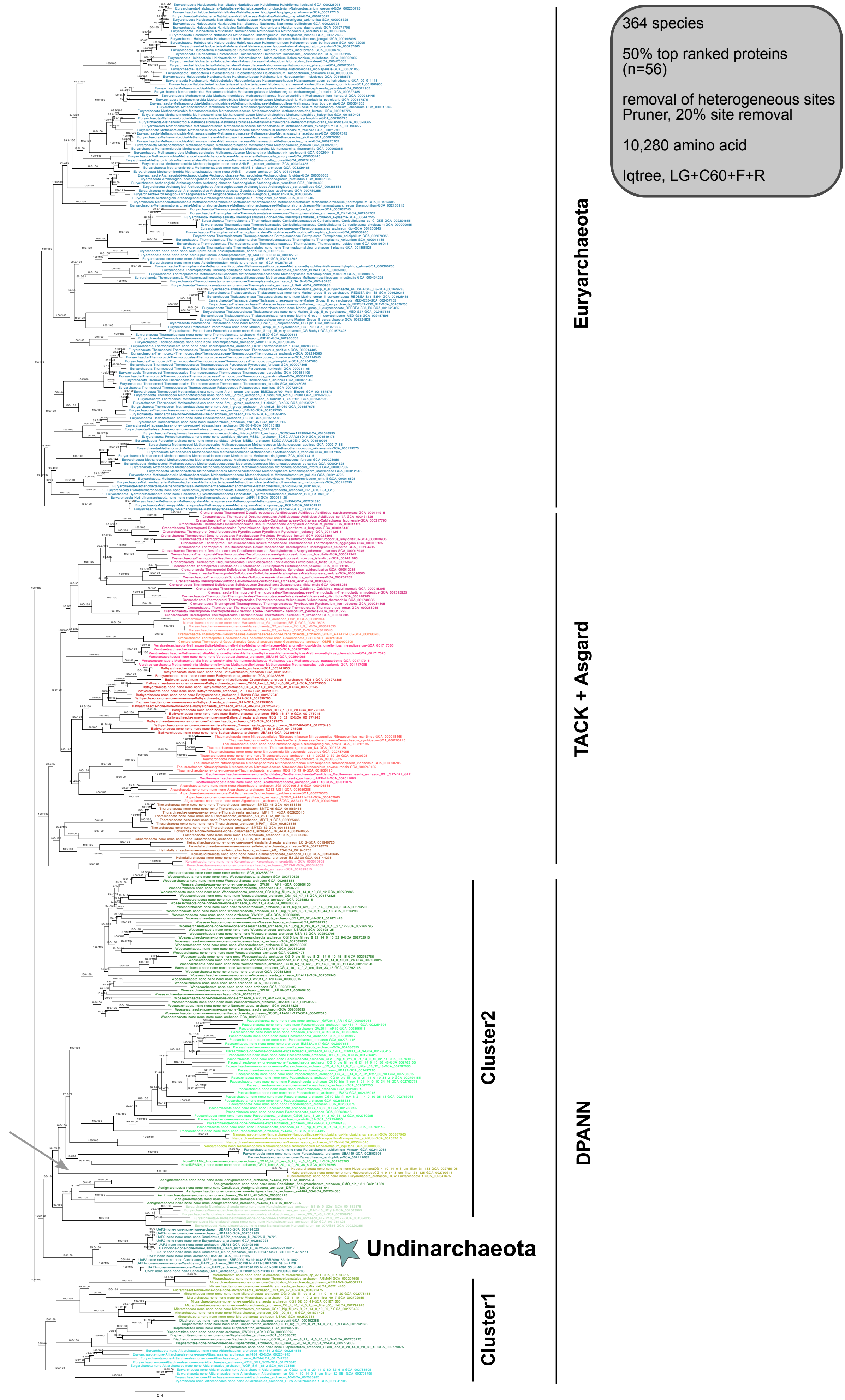


**Supplementary Figure 18 | Phylogenetic placement of Undinarchaeota based on an alignment generated with the 50% top ranked proteins (n=56) and the 364 species set.** 20% of heterogeneous sites were removed from the alignment using the chi2 test (alignment length = 10,280 aa). A ML phylogenetic tree was inferred with the LG+C60+F+R model with an ultrafast bootstrap approximation (left) and SH-like approximate likelihood tests (right), each run with 1000 replicates. The tree was artificially rooted with the DPANN archaea and the grey arrow shows the root position inferred with minimal ancestor deviation rooting (Tria et al., 2017). Scale bar: Average number of substitutions per site. Tree statistics for tree number 13 can be found in Supplementary Table 6.

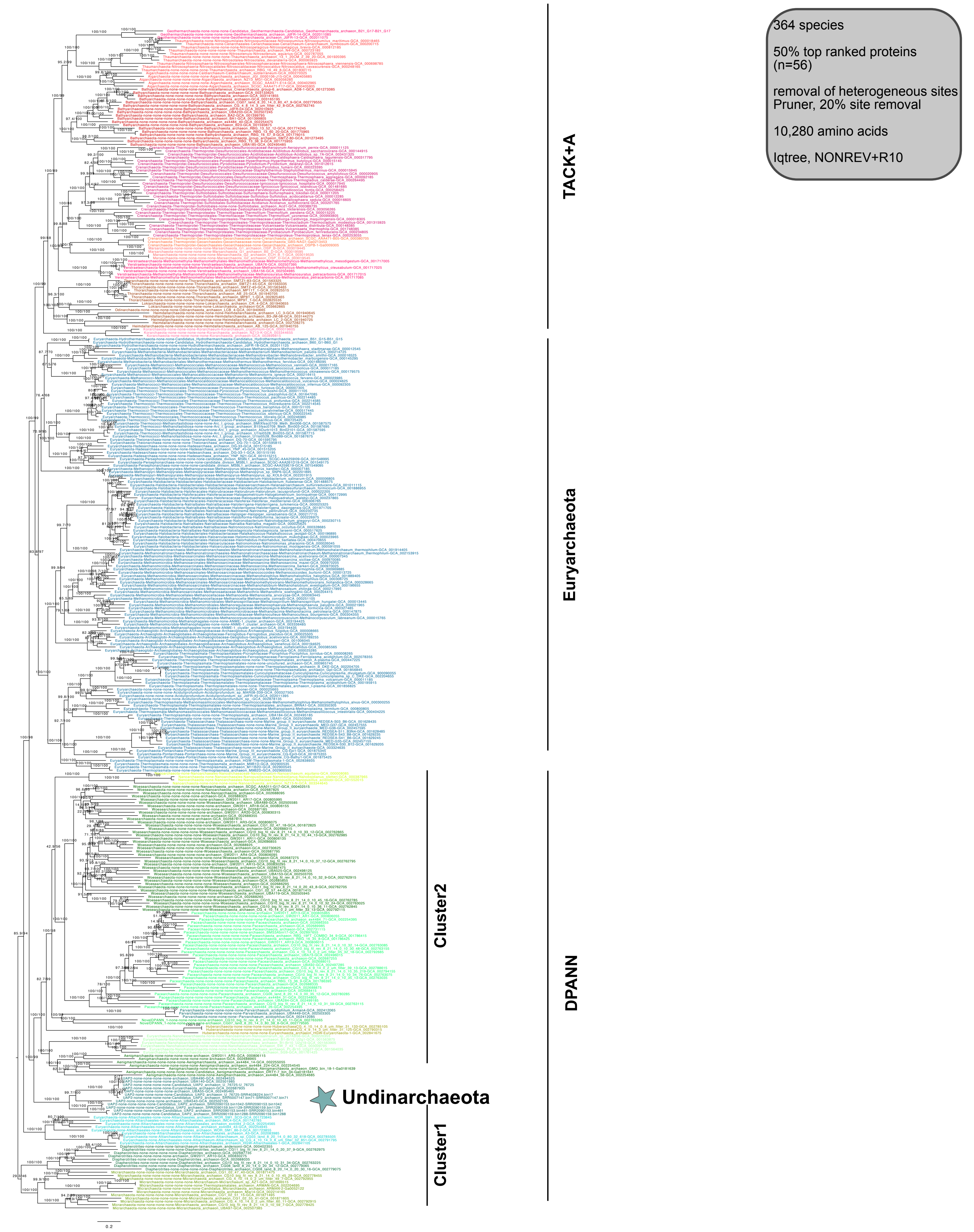


**Supplementary Figure 19 | Phylogenetic placement of Undinarchaeota based on an alignment generated with the 50% top ranked proteins (n=56) and the 364 species set.** 20% of heterogeneous sites were removed from the alignment with the chi2 test (alignment length = 10,280 aa). An ML phylogenetic tree was inferred with a non-reversible model (NONREV+R10) with an ultrafast bootstrap approximation (left) and SH-like approximate likelihood tests (right), each run with 1000 replicates. The root was inferred with the non-reversible model in iqtree. Scale bar: Average number of substitutions per site. Tree statistics for tree number 14 can be found in Supplementary Table 6.

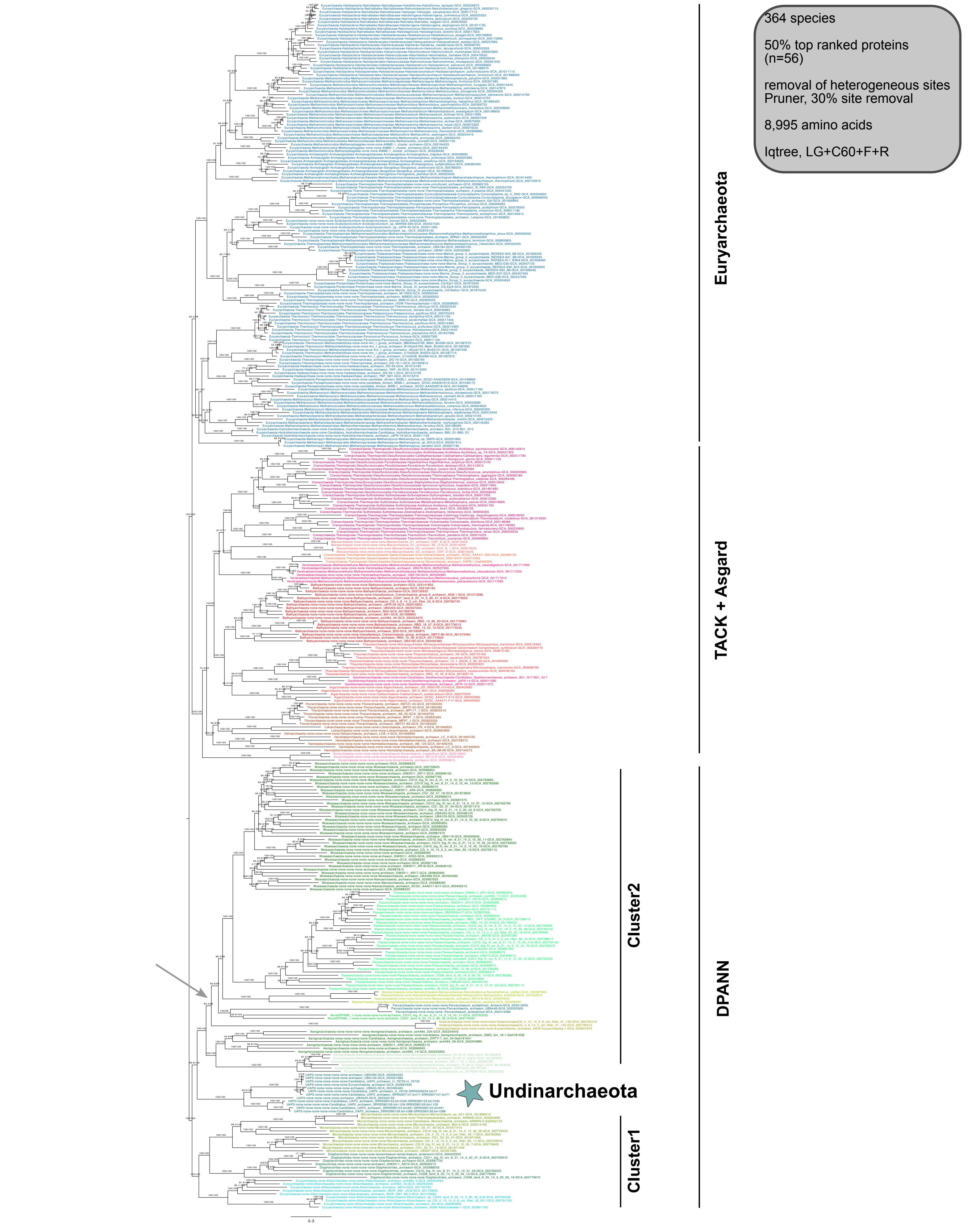


**Supplementary Figure 20 | Phylogenetic placement of Undinarchaeota based on an alignment generated with the 50% top ranked proteins (n=56) and the 364 species set.** 30% of heterogeneous sites were removed from the alignment using the chi2 test (alignment length = 8,995 aa). A ML phylogenetic tree was inferred with the LG+C60+F+R model with an ultrafast bootstrap approximation (left) and SH-like approximate likelihood tests (right), each run with 1000 replicates. The tree was artificially rooted with the DPANN archaea and the grey arrow shows the root inferred with minimal ancestor deviation rooting (Tria et al., 2017). Scale bar: Average number of substitutions per site. Tree statistics for tree number 15 can be found in Supplementary Table 6.

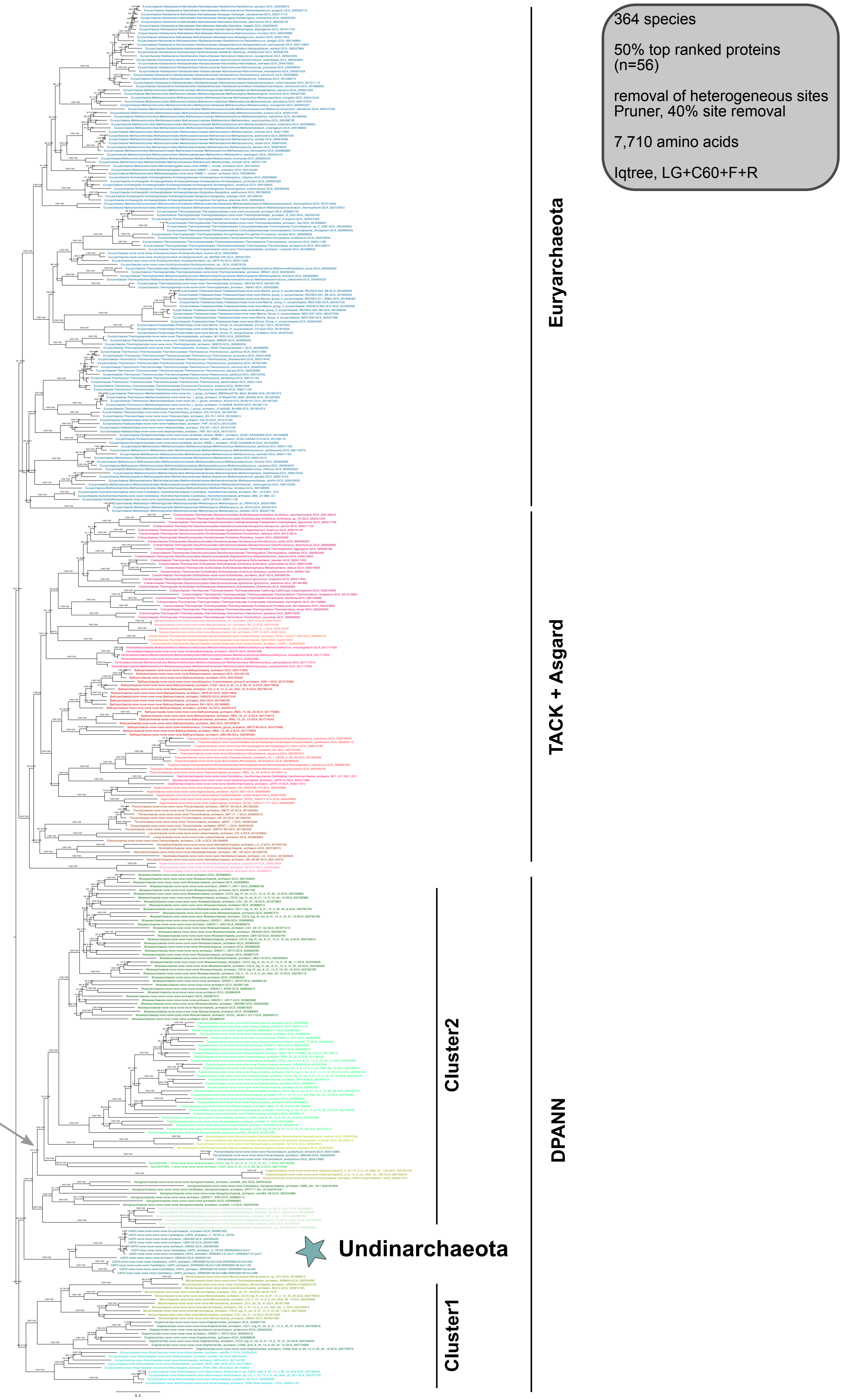


**Supplementary Figure 21 | Phylogenetic placement of Undinarchaeota based on an alignment generated with the 50% top ranked proteins (n=56) and the 364 species set.** 40% of heterogeneous sites were removed from the alignment using the chi2 test (alignment length = 7,710 aa). A ML phylogenetic tree was inferred with the LG+C60+F+R model with an ultrafast bootstrap approximation (left) and SH-like approximate likelihood tests (right), each run with 1000 replicates. The tree was artificially rooted with the DPANN archaea and the grey arrow shows the root position inferred with minimal ancestor deviation rooting (Tria et al., 2017). Scale bar: Average number of substitutions per site. Tree statistics for tree number 16 can be found in Supplementary Table 6.

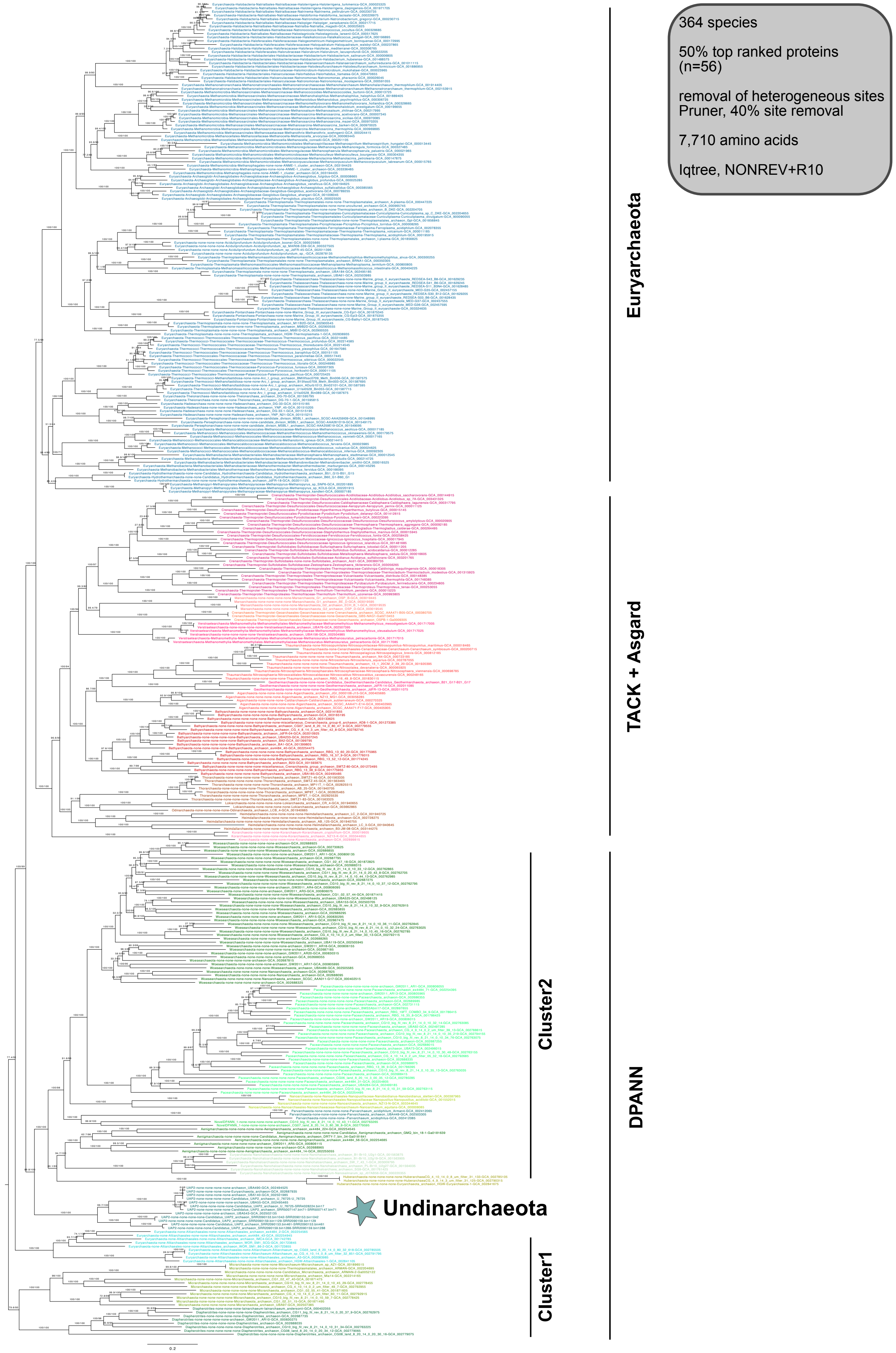


**Supplementary Figure 22 | Phylogenetic placement Undinarchaeota based on an alignment generated with the 50% top ranked proteins (n=56) and the 364 species set.** 40% of heterogeneous sites were removed from the alignment using the chi2 test (alignment length = 7,710 aa). A ML phylogenetic tree was inferred with a non-reversible model (NONREV+R10) with an ultrafast bootstrap approximation (left) and SH-like approximate likelihood tests (right), each run with 1000 replicates. The tree was rooted using the non-reversible model in iqtree. Scale bar: Average number of substitutions per site. Tree statistics for tree number 17 can be found in Supplementary Table 6.

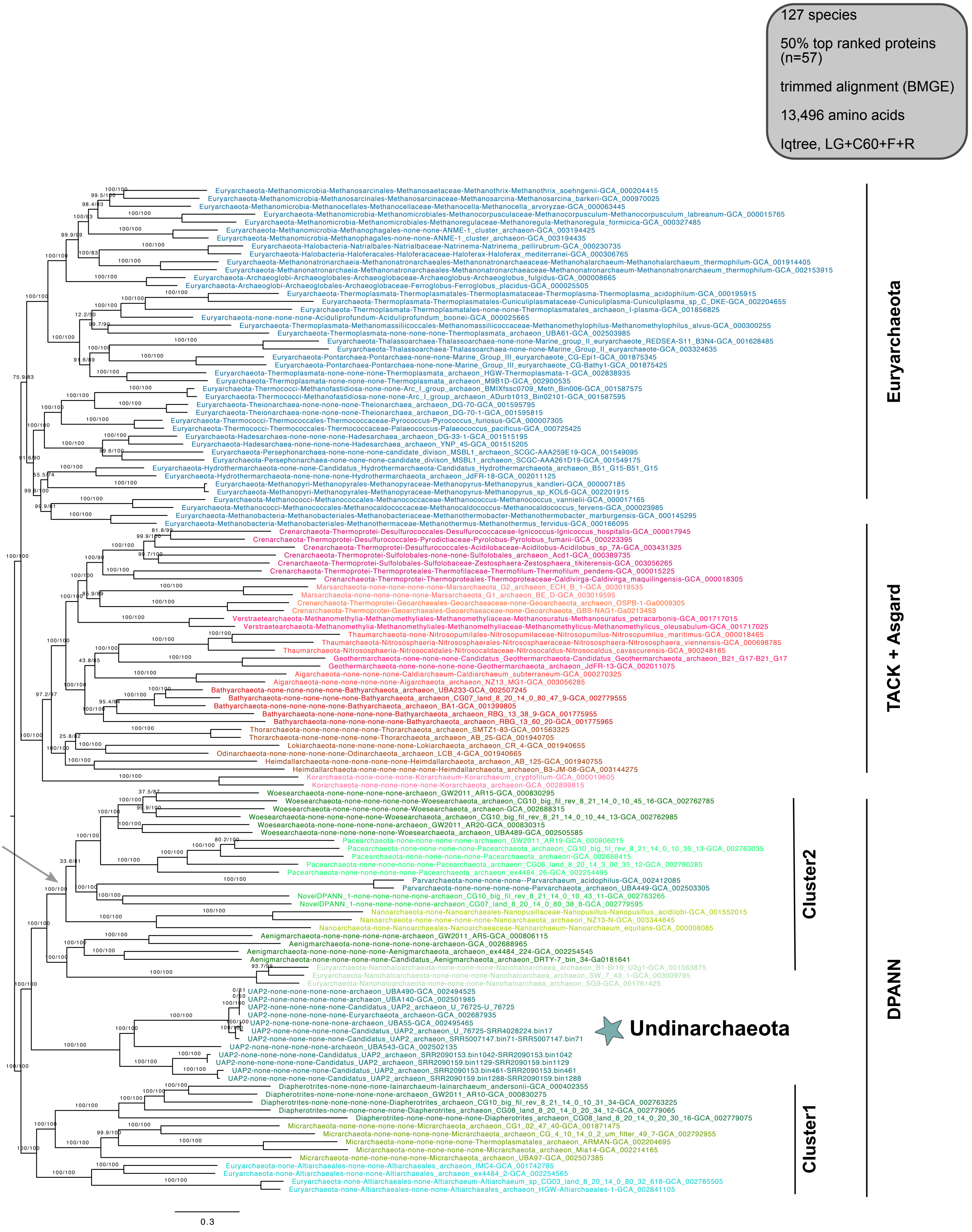


**Supplementary Figure 23 | Phylogenetic placement of Undinarchaeota based on an alignment generated with the 50% top ranked proteins (n=57) and the 127 species set.** The alignment was trimmed with BMGE (alignment length = 13,496 aa). A ML phylogenetic tree was inferred with the LG+C60+F+R model with an ultrafast bootstrap approximation (left) and SH-like approximate likelihood tests (right), each run with 1000 replicates. The tree was artificially rooted with the DPANN archaea and the grey arrow shows the root position inferred with minimal ancestor deviation rooting (Tria et al., 2017). Scale bar: Average number of substitutions per site. Tree statistics for tree number 18 can be found in Supplementary Table 6.

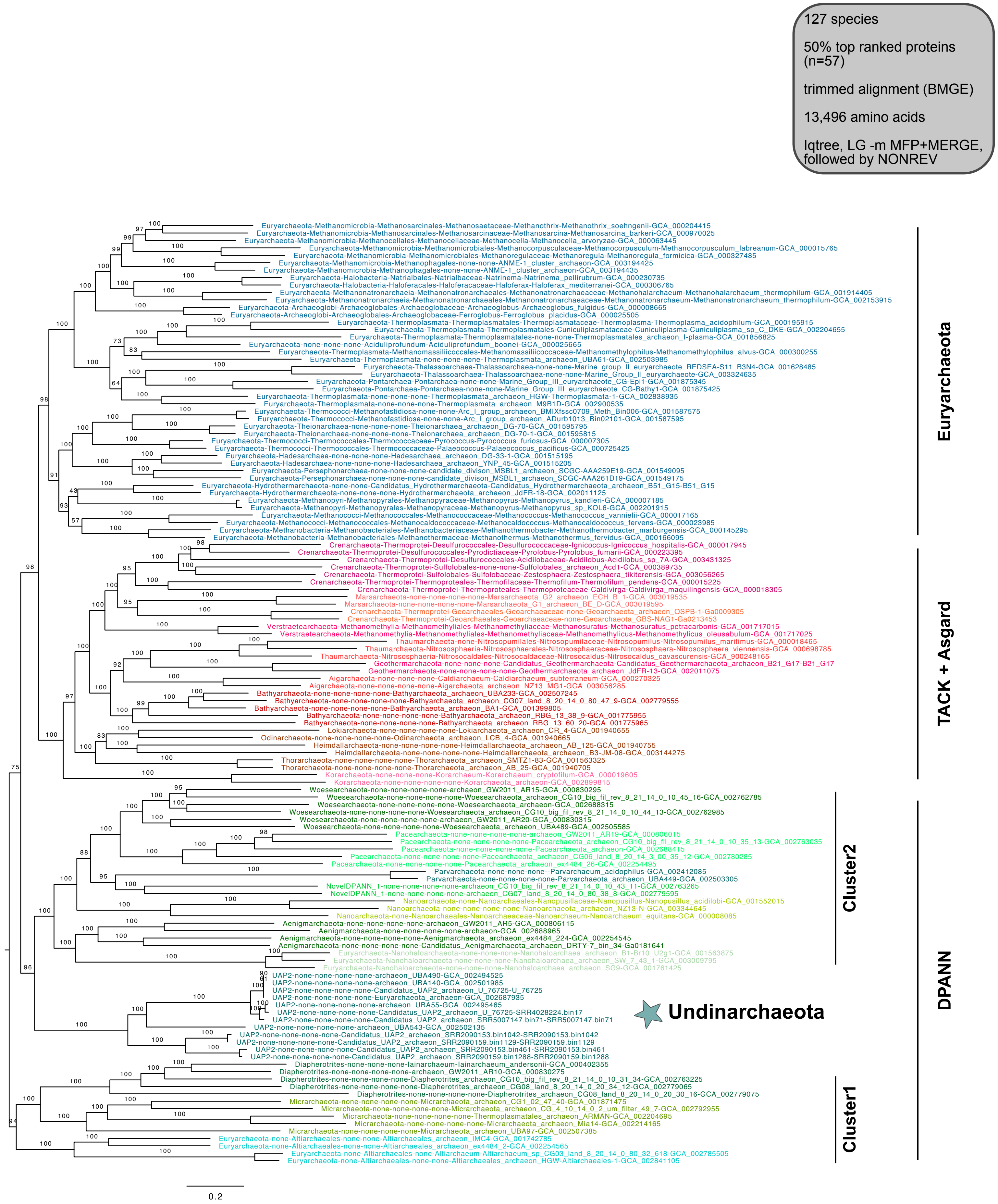


**Supplementary Figure 24 | Phylogenetic placement of Undinarchaeota based on an alignment generated with the 50% top ranked proteins (n=57) and the 127 species set.** The aligment was trimmed with BMGE (alignment length= 13,496 aa). An initial ML phylogenetic tree was inferred with the LG model (-m MFP+MERGE) followed by a tree generated with a non-reversible model with an ultrafast bootstrap approximation run with 1000 replicates. The tree was rooted with the non-reversible model in iqtree. Scale bar: Average number of substitutions per site. Tree statistics for tree number 19 can be found in Supplementary Table 6.

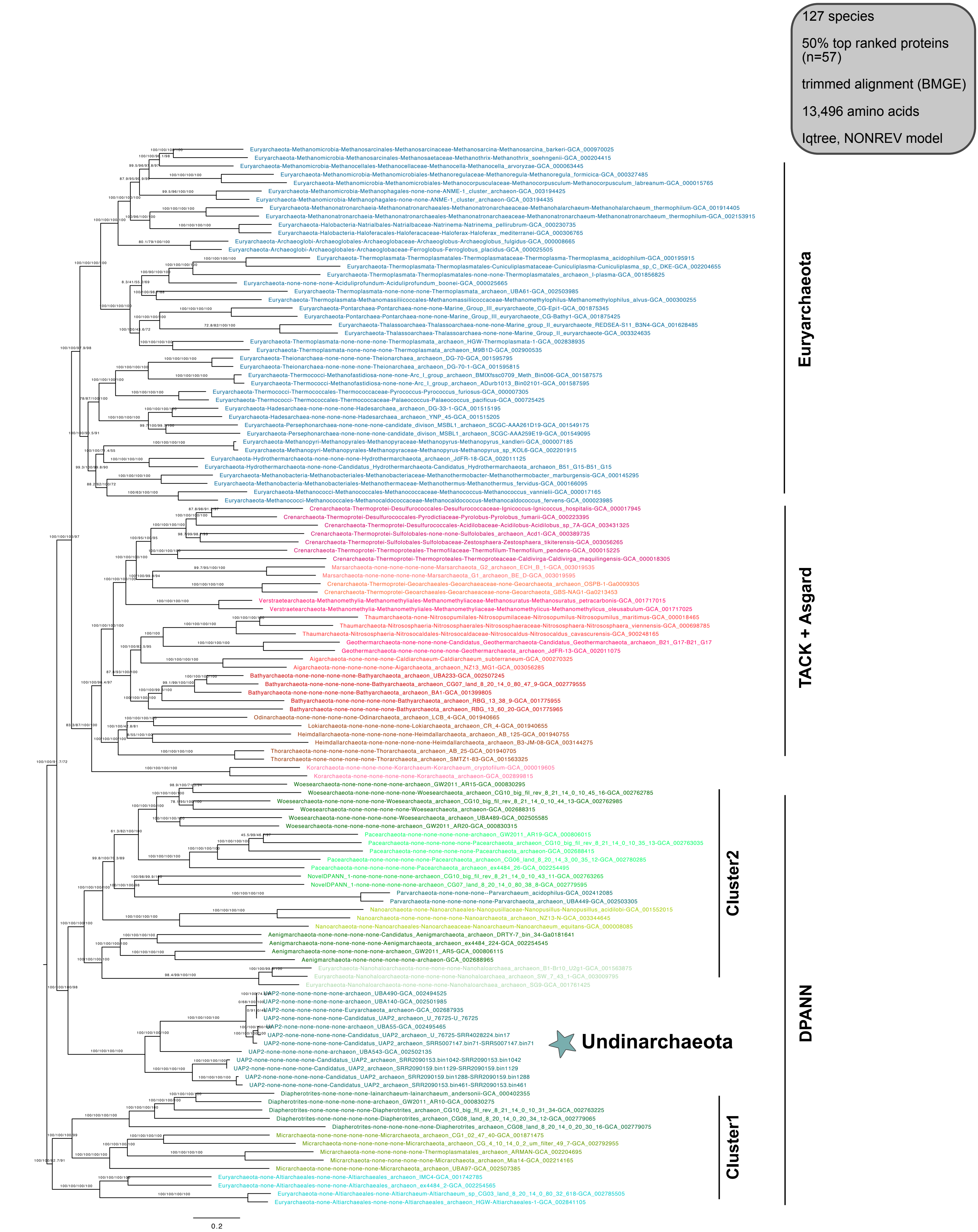


**Supplementary Figure 25 | Phylogenetic placement of Undinarchaeota based on an alignment generated with the 50% top ranked proteins (n=57) and the 127 species set.** The alignment was trimmed with BMGE (alignment length = 13,496 aa). A ML phylogenetic tree was inferred with the NONREV model. The first two values show the support for the reversible and the second two for the non-reversible mode. Values 1 and 3 were generated with an ultrafast bootstrap approximation and values 2 and 4 with the SH-like approximate likelihood tests, each run with 1000 replicates. The tree was rooted with the non-reversible model in iqtree. Scale bar: Average number of substitutions per site. Tree statistics for tree number 20 can be found in Supplementary Table 6.

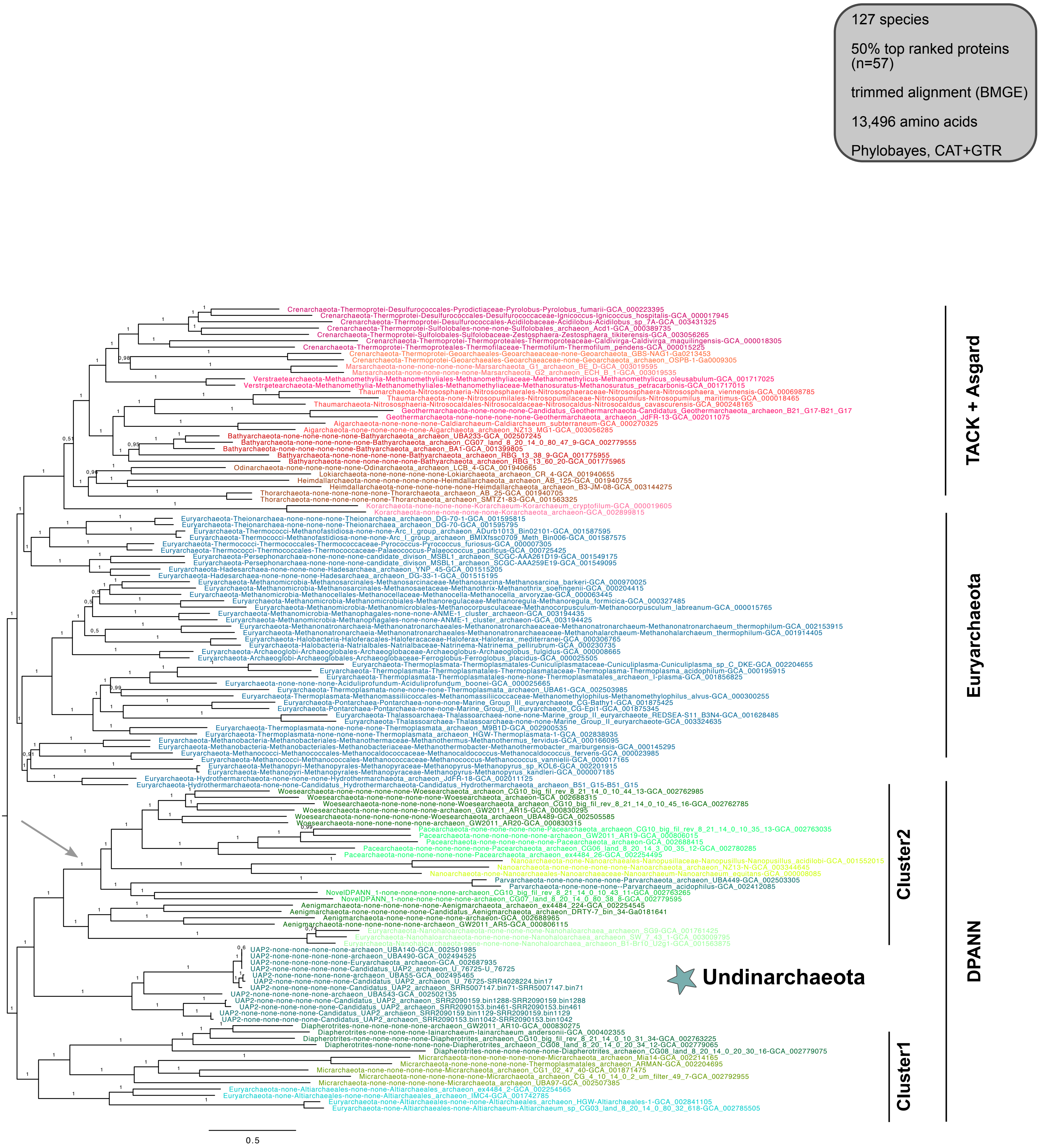


**Supplementary Figure 26 | Phylogenetic placement of Undinarchaeota based on an alignment generated with the 50% top ranked proteins (n=57) and the 127 species set.** The alignment was trimmed with BMGE (alignment length = 13,496 aa). A Bayesian phylogenetic tree was inferred with the CAT+GTR model with 14,107 cycles (25% burn-in). The tree was artificially rooted with the DPANN archaea and the grey arrow shows the root position as inferred with minimal ancestor deviation rooting (Tria et al., 2017). Scale bar: Average number of substitutions per site. Tree statistics for tree number 21 can be found in Supplementary Table 6.

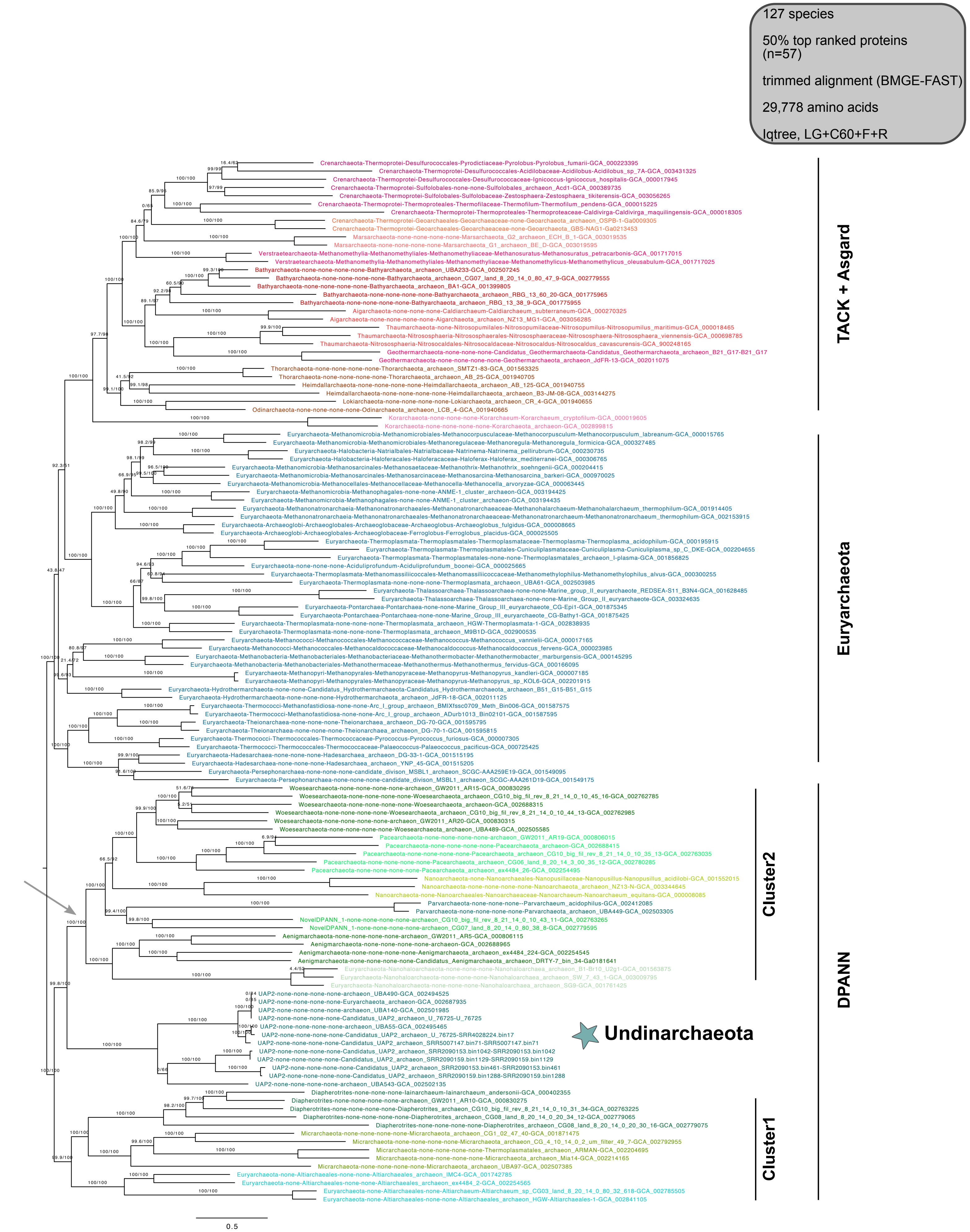


**Supplementary Figure 27 | Phylogenetic placement of Undinarchaeota based on an alignment generated with the 50% top ranked proteins (n=57) and the 127 species set.** The alignment was trimmed with BMGE-FAST (alignment length= 29,778 aa). A ML phylogenetic tree was inferred with the LG+C60+F+R model with an ultrafast bootstrap approximation (left) and SH-like approximate likelihood tests (right), each run with 1000 replicates. The tree was artificially rooted with the DPANN archaea and the grey arrow shows the root position inferred with minimal ancestor deviation rooting (Tria et al., 2017). Scale bar: Average number of substitutions per site. Tree statistics for tree number 22 can be found in Supplementary Table 6.

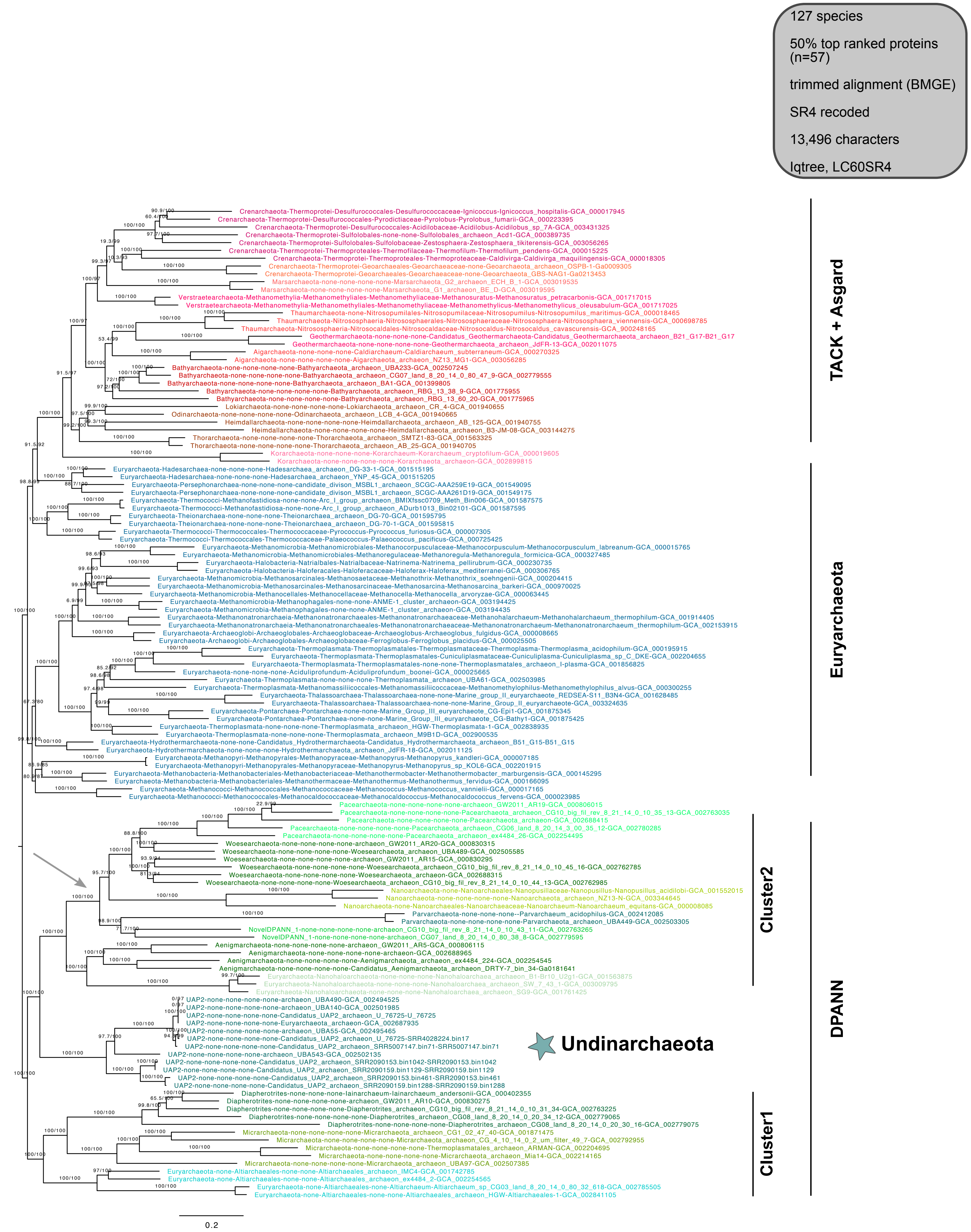


**Supplementary Figure 28 | Phylogenetic placement of Undinarchaeota based on an alignment generated with the 50% top ranked proteins (n=57) and the 127 species set.** The alignment was trimmed with BMGE and decoded into 4 character states (SR4 recoding; alignment length = 13,496 characters). A ML phylogenetic tree was inferred with the C60SR4 model with an ultrafast bootstrap approximation (left) and SH-like approximate likelihood tests (right), each run with 1000 replicates. The tree was artificially rooted with the DPANN archaea and the grey arrow shows the root position inferred with minimal ancestor deviation rooting (Tria et al., 2017). Scale bar: Average number of substitutions per site. Tree statistics for tree number 23 can be found in Supplementary Table 6.

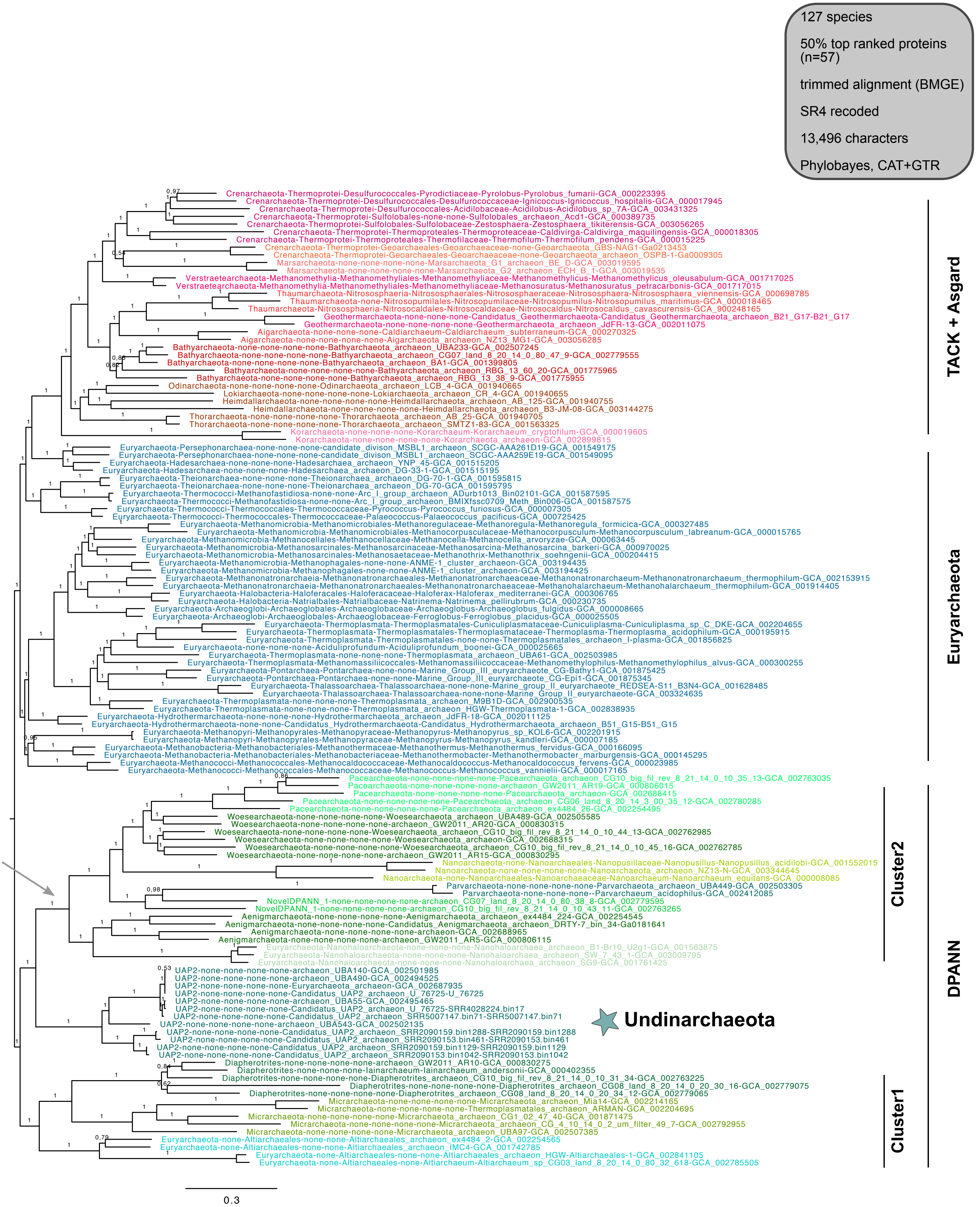


**Supplementary Figure 29 | Phylogenetic placement of Undinarchaeota based on an alignment generated with the 50% top ranked proteins (n=57) and the 127 species set.** The alignment was trimmed with BMGE and recoded into 4 character states (SR4 decoding; alignment length = 13,496 characters). A Bayesian phylogenetic tree was inferred with the CAT+GTR model. The tree was artificially rooted with the DPANN archaea and the grey arrow shows the root inferred with minimal ancestor deviation rooting (Tria et al., 2017). Scale bar: Average number of substitutions per site. Tree statistics for tree number 24 can be found in Supplementary Table 6.

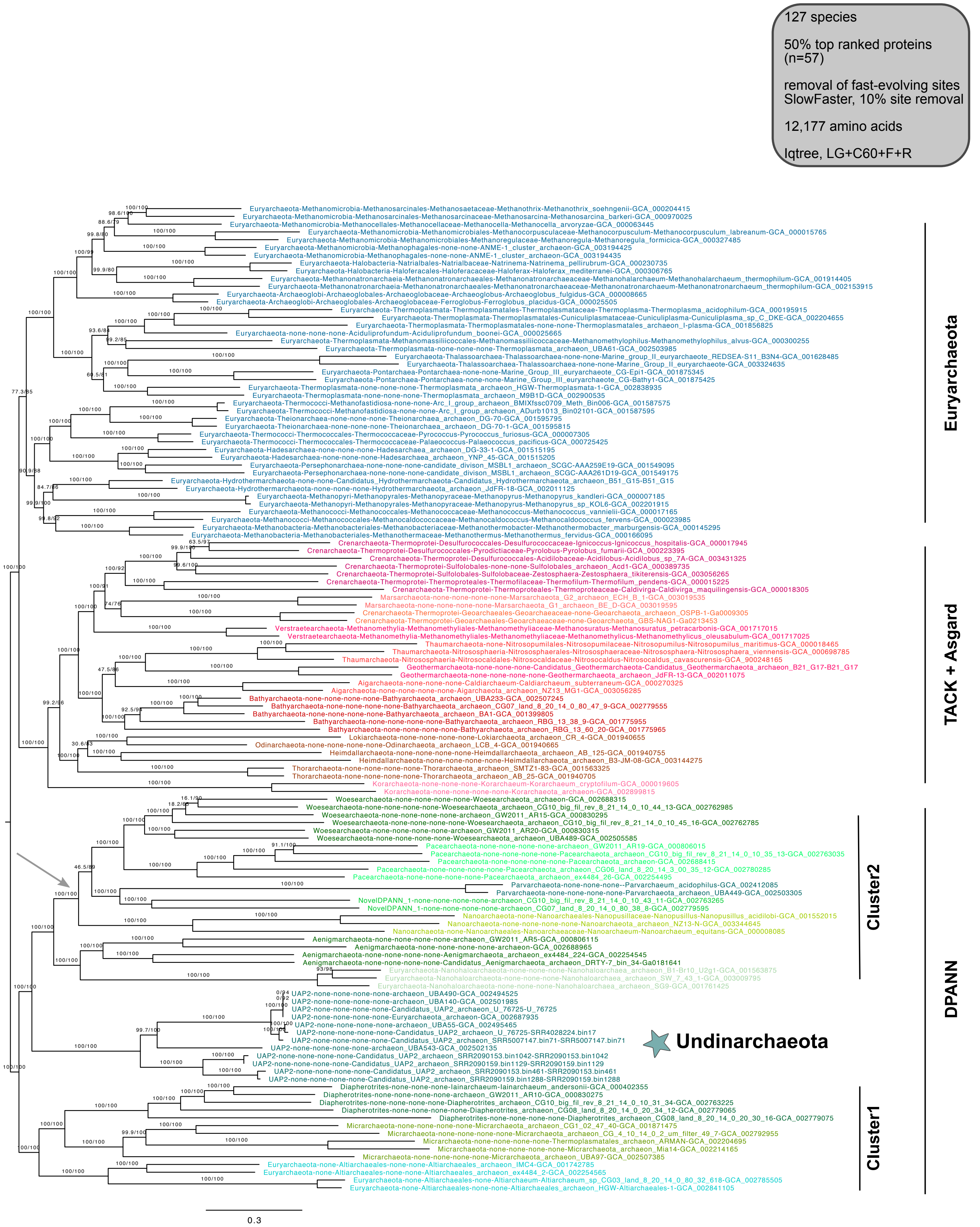


**Supplementary Figure 30 | Phylogenetic placement of Undinarchaeota based on an alignment generated with the 50% top ranked proteins (n=57) and the 127 species set.** 10% of the fast-evolving sites were removed from the alignment with SlowFaster (alignment length = 12,177 aa). A ML phylogenetic tree was inferred with the LG+C60+F+R model with an ultrafast bootstrap approximation (left) and SH-like approximate likelihood tests (right), each run with 1000 replicates. The tree was artificially rooted with the DPANN archaea and the grey arrow shows the root inferred with minimal ancestor deviation rooting (Tria et al., 2017). Scale bar: Average number of substitutions per site. Tree statistics for tree number 25 can be found in Supplementary Table 6.

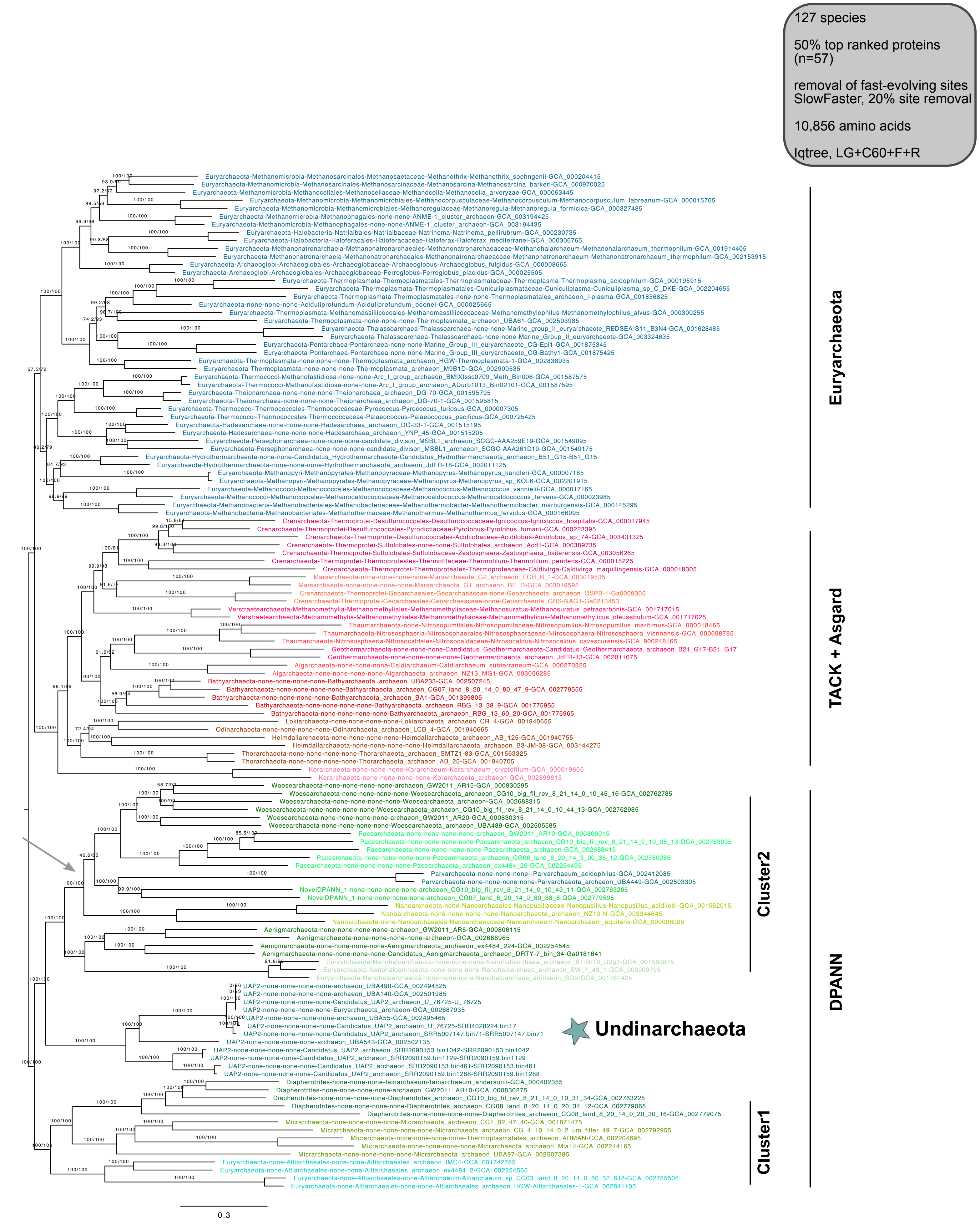


**Supplementary Figure 31 | Phylogenetic placement of Undinarchaeota based on an alignment generated with the 50% top ranked proteins (n=57) and the 127 species set.** 20% of the fast-evolving sites were removed from the alignment with SlowFaster (alignment length = 10,856 aa). A ML phylogenetic tree was inferred with the LG+C60+F+R model with an ultrafast bootstrap approximation (left) and SH-like approximate likelihood tests (right), each run with 1000 replicates. The tree was artificially rooted with the DPANN archaea and the grey arrow shows the root position inferred with minimal ancestor deviation rooting (Tria et al., 2017). Scale bar: Average number of substitutions per site.Tree statistics for tree number 26 can be found in Supplementary Table 6.

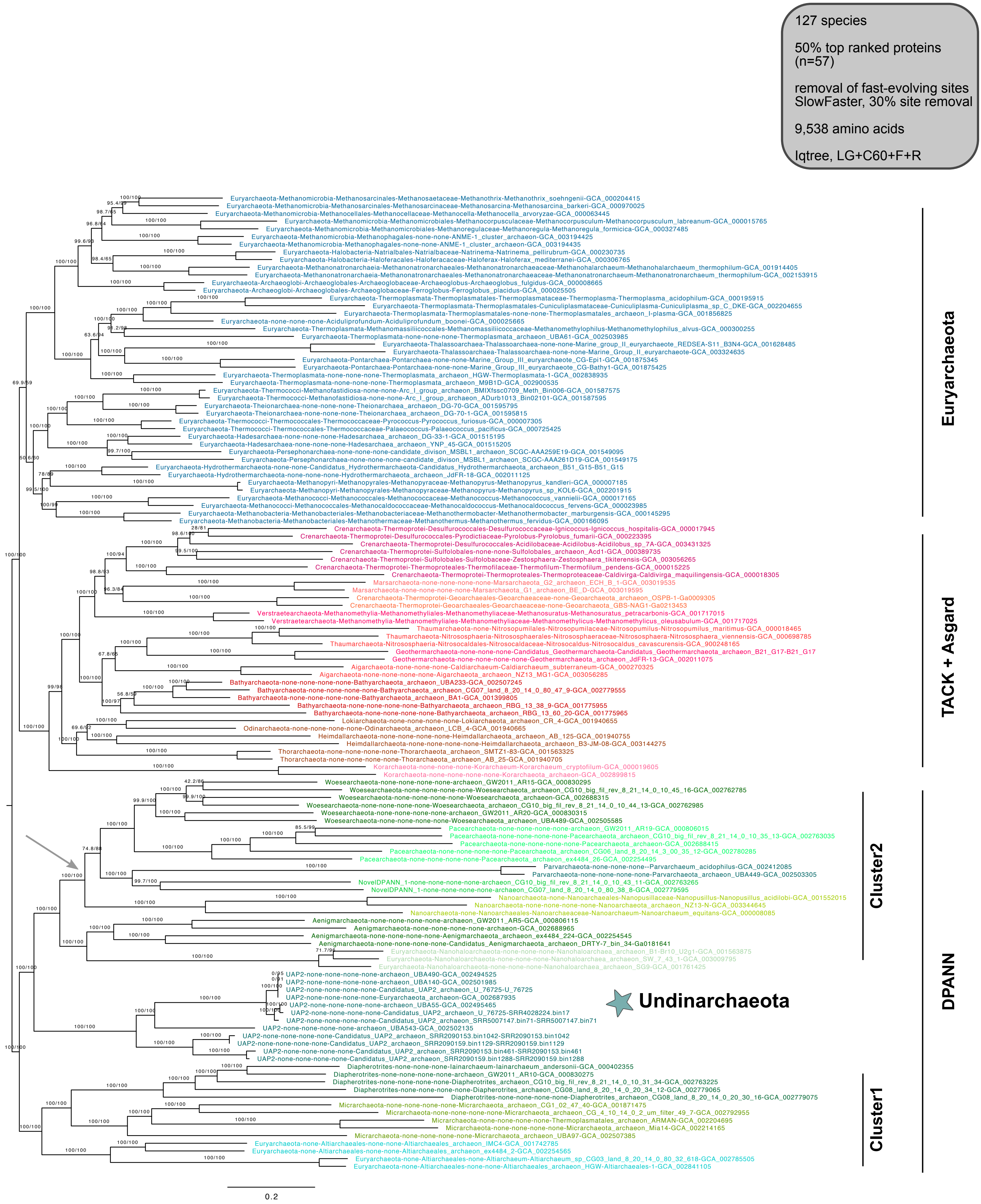


**Supplementary Figure 32 | Phylogenetic placement of Undinarchaota based on an alignment generated with the 50% top ranked proteins (n=57) and the 127 species set.** 30% of the fast-evolving sites were removed from the alignment with SlowFaster (alignment length = 9,538 aa). A ML phylogenetic tree was inferred with the LG+C60+F+R model with an ultrafast bootstrap approximation (left) and SH-like approximate likelihood tests (right), each run with 1000 replicates. The tree was artificially rooted with the DPANN archaea and the grey arrow shows the root position inferred with minimal ancestor deviation rooting (Tria et al., 2017). Scale bar: Average number of substitutions per site. Tree statistics for tree number 27 can be found in Supplementary Table 6.

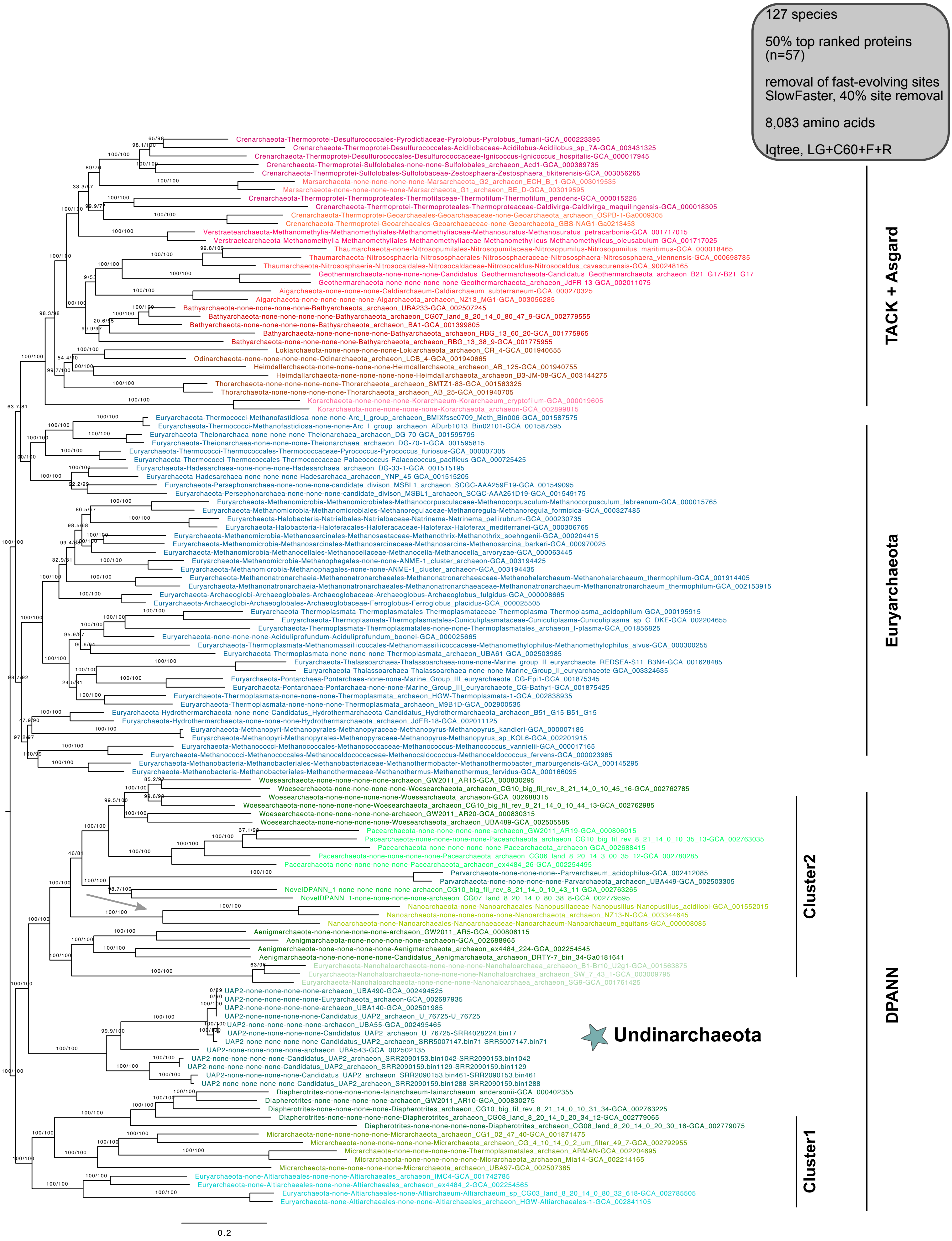


**Supplementary Figure 33 | Phylogenetic placement of Undinarchaota based on an alignment generated with the 50% top ranked proteins (n=57) and the 127 species set.** 40% of the fast-evolving sites were removed from the alignment with SlowFaster (alignment length = 8,083 aa). A ML phylogenetic tree was inferred with the LG+C60+F+R model with an ultrafast bootstrap approximation (left) and SH-like approximate likelihood tests (right), each run with 1000 replicates. The tree was artificially rooted with the DPANN archaea and the grey arrow points to the root position as inferred by minimal ancestor deviation rooting (Tria et al., 2017). Scale bar: Average number of substitutions per site. Tree statistics for tree number 28 can be found in Supplementary Table 6.

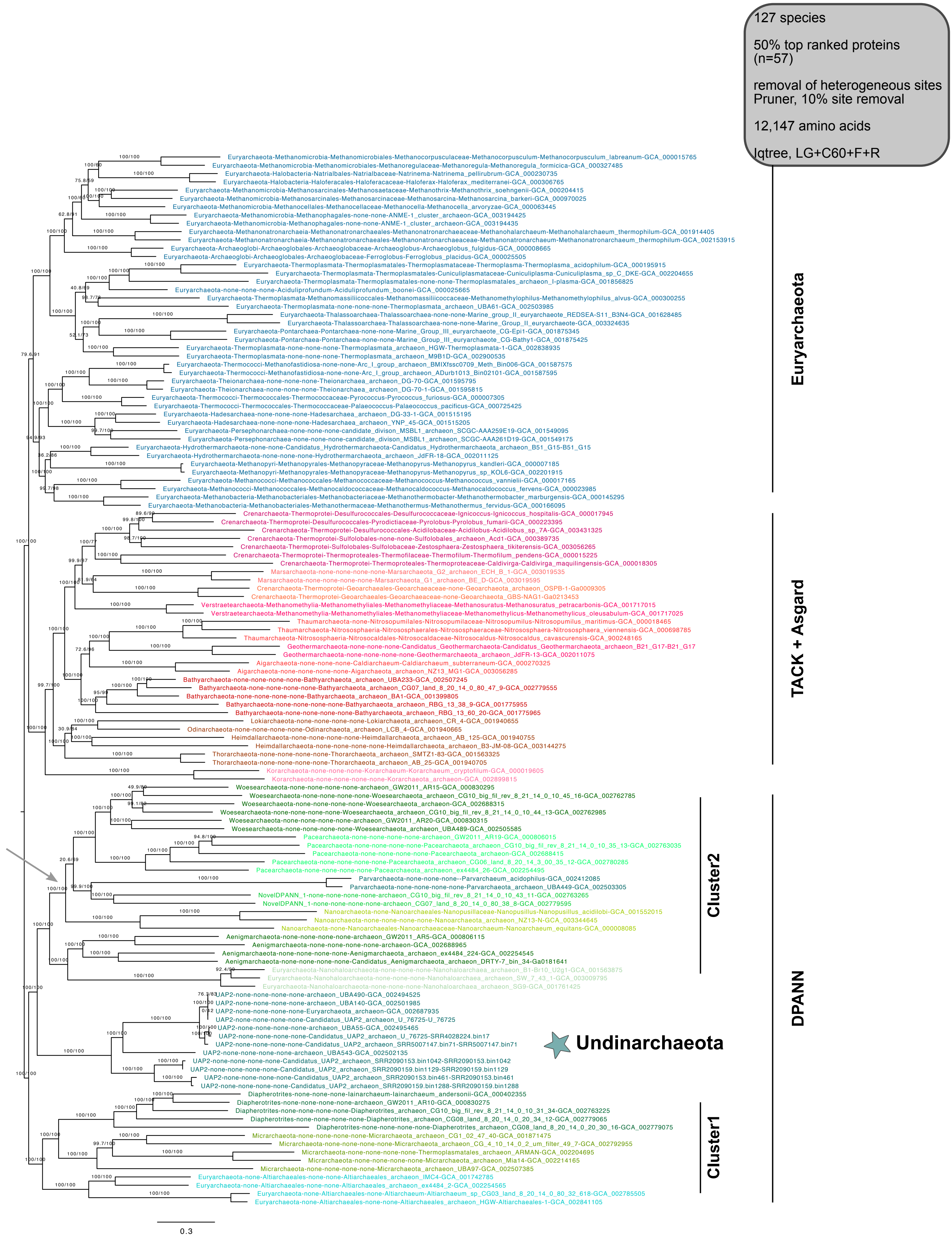


**Supplementary Figure 34 | Phylogenetic placement Undinarchaeota based on an alignment generated with the 50% top ranked proteins (n=57) and the 127 species set.** 10% of the heterogeneous sites were removed from the alignment using the chi2 test (alignment length = 12,147 aa). A ML phylogenetic tree was inferred with the LG+C60+F+R model with an ultrafast bootstrap approximation (left) and SH-like approximate likelihood tests (right), each run with 1000 replicates. The tree was artificially rooted with the DPANN archaea and the grey arrow shows the root position inferred with minimal ancestor deviation rooting (Tria et al., 2017). Scale bar: Average number of substitutions per site. Tree statistics for tree number 29 can be found in Supplementary Table 6.

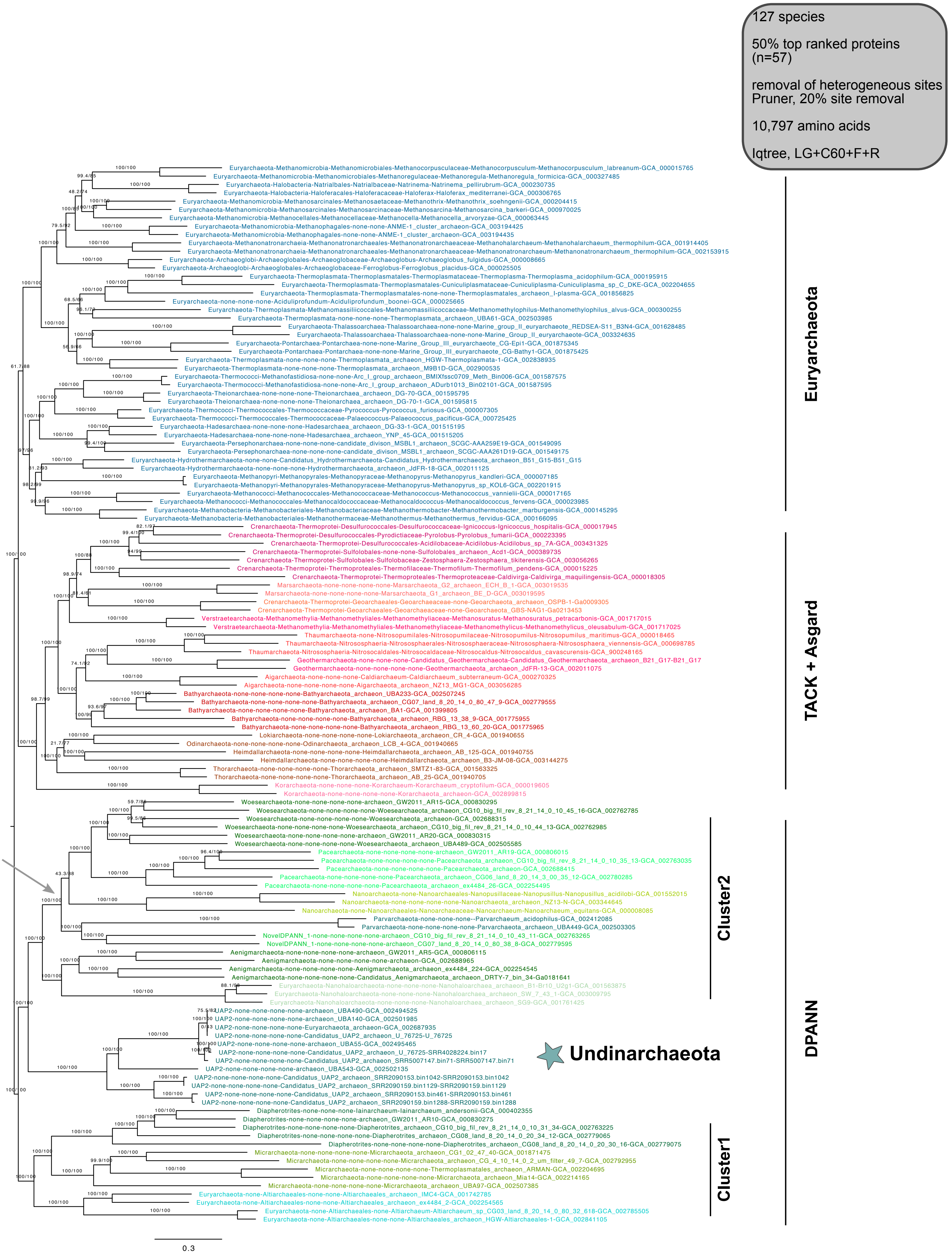


**Supplementary Figure 35 | Phylogenetic placement of Undinarchaeota based on an alignment generated with the 50% top ranked proteins (n=57) and the 127 species set.** 20% of the heterogeneous sites were removed from the alignment with the chi2 test (alignment length = 10,797 aa). A ML phylogenetic tree was inferred with the LG+C60+F+R model with an ultrafast bootstrap approximation (left) and SH-like approximate likelihood tests (right), each run with 1000 replicates. The tree was artificially rooted with the DPANN archaea and the grey arrow shows the root position inferred with minimal ancestor deviation rooting (Tria et al., 2017). Scale bar: Average number of substitutions per site. Tree statistics for tree number 30 can be found in Supplementary Table 6.

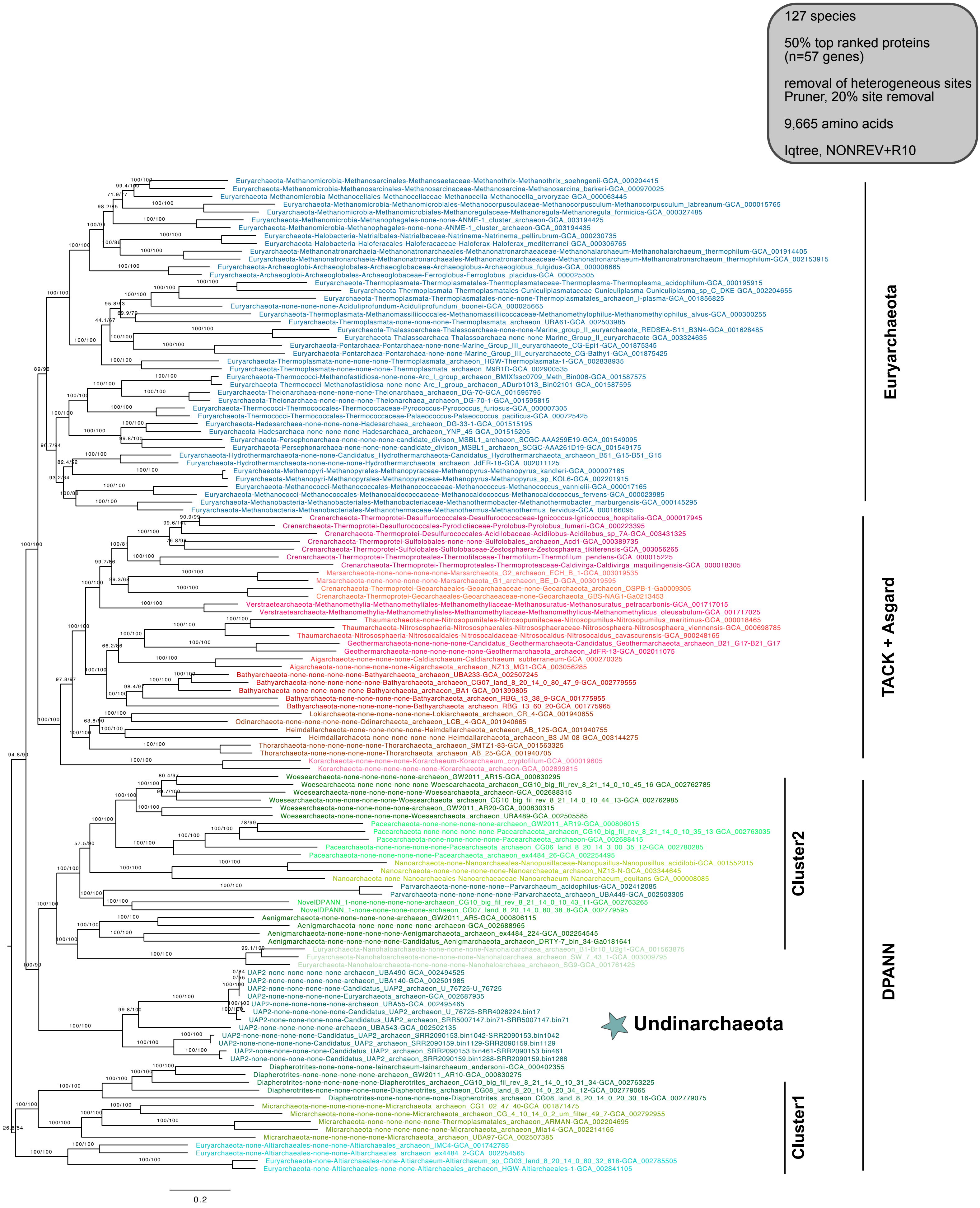


**Supplementary Figure 36 | Phylogenetic placement of Undinarchaeota based on an alignment generated with the 50% top ranked proteins (n=57) and the 127 species set.** 20% of the heterogeneous sites were removed from the alignment with the chi2 test (alignment length = 9,665 aa). A ML phylogenetic tree was inferred with the NONREV model with an ultrafast bootstrap approximation (left) and SH-like approximate likelihood tests (right), each run with 1000 replicates. The tree was rooted using the non-reversible model in iqtree. Scale bar: Average number of substitutions per site. Tree statistics for tree number 31 can be found in Supplementary Table 6.

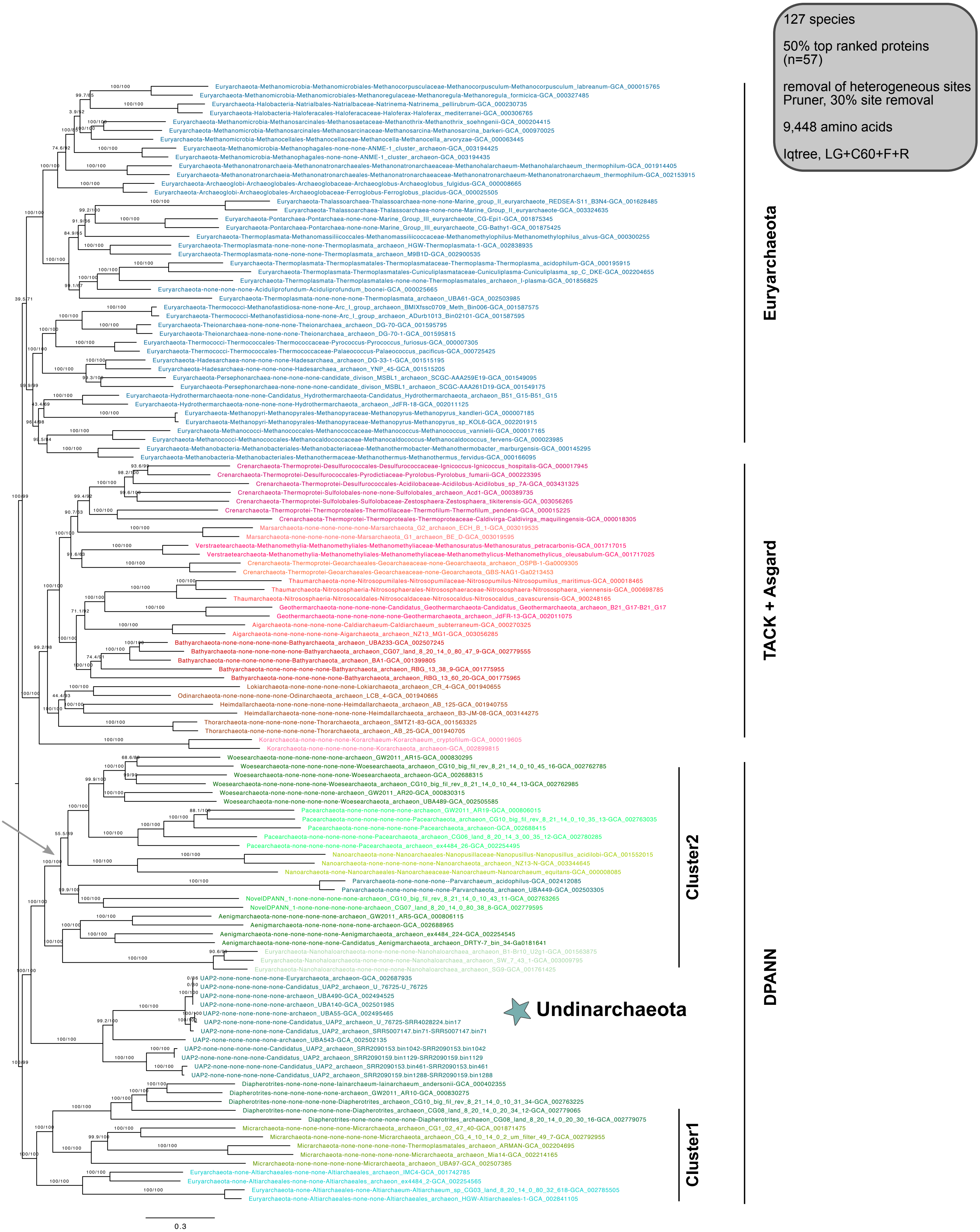


**Supplementary Figure 37 | Phylogenetic placement of Undinarchaeota based on an alignment generated with the 50% top ranked proteins (n=57) and the 127 species set.** 30% of the heterogeneous sites were removed from the alignment using the chi2 test (alignment length = 9,448 aa). A ML phylogenetic tree was inferred with the LG+C60+F+R model with an ultrafast bootstrap approximation (left) and SH-like approximate likelihood tests (right), each run with 1000 replicates. The tree was artificially rooted with the DPANN archaea and the grey arrow shows the root position inferred with minimal ancestor deviation rooting (Tria et al., 2017). Scale bar: Average number of substitutions per site. Tree statistics for tree number 32 can be found in Supplementary Table 6.

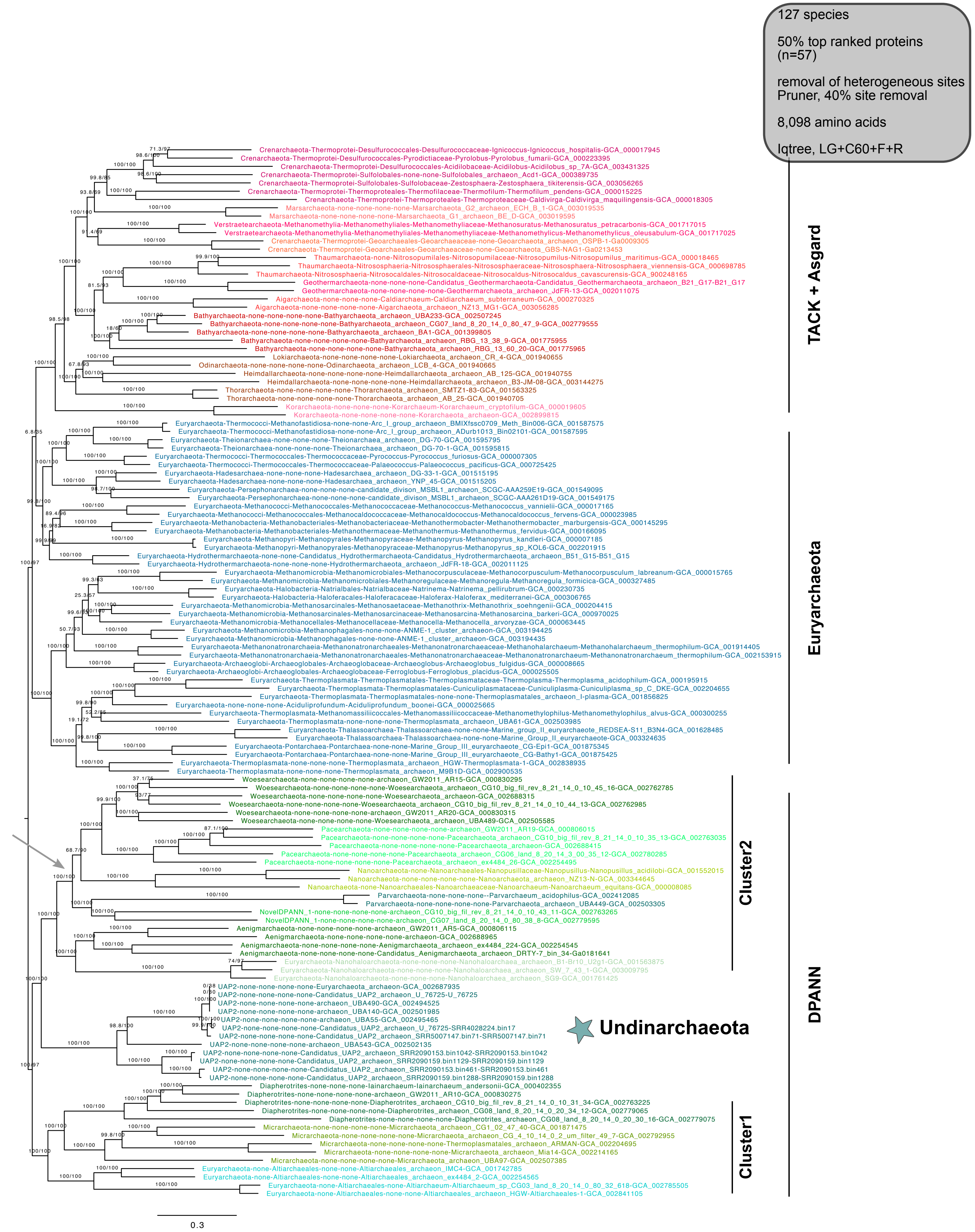


**Supplementary Figure 38 | Phylogenetic placement of Undinarchaeota based on an alignment generated with the 50% top ranked proteins (n=57) and the 127 species set.** 40% of the heterogeneous sites were removed from the alignment using the chi2 test (alignment length = 8,098 aa). A ML phylogenetic tree was inferred with the LG+C60+F+R model with an ultrafast bootstrap approximation (left) and SH-like approximate likelihood tests (right), each run with 1000 replicates. The tree was artificially rooted with the DPANN archaea and the grey arrow shows the root position inferred with minimal ancestor deviation rooting. Scale bar: Average number of substitutions per site. Tree statistics for tree number 33 can be found in Supplementary Table 6.

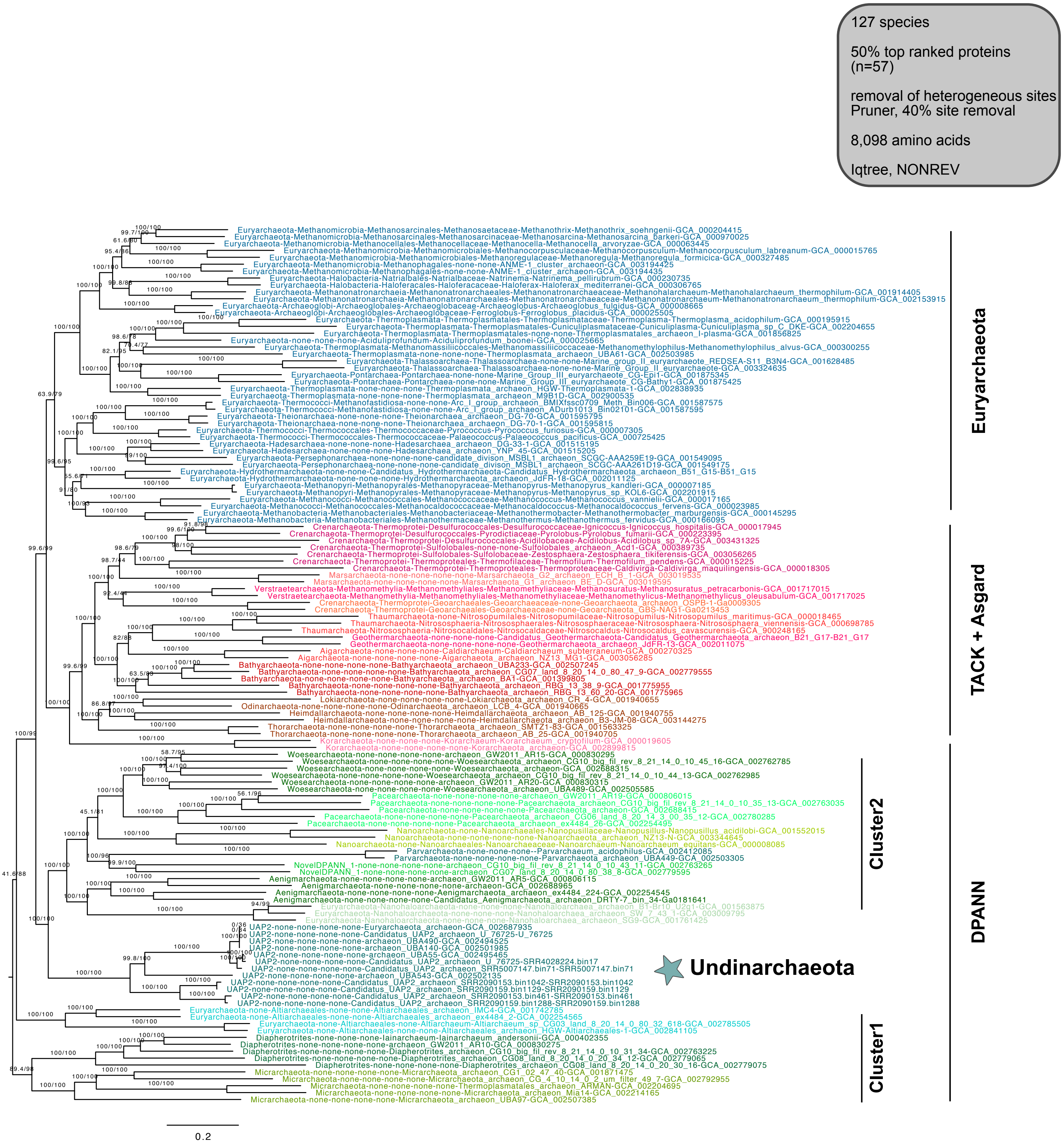


**Supplementary Figure 39 | Phylogenetic placement of Undinarchaeota based on an alignment generated with the 50% top ranked proteins (n=57) and the 127 species set.** 40% of the heterogeneous sites were removed from the alignment using the chi2 test (alignment length = 8,098 aa). A ML phylogenetic tree was inferred with the NONREV model with an ultrafast bootstrap approximation (left) and SH-like approximate likelihood tests (right), each run with 1000 replicates. The tree was rooted with the non-reversible model in iqtree. Scale bar: Average number of substitutions per site. Tree statistics for tree number 34 can be found in Supplementary Table 6.

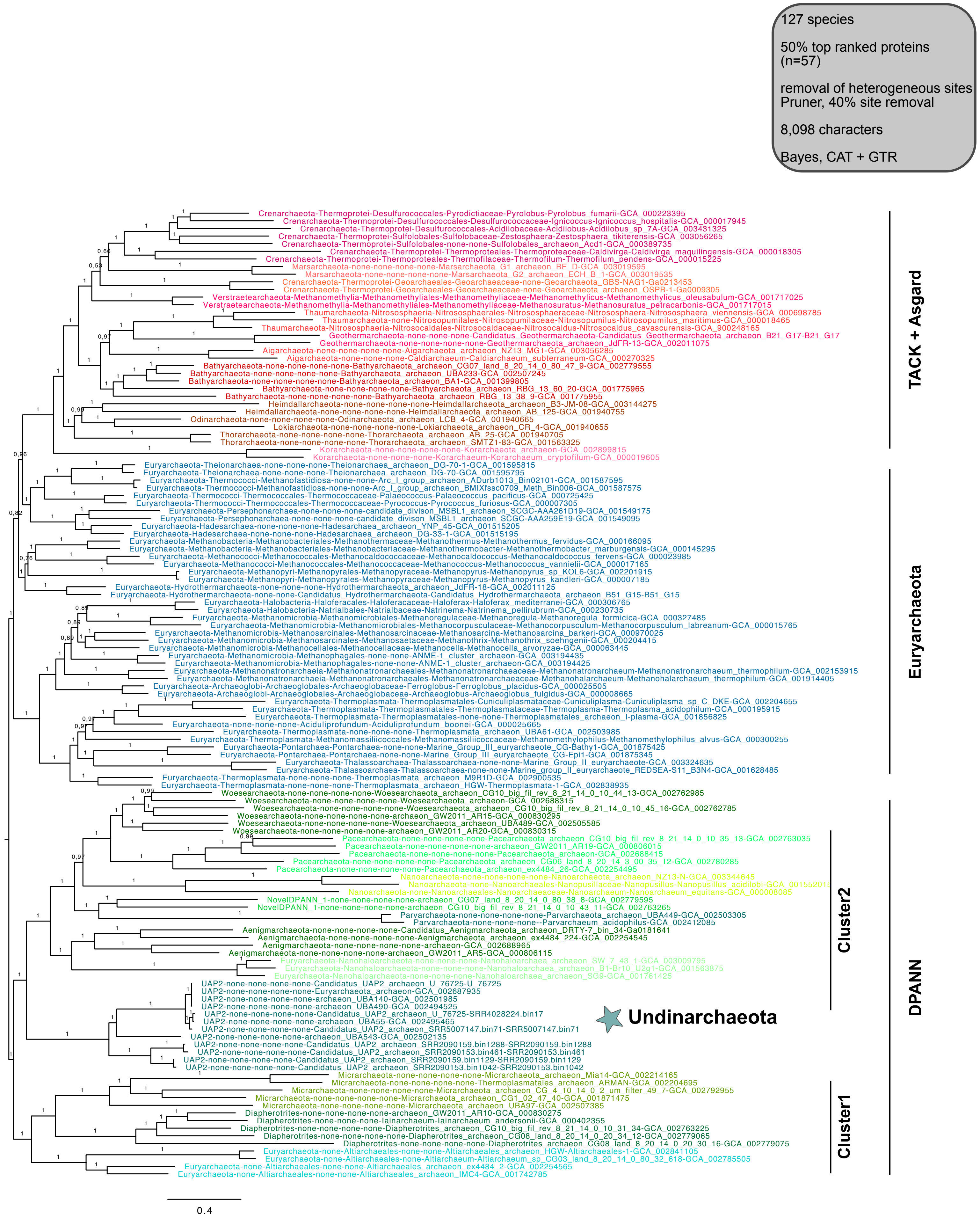


**Supplementary Figure 40 | Phylogenetic placement of Undinarchaeota based on an alignment generated with the 50% top ranked proteins (n=57) and the 127 species set.** 40% of heterogenous sites were removed from the alignment with the chi2 test. A Bayesian phylogenetic tree was inferred with the CAT+GTR model run with two chains for 6,788 cycles (25% burn-in). The tree was artificially rooted with the DPANN archaea and the grey arrow shows the root inferred with minimal ancestor deviation rooting (Tria et al., 2017). Scale bar: Average number of substitutions per site. Tree statistics for tree number 35 can be found in Supplementary Table 6.

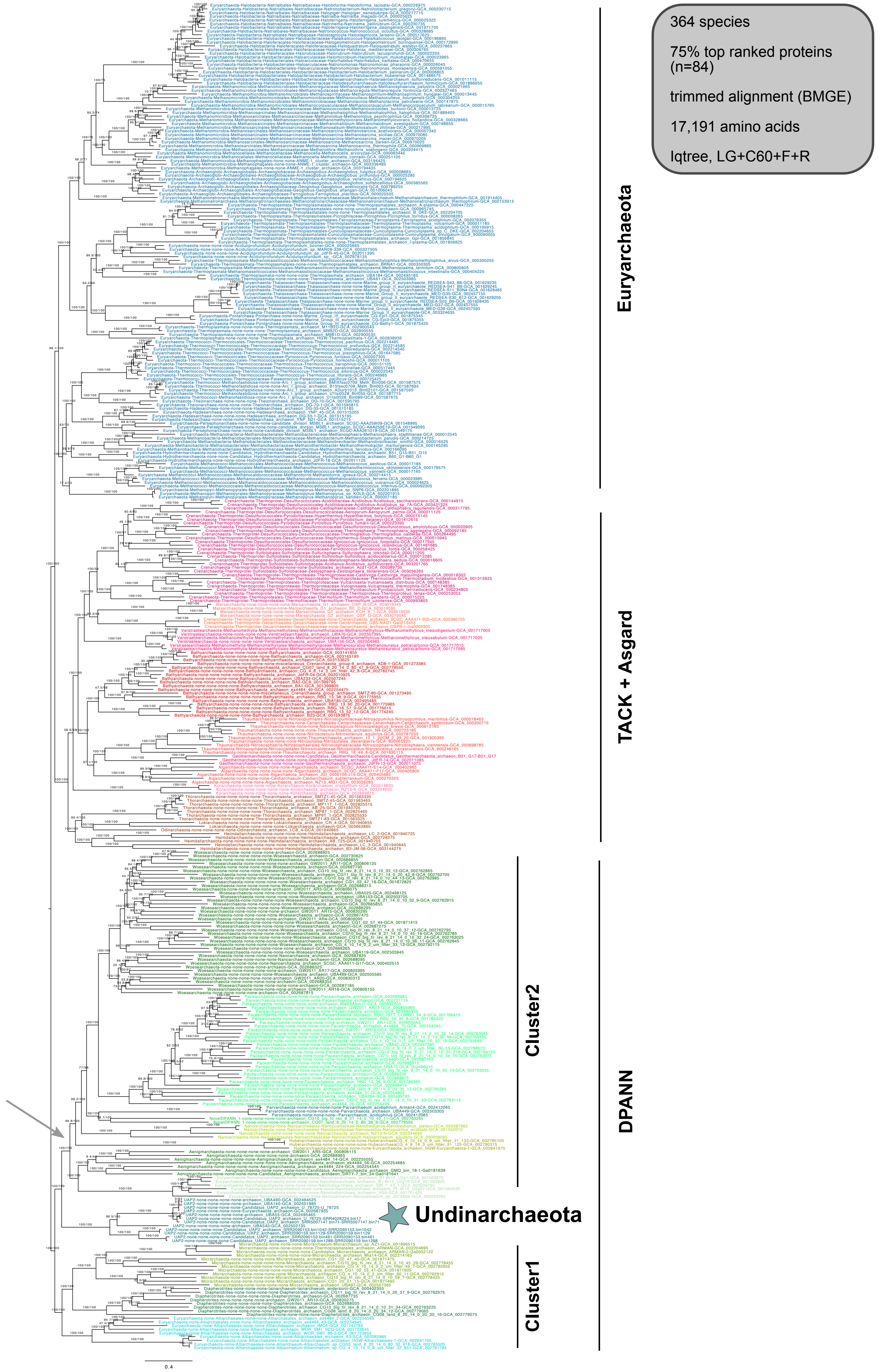


**Supplementary Figure 41 | Phylogenetic placement of Undinarchaeota based on an alignment generated with the 75% top ranked proteins (n=84) and the 364 species set.** The alignment was trimmed with BMGE (alignment length = 17,191 aa). A ML phylogenetic tree was inferred with the LG+C60+F+R model with an ultrafast bootstrap approximation (left) and SH-like approximate likelihood tests (right), each run with 1000 replicates. The tree was artificially rooted with the DPANN archaea and the grey arrow shows the root inferred with minimal ancestor deviation rooting (Tria et al., 2017). Scale bar: Average number of substitutions per site. Tree statistics for tree number 36 can be found in Supplementary Table 6.

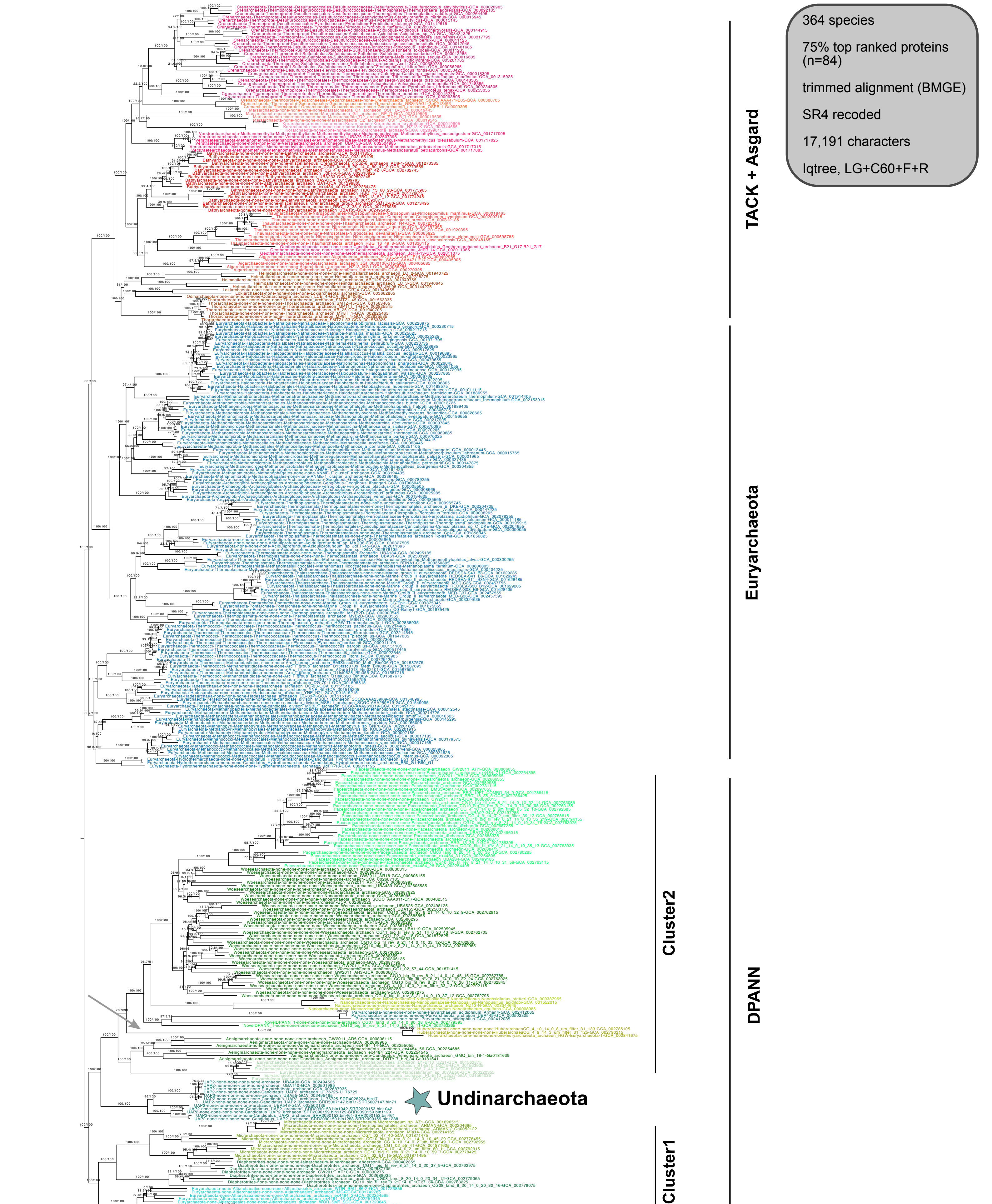


**Supplementary Figure 42 | Phylogenetic placement of Undinarchaeota based on an alignment generated with the 75% top ranked proteins (n=84) and the 364 species set.** The alignment was trimmed with BMGE and recoded into 4 character states (SR4 decoding; alignment length = 17,191 characters). A ML phylogenetic tree was inferred with the LG+C60+F+R model with an ultrafast bootstrap approximation (left) and SH-like approximate likelihood tests (right), each run with 1000 replicates. The tree was artificially rooted with the DPANN archaea and the grey arrow shows the root as inferred with minimal ancestor deviation rooting (Tria et al., 2017). Scale bar: Average number of substitutions per site. Tree statistics for tree number 37 can be found in Supplementary Table 6.

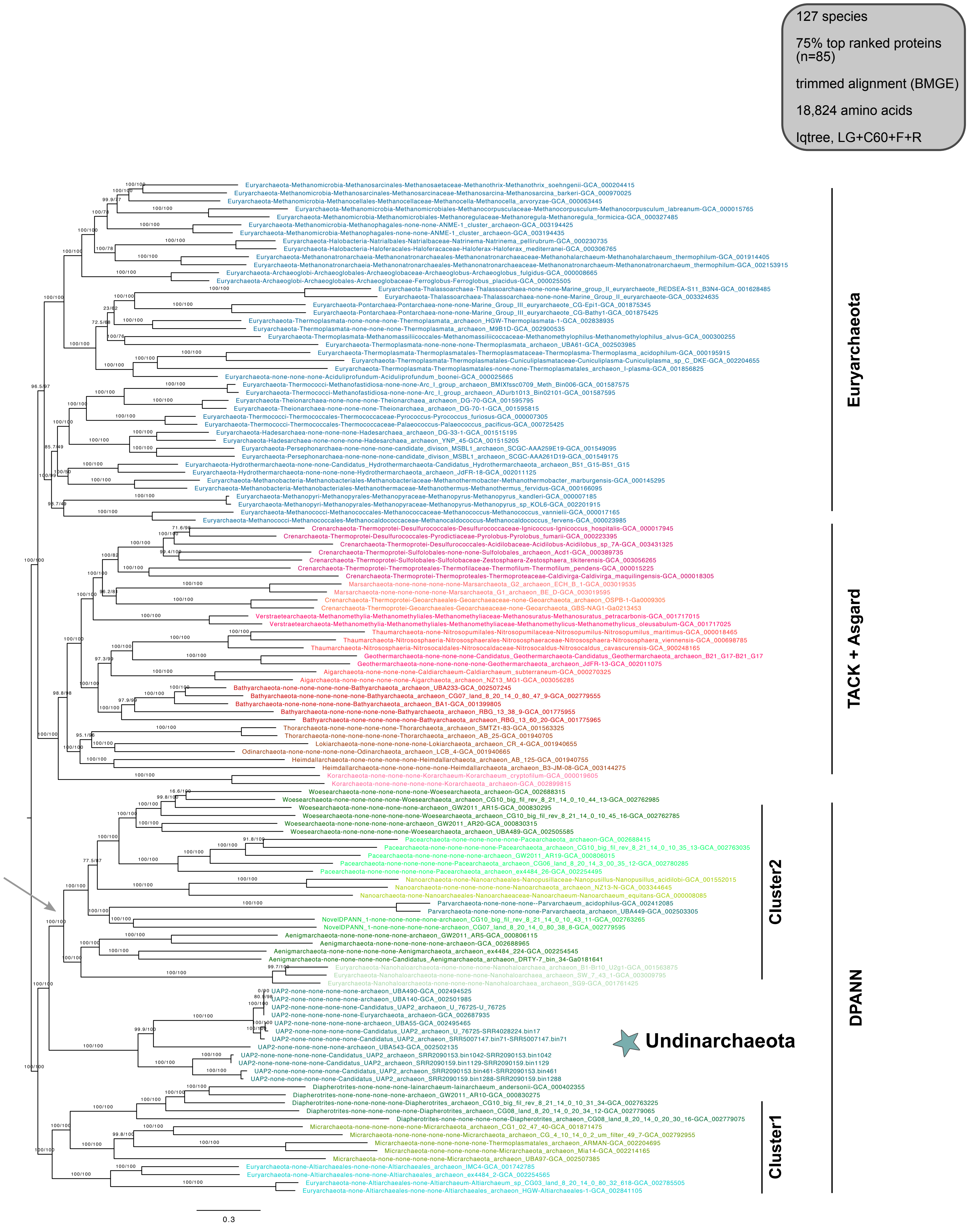


**Supplementary Figure 43 | Phylogenetic placement of Undinarchaeota based on an alignment generated with the 75% top ranked proteins (n=85) and the 127 species set.** The alignment was trimmed with BMGE (alignment length = 18,824 aa). A ML phylogenetic tree was inferred with the LG+C60+F+R model with an ultrafast bootstrap approximation (left) and SH-like approximate likelihood tests (right), each run with 1000 replicates. The tree was artificially rooted with the DPANN archaea and the grey arrow shows the root position inferred with minimal ancestor deviation rooting (Tria et al., 2017). Scale bar: Average number of substitutions per site. Tree statistics for tree number 38 can be found in Supplementary Table 6.

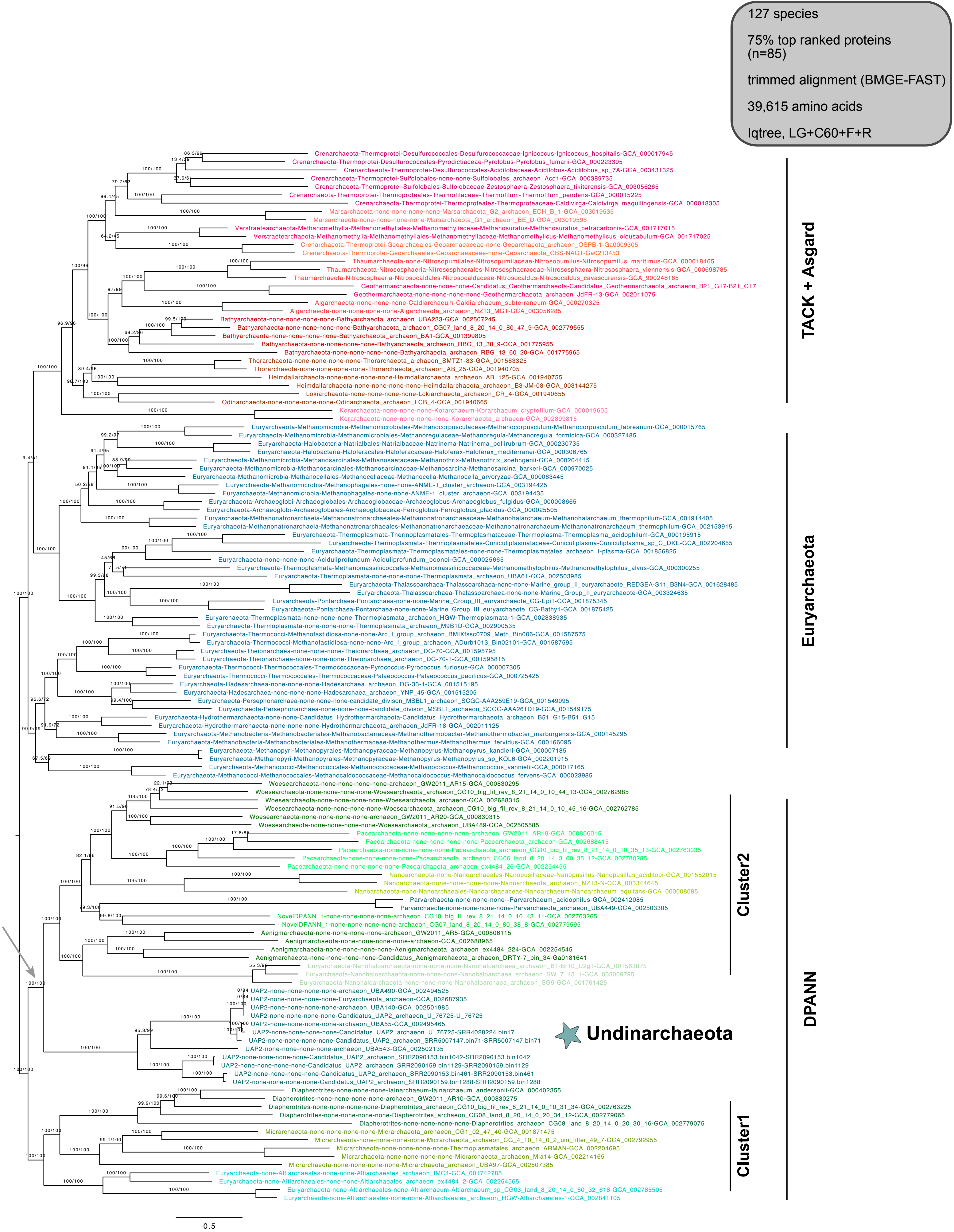


**Supplementary Figure 44 | Phylogenetic placement of Undinarchaeota based on an alignment generated with the 75% top ranked proteins (n=85) and the 127 species set.** The alignment was trimmed with BMGE-FAST (alignment length = 39,615 aa). A ML phylogenetic tree was inferred with the LG+C60+F+R model with an ultrafast bootstrap approximation (left) and SH-like approximate likelihood tests (right), each run with 1000 replicates. The tree was artificially rooted with the DPANN archaea and the grey arrow shows the root position inferred with minimal ancestor deviation rooting. Scale bar: Average number of substitutions per site. Tree statistics for tree number 39 can be found in Supplementary Table 6.

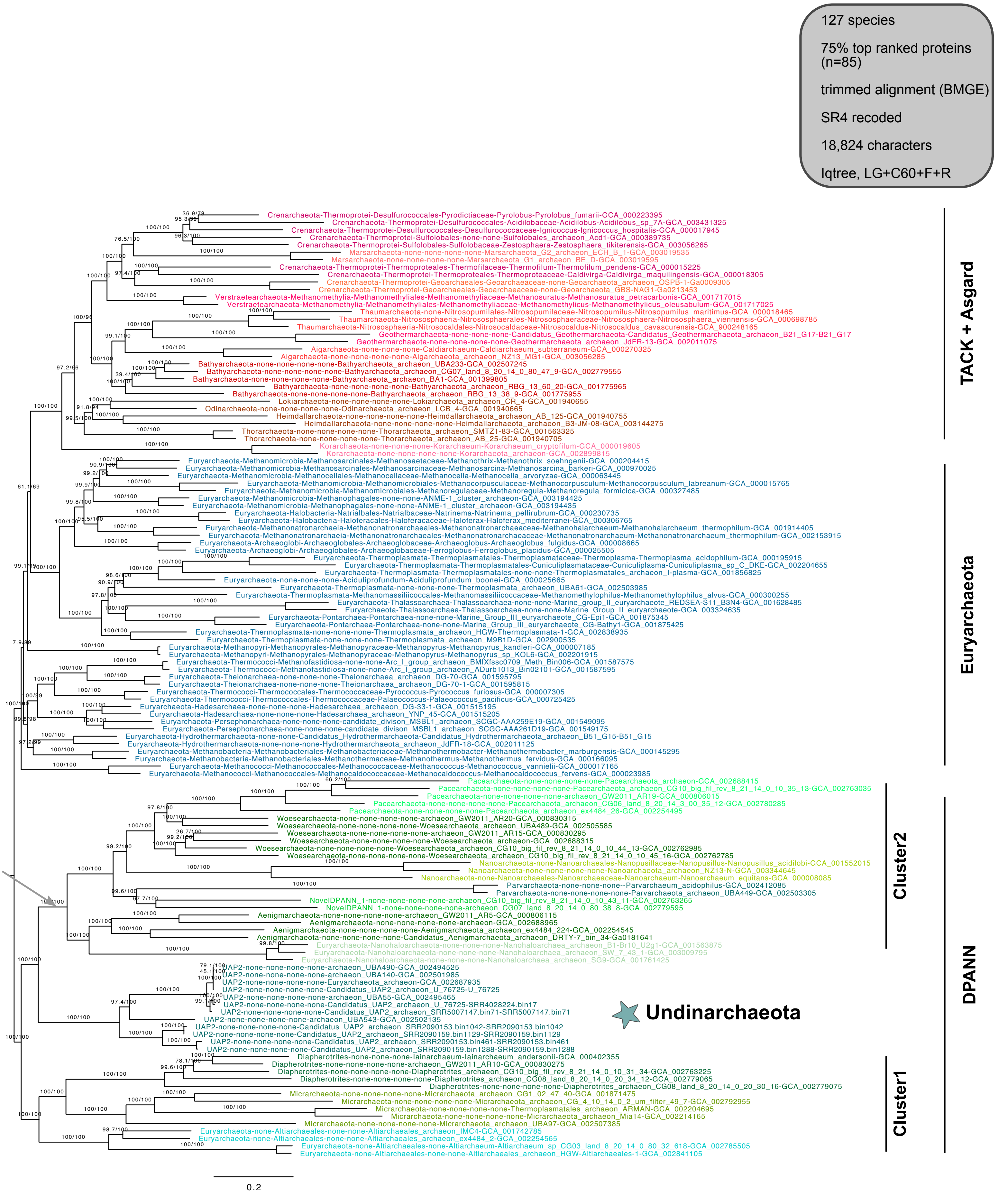


**Supplementary Figure 45 | Phylogenetic placement of Undinarchaeota based on an alignment generated with the 75% top ranked proteins (n=85) and the 127 species set.** The aligment was trimmed with BMGE and decoded into 4 character states (SR4 recoding; alignment length = 18,824 characters). A ML phylogenetic tree was inferred with the LG+C60+F+R model with an ultrafast bootstrap approximation (left) and SH-like approximate likelihood tests (right), each run with 1000 replicates. The tree was artificially rooted with the DPANN archaea and the grey arrow shows the root position inferred with minimal ancestor deviation rooting (Tria et al., 2017). Scale bar: Average number of substitutions per site. Tree statistics for tree number 40 can be found in Supplementary Table 6.

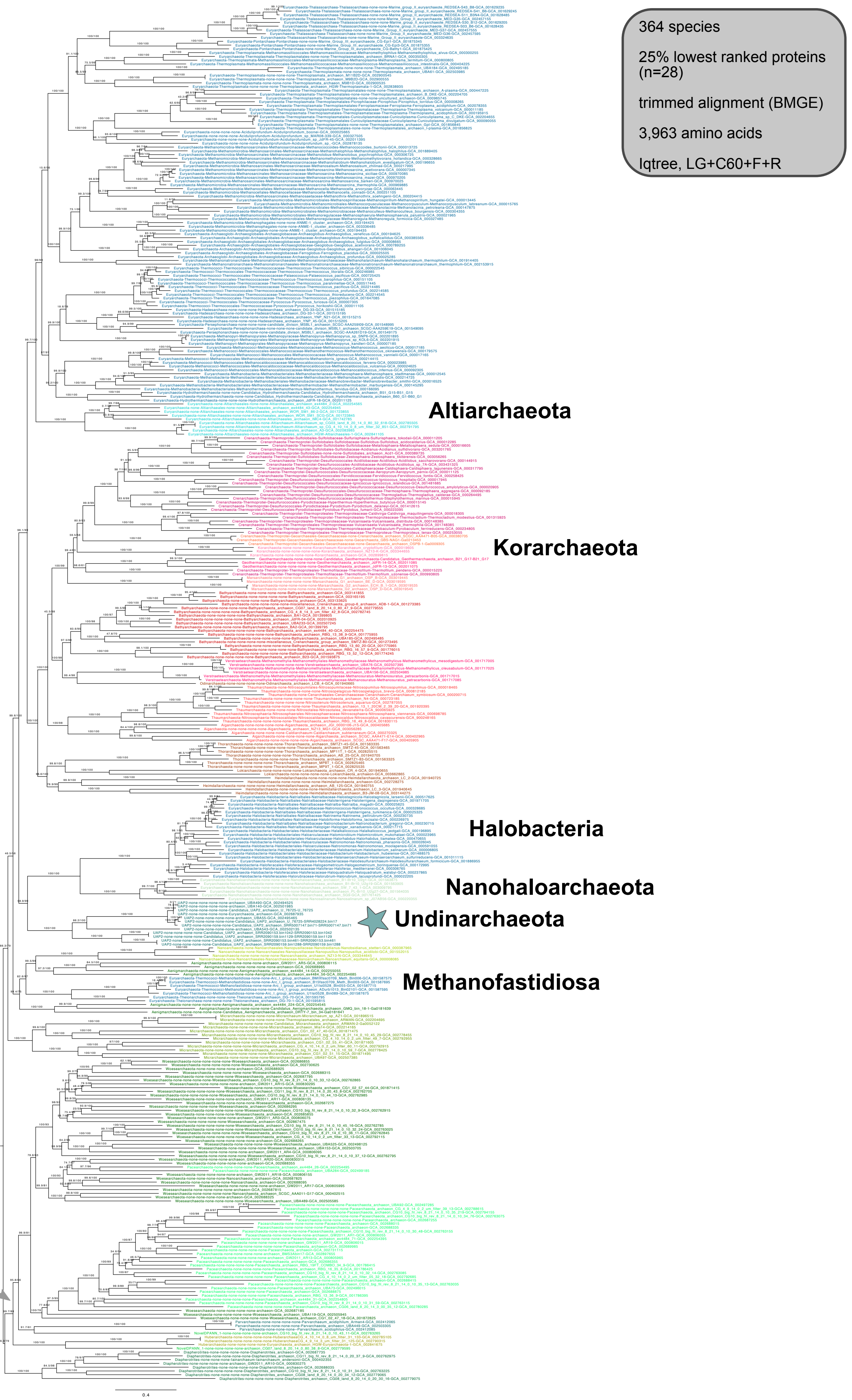


**Supplementary Figure 46 | Phylogenetic placement of Undinarchaeota based on an alignment generated with the 25% lowest ranking proteins (n=28) and the 364 species set.** The alignment was trimmed with BMGE (alignment length = 3,963 aa). A ML phylogenetic tree was inferred with the LG+C60+F+R model with an ultrafast bootstrap approximation (left) and SH-like approximate likelihood tests (right), each run with 1000 replicates. The tree was artificially rooted with the DPANN archaea and the grey arrow shows the root inferred with minimal ancestor deviation rooting (Tria et al., 2017). Scale bar: Average number of substitutions per site. Tree statistics for tree number 41 can be found in Supplementary Table 6.

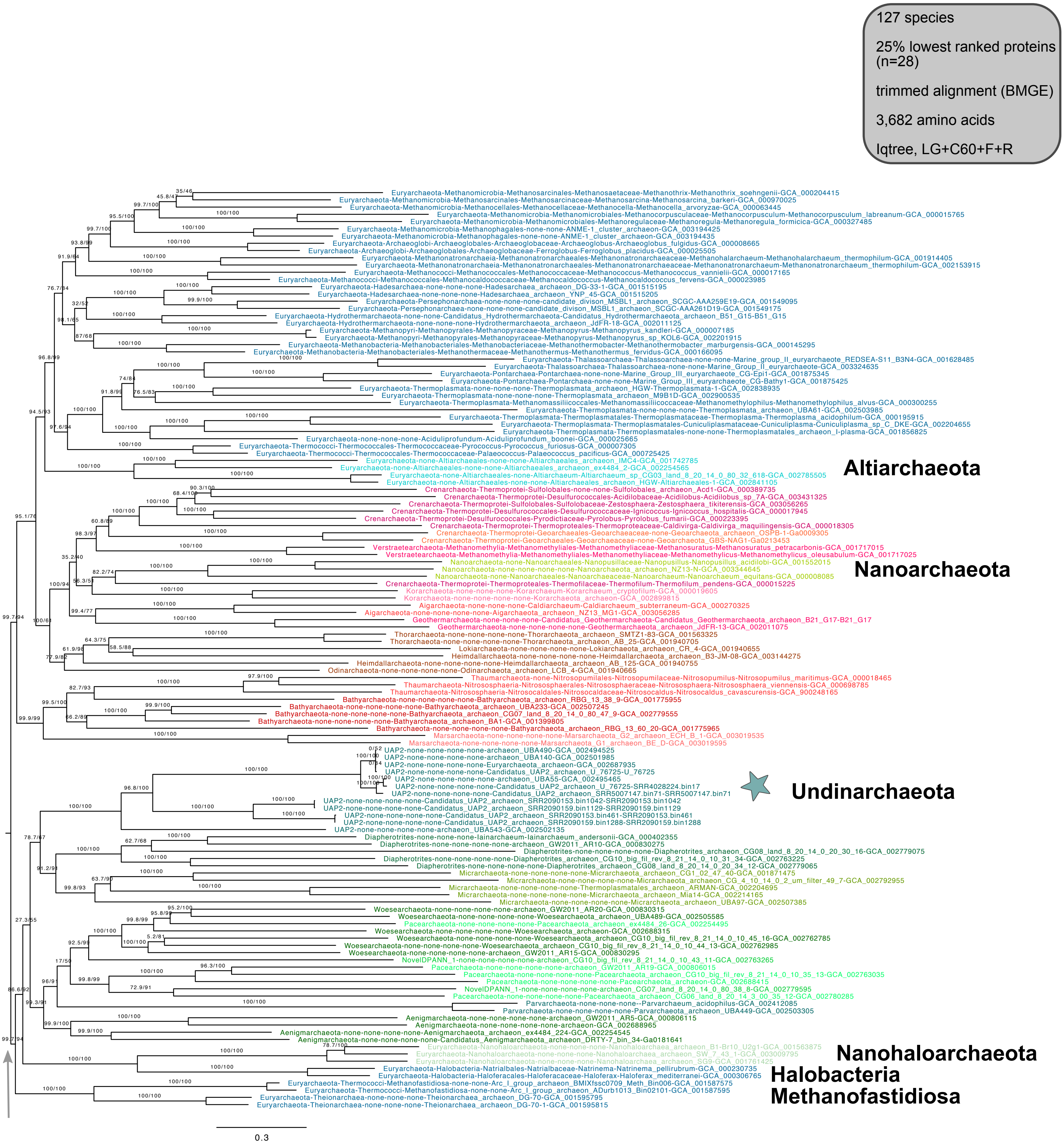


**Supplementary Figure 47 | Phylogenetic placement of Undinarchaeota based on an alignment generated with the 25% lowest ranked proteins (n=28) and the 127 species set.** The alignment was trimmed with BMGE (alignment length = 3,682 aa). A ML phylogenetic tree was inferred with the LG+C60+F+R model with an ultrafast bootstrap approximation (left) and SH-like approximate likelihood tests (right), each run with 1000 replicates. The tree was artificially rooted with the DPANN archaea and the grey arrow shows the root position inferred with minimal ancestor deviation rooting (Tria et al., 2017). Scale bar: Average number of substitutions per site. Tree statistics for tree number 42 can be found in Supplementary Table 6.

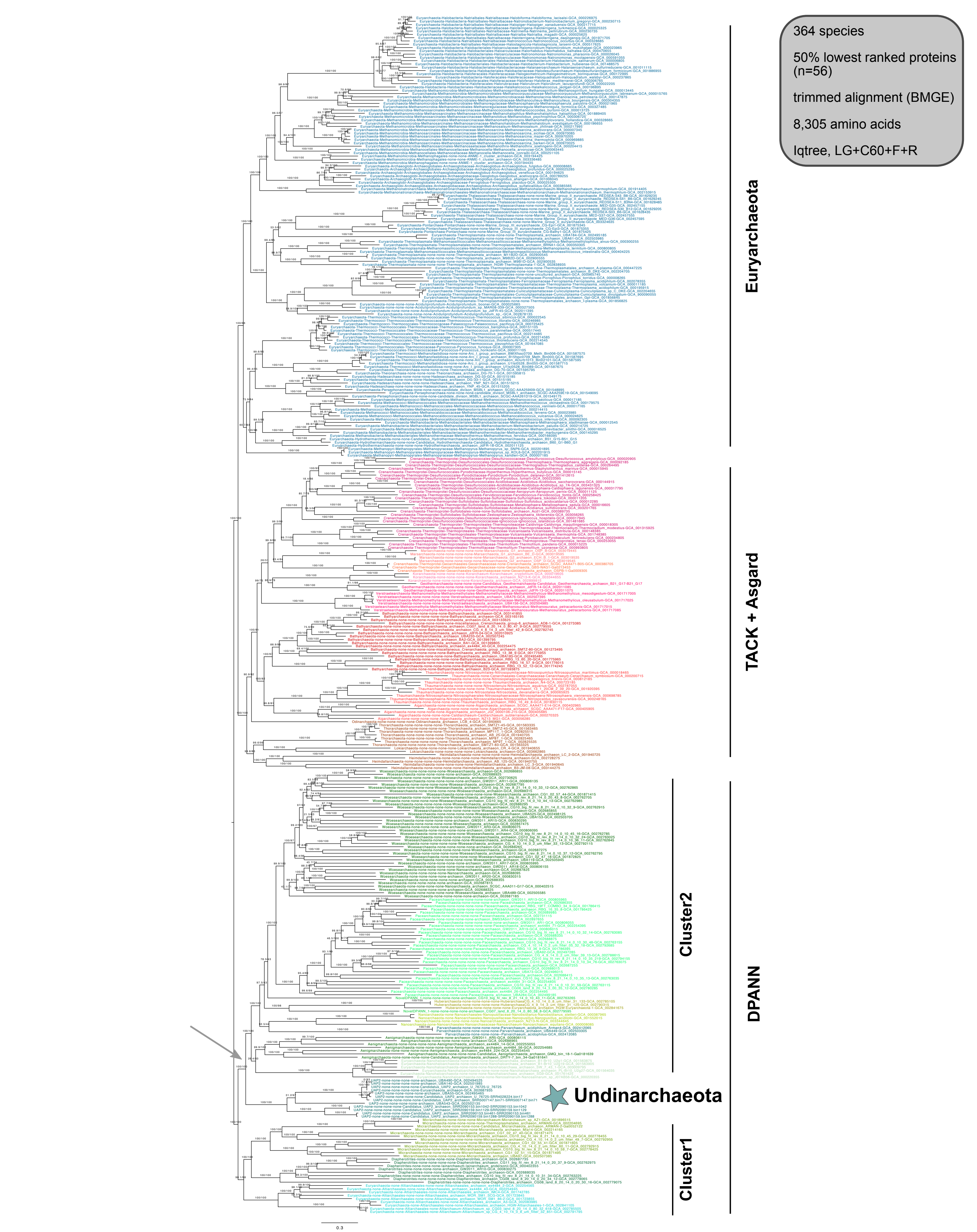


**Supplementary Figure 48 | Phylogenetic placement of Undinarchaeota based on an alignment generated with the 50% lowest ranked proteins (n=56) and the 364 species set.** The alignment was trimmed with BMGE (alignment length = 8,305 aa). A ML phylogenetic tree was inferred with the LG+C60+F+R model with an ultrafast bootstrap approximation (left) and SH-like approximate likelihood tests (right), each run with 1000 replicates. The tree was artificially rooted with the DPANN archaea and the grey arrow shows the root position inferred with minimal ancestor deviation rooting (Tria et al., 2017). Scale bar: Average number of substitutions per site. Tree statistics for tree number 43 can be found in Supplementary Table 6.

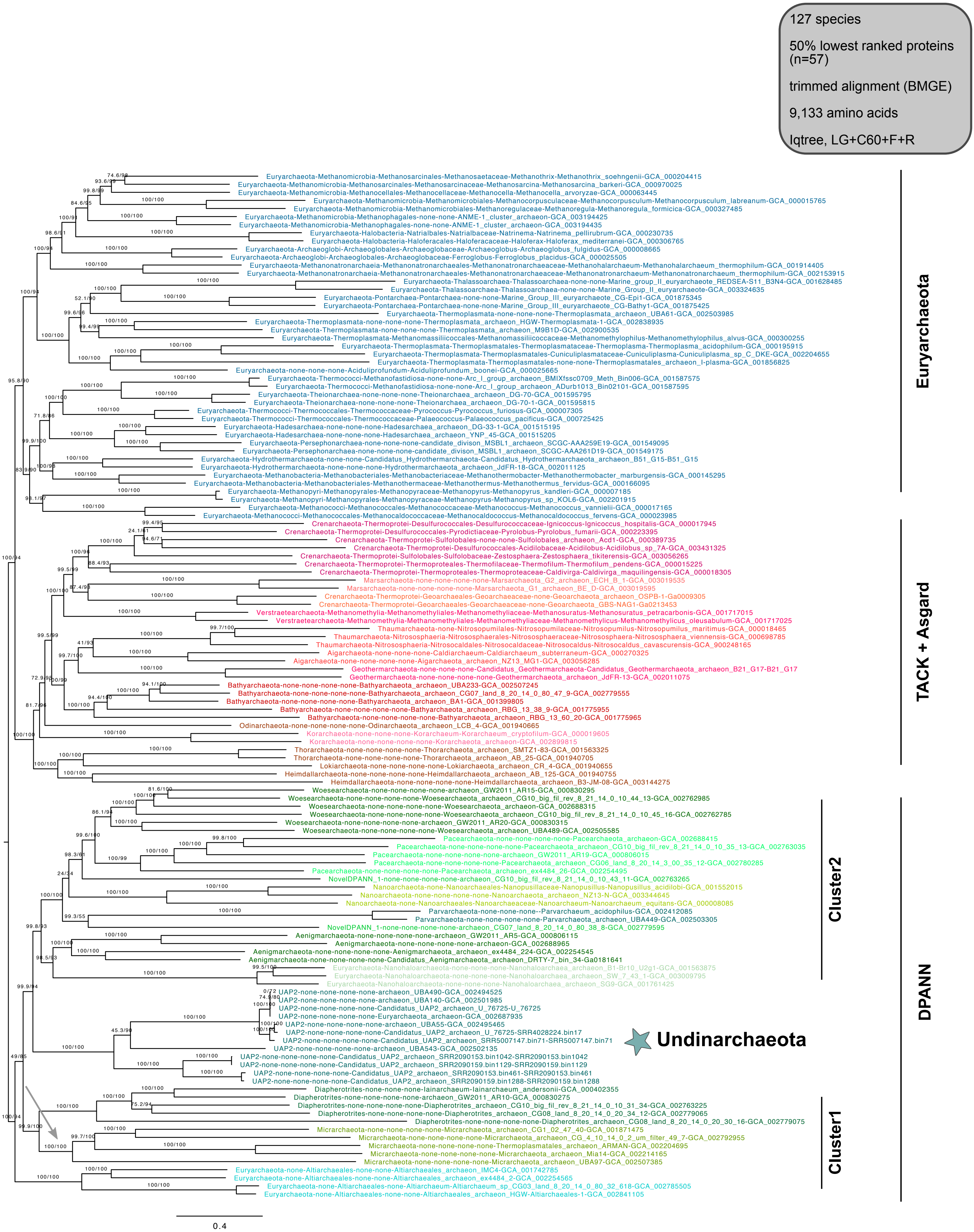


**Supplementary Figure 49 | Phylogenetic placement of Undinarchaeota based on an alignment generated with the 50% lowest ranked proteins (n=57) and the 127 species set.** The alignment was trimmed with BMGE (alignment length = 9,133 aa). A ML phylogenetic tree was inferred with the LG+C60+F+R model with an ultrafast bootstrap approximation (left) and SH-like approximate likelihood tests (right), each run with 1000 replicates. The tree was artificially rooted with the DPANN archaea and the grey arrow shows the root inferred with minimal ancestor deviation rooting (Tria et al., 2017). Scale bar: Average number of substitutions per site. Tree statistics for tree number 44 can be found in Supplementary Table 6.

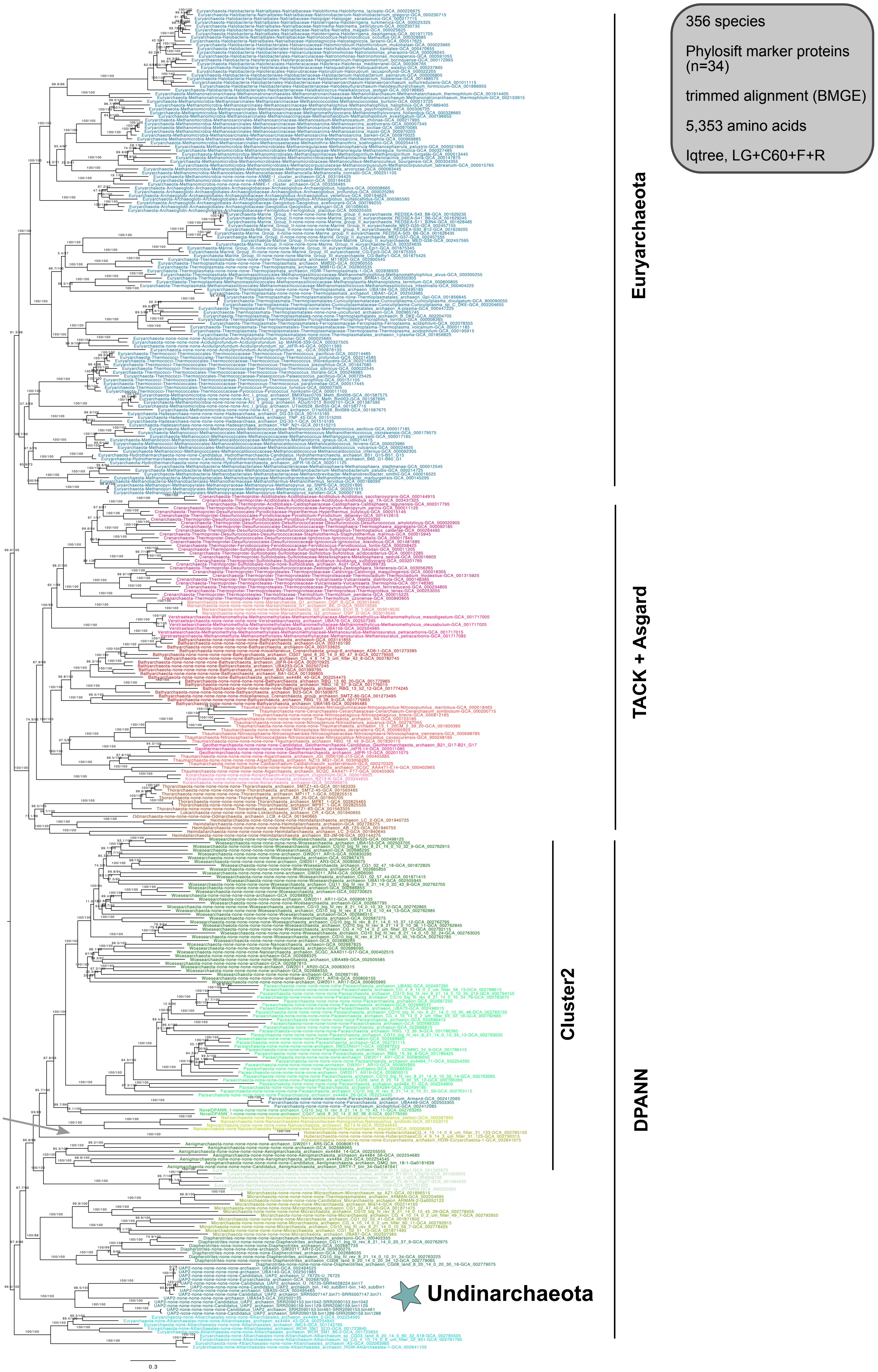


**Supplementary Figure 50 | Phylogenetic placement of Undinarchaeota based on an alignment generated with the phylosift marker proteins (n=34) and the 356 species set.** The alignment was trimmed with BMGE (alignment length = 5,353 aa). A ML phylogenetic tree was inferred with the LG+C60+F+R model with an ultrafast bootstrap approximation (left) and SH-like approximate likelihood tests (right), each run with 1000 replicates. The tree was artificially rooted with the DPANN archaea and the grey arrow shows the root position inferred with minimal ancestor devation rooting (Tria et al., 2017). Scale bar: Average number of substitutions per site. Tree statistics for tree number 45 can be found in Supplementary Table 6.

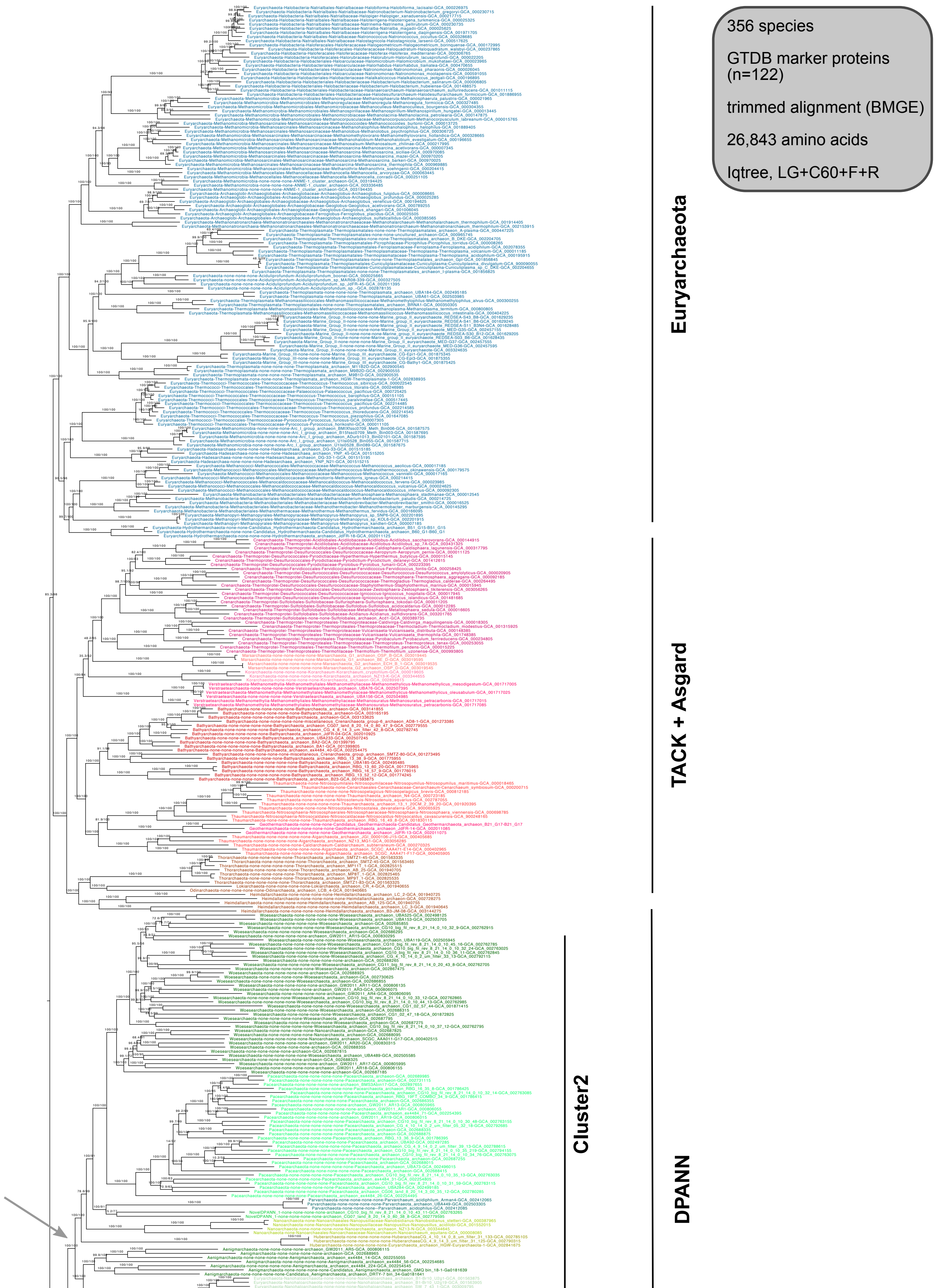


**Supplementary Figure 51 | Phylogenetic placement of Undinarchaeota based on an alignment generated with the GTDB archaeal marker proteins (n=122) and the 356 species set.** The alignment was trimmed with BMGE (alignment length = 26,843 aa). A ML phylogenetic tree was inferred with the LG+C60+F+R model using iqtree with an ultrafast bootstrap approximation (left) and SH-like approximate likelihood tests (right), each run with 1000 replicates. The tree was artificially rooted with the DPANN archaea and the grey arrow shows the root position inferred with minimal ancestor deviation rooting (Tria et al., 2017). Scale bar: Average number of substitutions per site. Tree statistics for tree number 46 can be found in Supplementary Table 6.

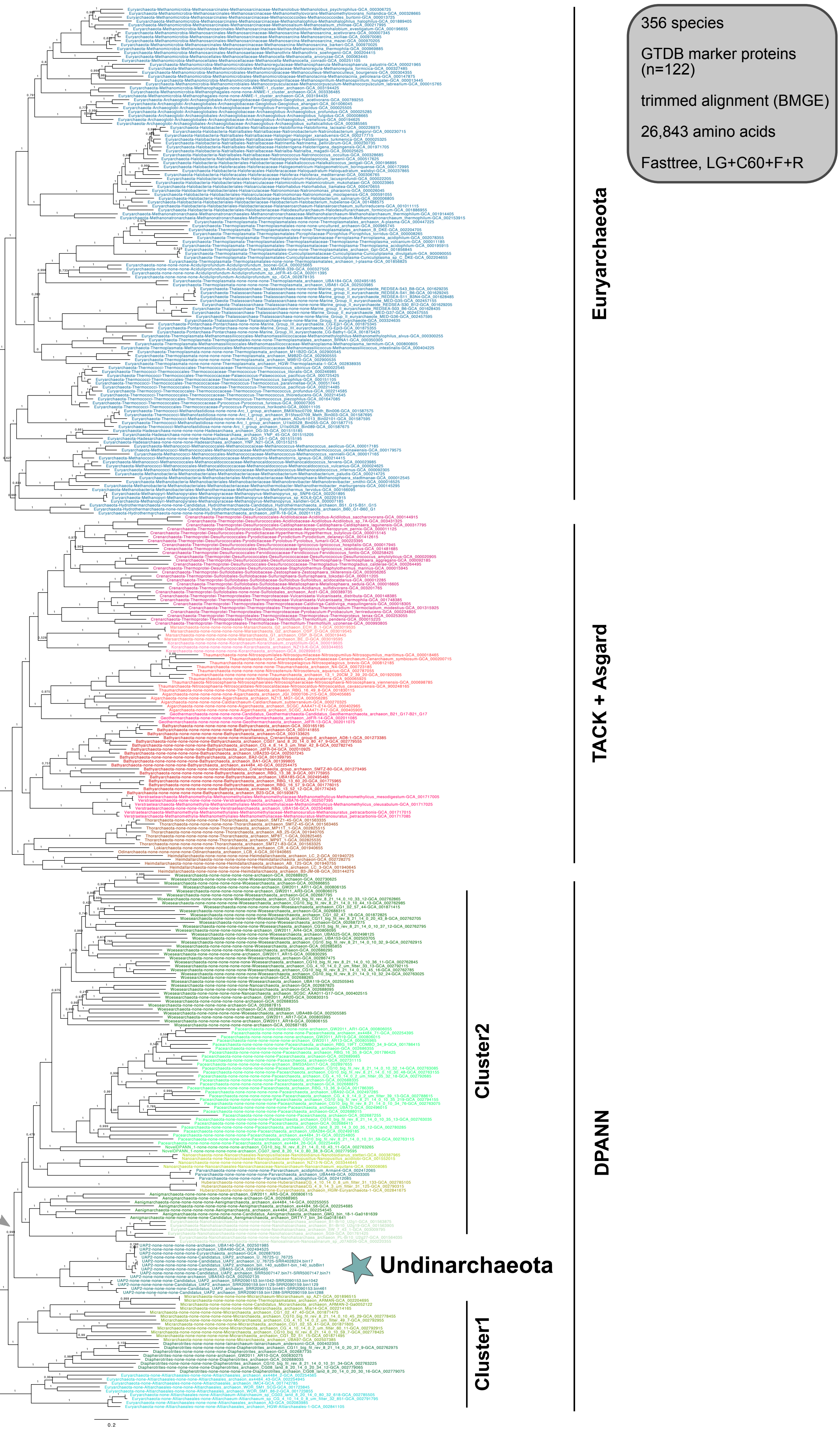


**Supplementary Figure 52 | Phylogenetic placement of Undinarchaeota based on an alignment generated with the archaeal GTDB marker proteins (n=122) and the 356 species set.** The alignment was trimmed using BMGE (alignment length = 26,843 aa). An approximately-ML phylogenetic tree was inferred with the WAG+GAMMA model using fasttree with SH-like approximate likelihood tests run with 1000 replicates. The tree was artificially rooted with the DPANN archaea and the grey arrow shows the root inferred with minimal ancestor deviation rooting (Tria et al., 2017). Scale bar: Average number of substitutions per site. Tree statistics for tree number 47 can be found in Supplementary Table 6.

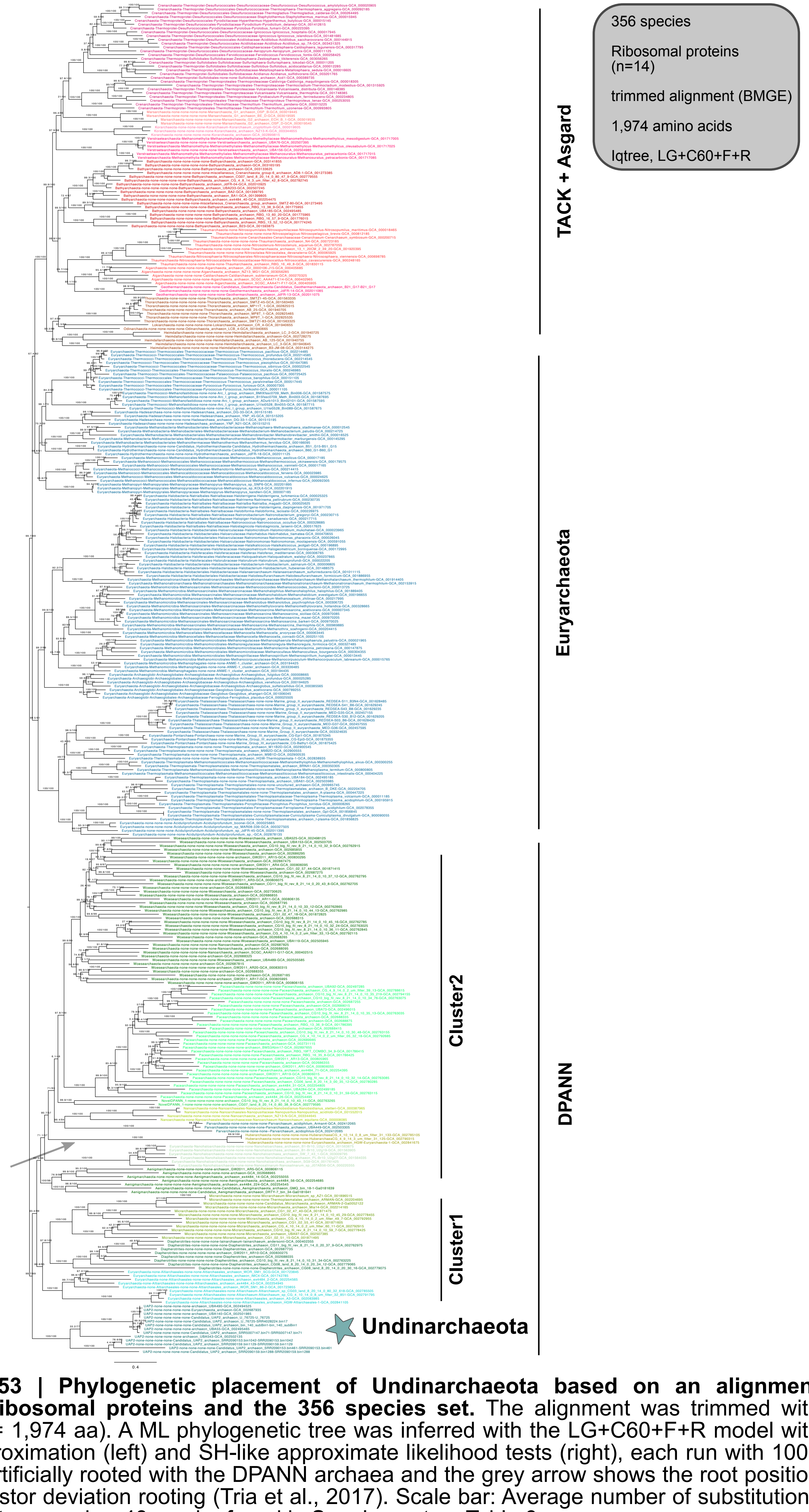


**Supplementary Figure 53 | Phylogenetic placement of Undinarchaeota based on an alignment generated with the 14 ribosomal proteins and the 356 species set.** The alignment was trimmed with BMGE (alignment length = 1,974 aa). A ML phylogenetic tree was inferred with the LG+C60+F+R model with an ultrafast bootstrap approximation (left) and SH-like approximate likelihood tests (right), each run with 1000 replicates. The tree was artificially rooted with the DPANN archaea and the grey arrow shows the root position inferred with minimal ancestor deviation rooting (Tria et al., 2017). Scale bar: Average number of substitutions per site. Tree statistics for tree number 48 can be found in Supplementary Table 6.

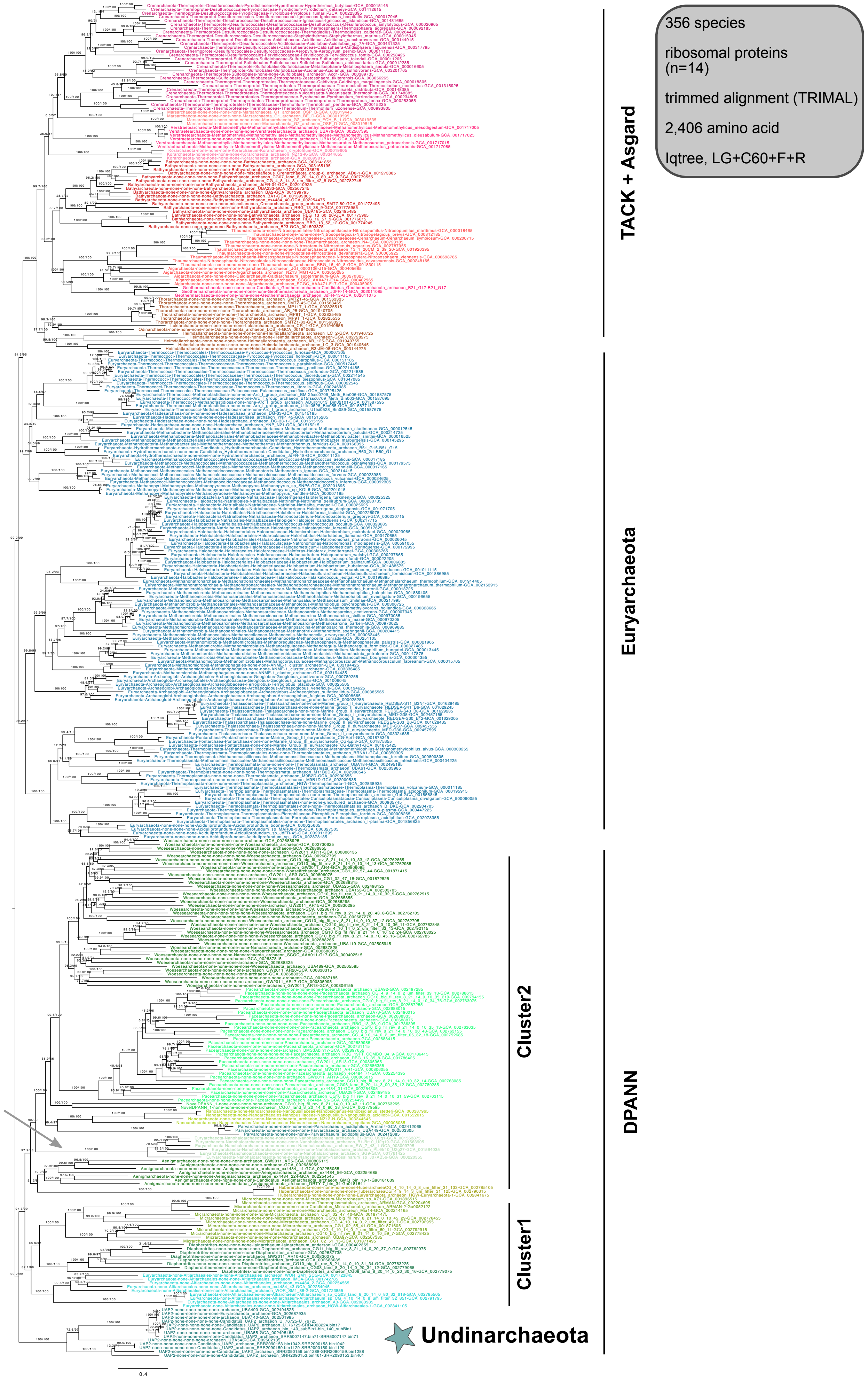


**Supplementary Figure 54 | Phylogenetic placement Undinarchaeota lineage based on an alignment generated with 14 ribosomal proteins and the 356 species set.** The alignment was trimmed with TRIMAL (alignment length = 2,406 aa). A ML phylogenetic tree was inferred with the LG+C60+F+R model with an ultrafast bootstrap approximation (left) and SH-like approximate likelihood tests (right), each run with 1000 replicates. The tree was artificially rooted with the DPANN archaea and the grey arrow shows the root position inferred with minimal ancestor deviation rooting (Tria et al., 2017). Scale bar: Average number of substitutions per site. Tree statistics for tree number 49 can be found in Supplementary Table 6.

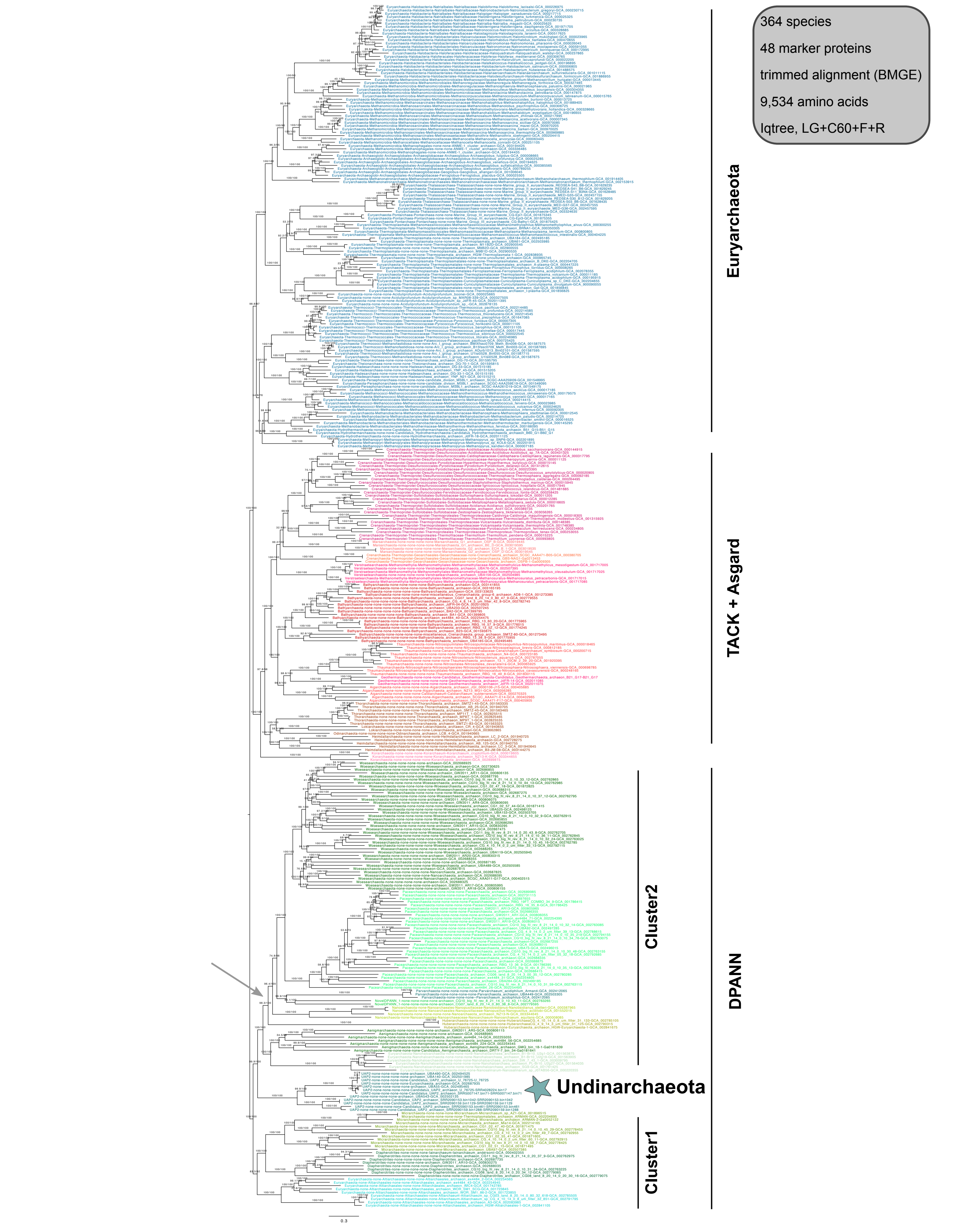


**Supplementary Figure 55 | Phylogenetic placement of Undinarchaeota based on an alignment generated with 48 universal marker proteins and the 364 species set.** The alignment was trimmed with BMGE (alignment length = 9,534 aa). A ML phylogenetic tree was inferred with the LG+C60+F+R model with an ultrafast bootstrap approximation (left) and SH-like approximate likelihood tests (right), each run with 1000 replicates. The tree was artificially rooted with the DPANN archaea and the grey arrow shows the root position inferred with minimal ancestor rooting (Tria et al., 2017). Scale bar: Average number of substitutions per site. Tree statistics for tree number 50 can be found in Supplementary Table 6.

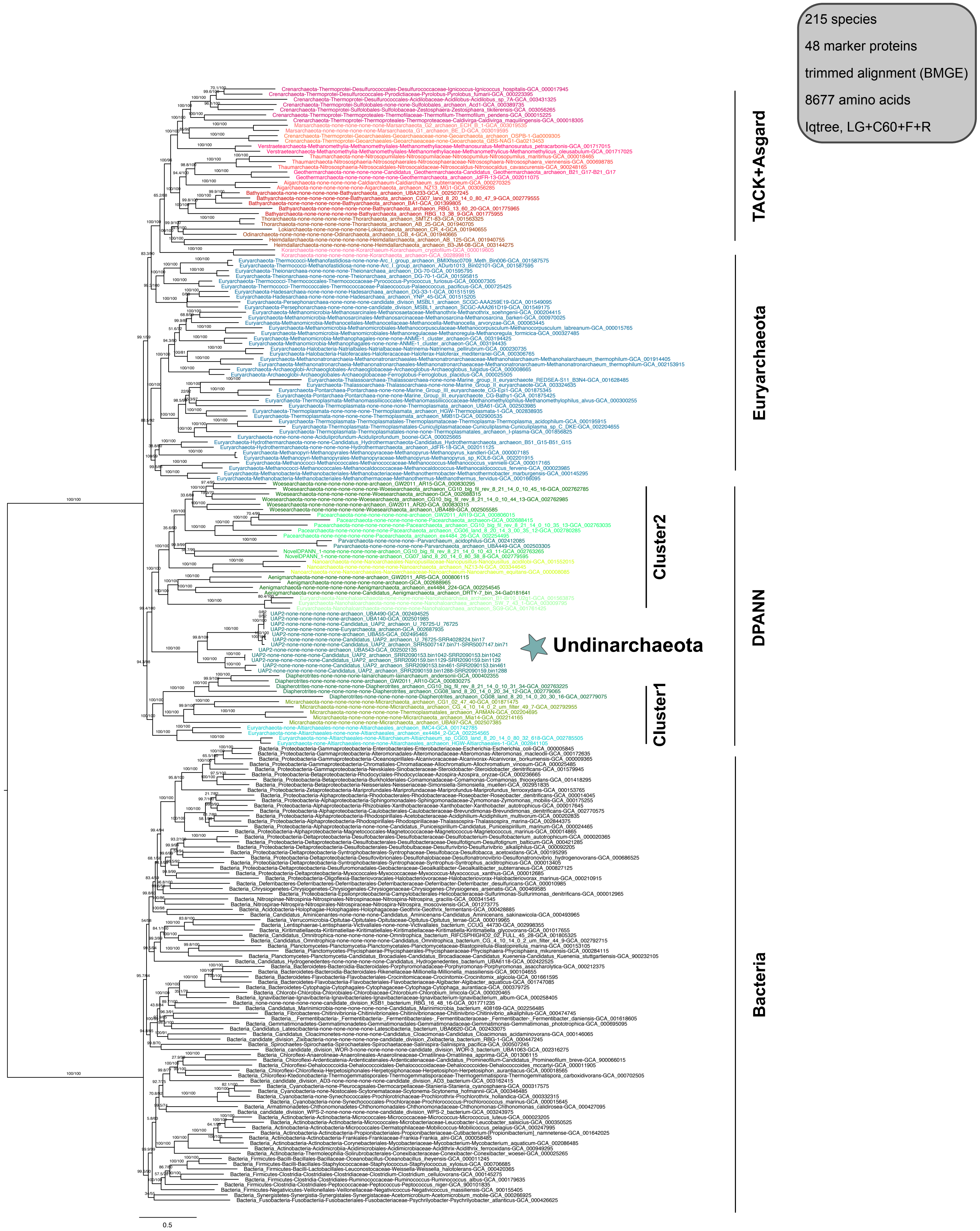


**Supplementary Figure 56 | Phylogenetic placement of Undinarchaeota based on an alignment generated with 48 universal marker proteins and the 127 archaeal + 88 bacterial species set.** The alignment was trimmed with BMGE (alignment length = 8,677 aa). A ML phylogenetic tree was inferred with the LG+C60+F+R model with an ultrafast bootstrap approximation (left) and SH-like approximate likelihood tests (right), each run with 1000 replicates. The tree was rooted using bacteria (black labels) as outgroup. Scale bar: Average number of substitutions per site. Tree statistics for tree number 51 can be found in Supplementary Table 6.

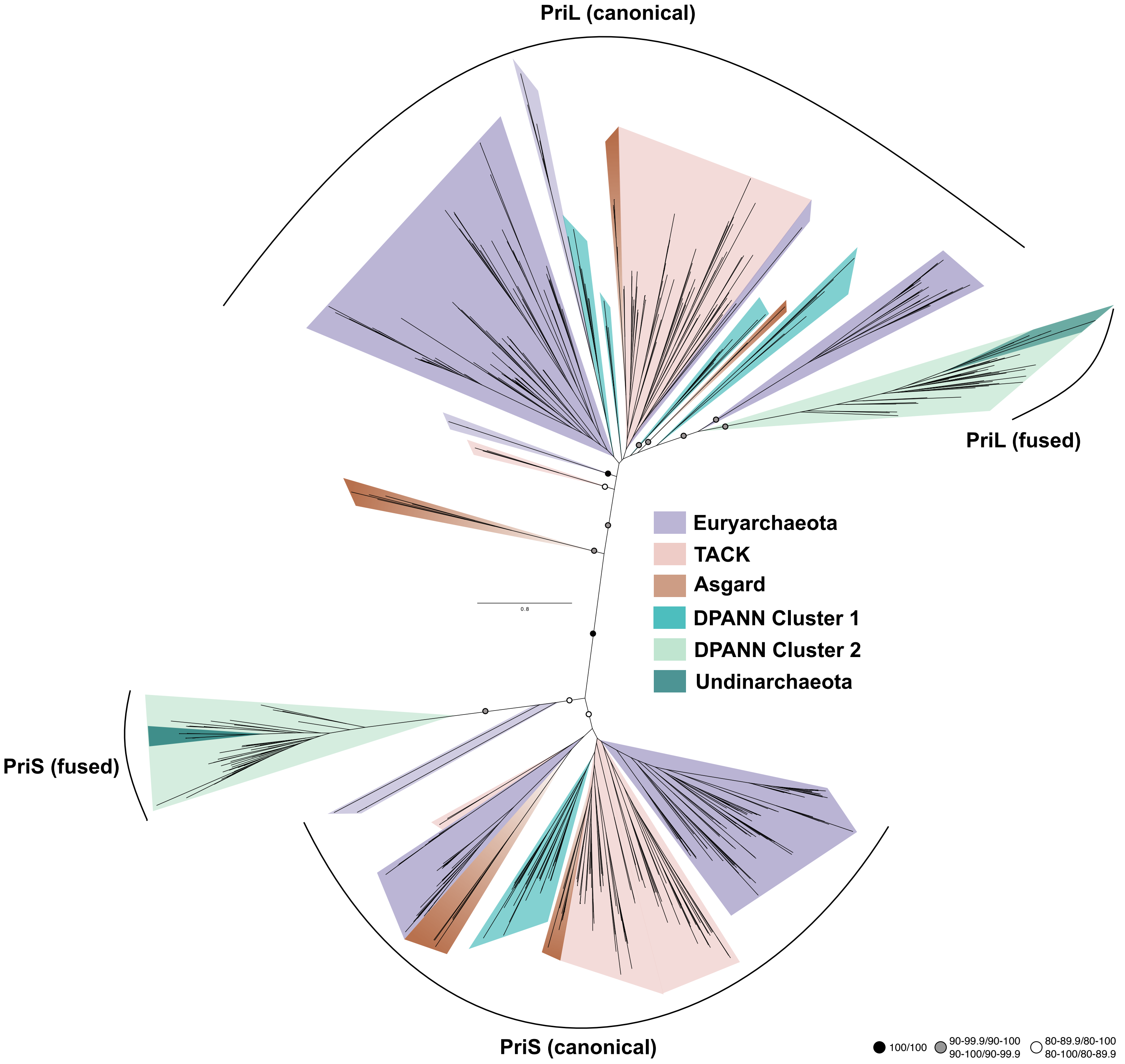


**Supplementary Figure 57 | Phylogenetic history of the primase subunits PriS and PriL in archaea.** An alignment was generated for all PriS and PriL sequences found in 364 archaea that was trimmed using TRIMAL (alignment length = 512 amino acids). The canonical PriS and PriL genes are encoded by two genes with the exception for most DPANN archaea that encode a fused version of the primase (Supplementary Table 10). These fused versions were split before aligning all sequences (n=585 sequences; see Methods for details). A maximum-likelihood phylogenetic tree was inferred with the LG+F+C10 model with an ultrafast bootstrap approximation (left) and SH-like approximate likelihood tests (right), each run with 1000 replicates. Bootstrap values above certain thresholds are indicated with colored circles. Scale bar: Average number of substitutions per site.

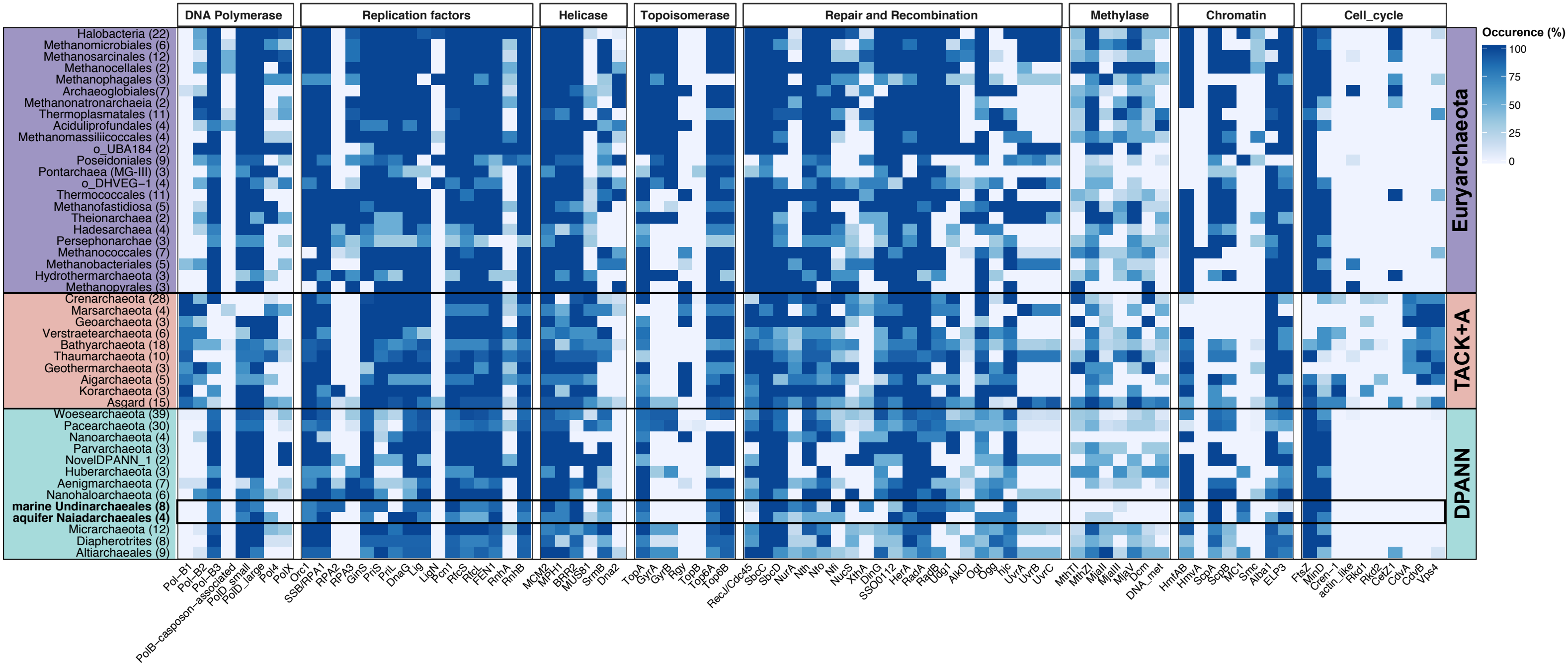


**Supplementary Figure 58 | Presence of key replication proteins across major archaeal lineages**. Heatmap of presence/absence patterns of key proteins are summarized across the total number of genomes included in each phylogenetic cluster (shown in percent). TACK + A = TACK + Asgard. Number in brackets = number of genomes analyzed for each phylogenetic cluster. Supplementary Table 19 lists the proteins used to generate the plot and Supplementary Table 9 lists the raw values.

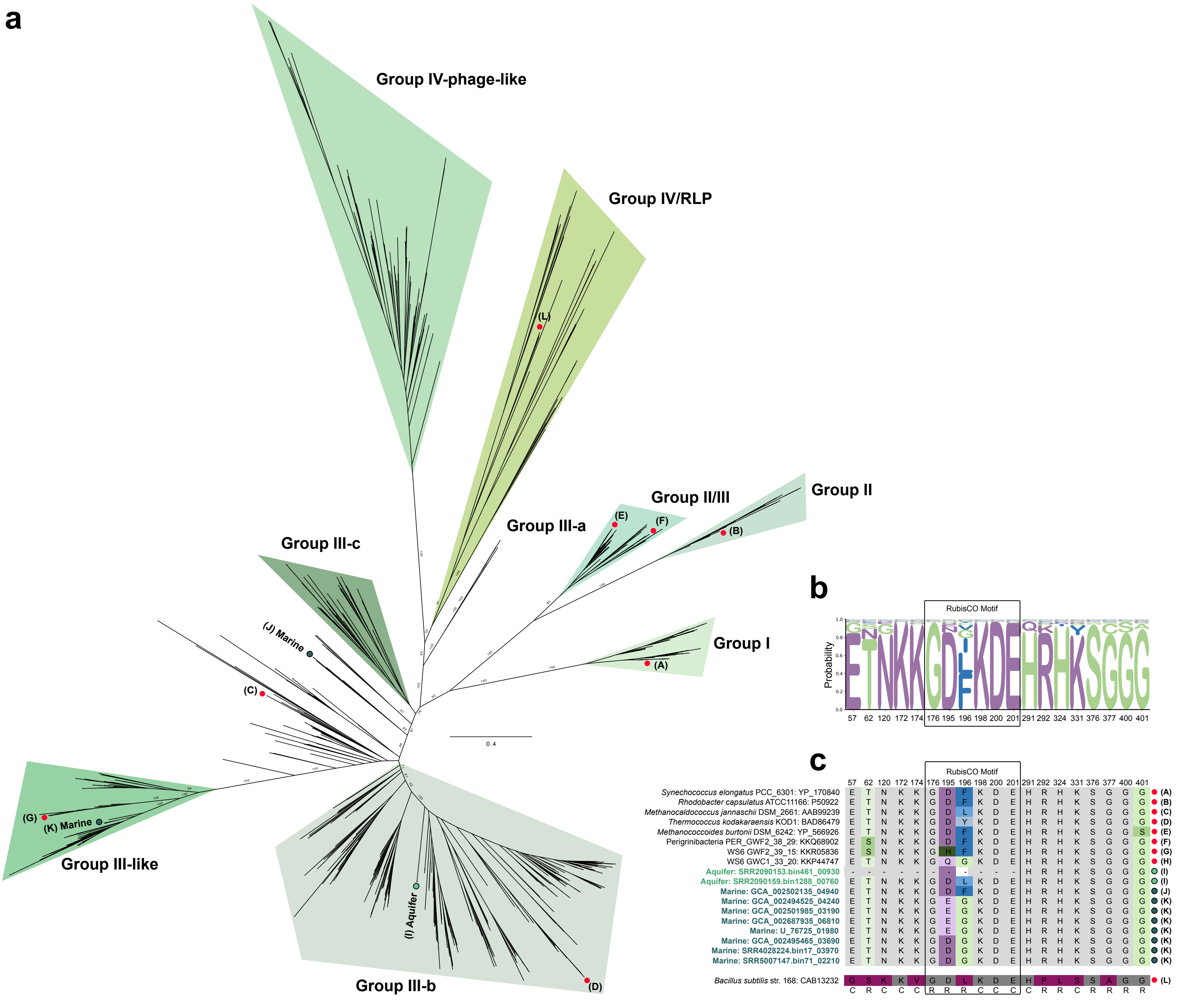


**Supplementary Figure 59 | Diversity of RubisCO proteins in archaea. a**, ML phylogenetic analysis of the RubisCO protein extracted from marine Undinarchaeales (dark green) and aquifer Naiadarchaeales MAGs (light green) that were added to an alignment from Jaffe et al., 2019 (n=786 sequences). The alignment was trimmed using BMGE (alignment length = 397 aa). A ML phylogenetic tree was inferred with the LG+G model with an ultrafast bootstrap approximation with 1000 replicates. Scale bar: Average number of substitutions per site. **b**, Probability plot of the occurence of each amino acid of the catalytic site of the RubisCO protein across 786 sequences. Color-coding is based on the hydrophobicity score. **c**, Conservation of the catalytic site of selected amino acid sequences of the catalytic site compared to the reference sequence of *Synechococcus elongatus* PCC_6301 as described by Jaffe et al., 2019. Differences in amino acid sequence compared to the reference are colored based on their hydrophobicity score and conserved sites are colored in grey. The position of each amino acid in the alignment is indicated at the top of the scheme and information on whether an amino acid represents a catalytic site (C) or RubisCO binding site (R) is indicated at the bottom. (A)-(L) = Reference sequences for different RubisCO groups.

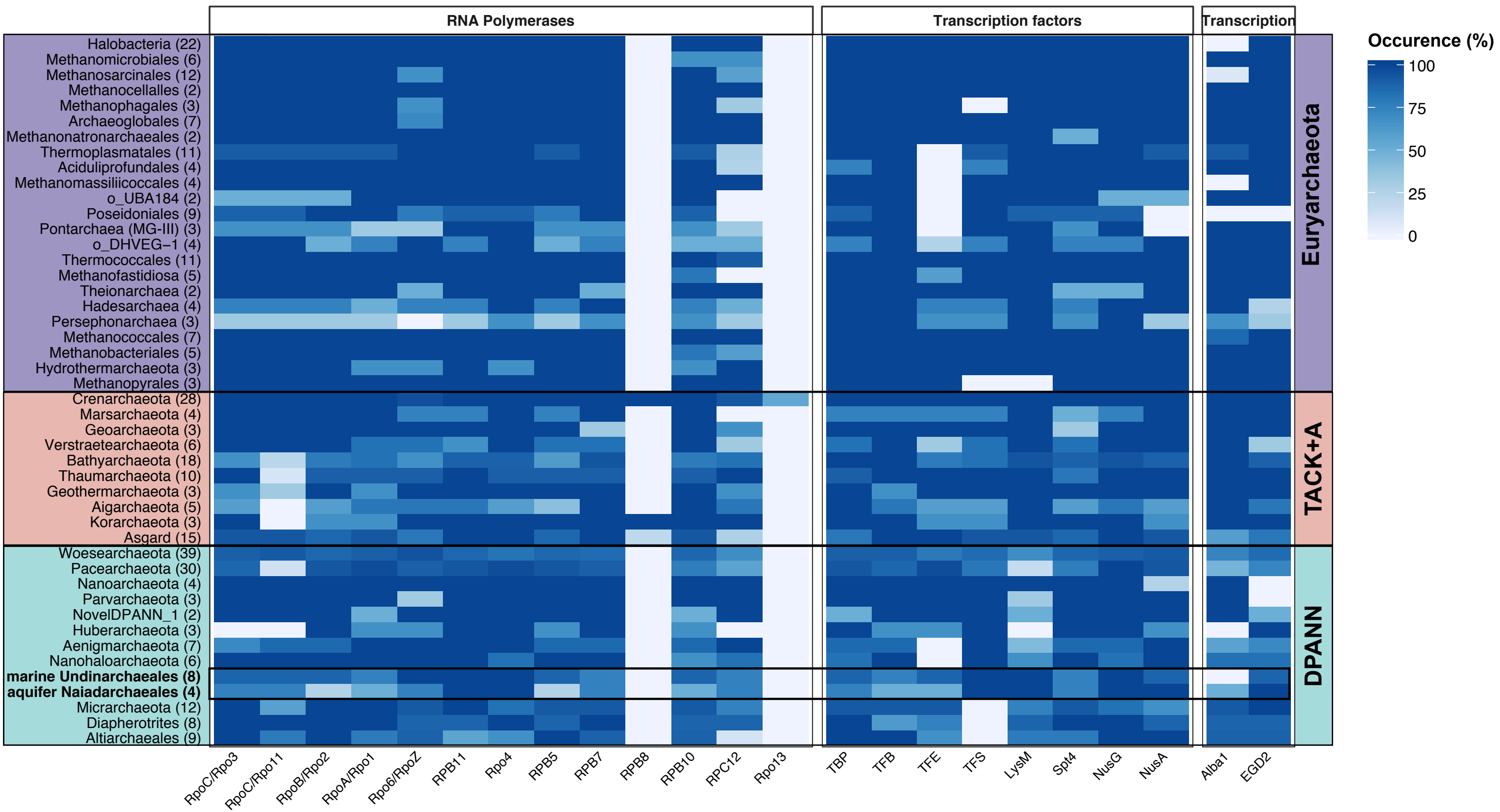


**Supplementary Figure 60 | Presence of key transcription-related proteins across major archaeal lineages**. Heatmap of presence/absence patterns of key proteins are summarized across the total number of genomes included in each phylogenetic cluster (shown in percent). TACK + A = TACK + Asgard. Number in brackets = number of genomes analyzed for each phylogenetic cluster. Supplementary Table 19 lists the proteins used to generate the plot and Supplementary Table 9 lists the raw values.

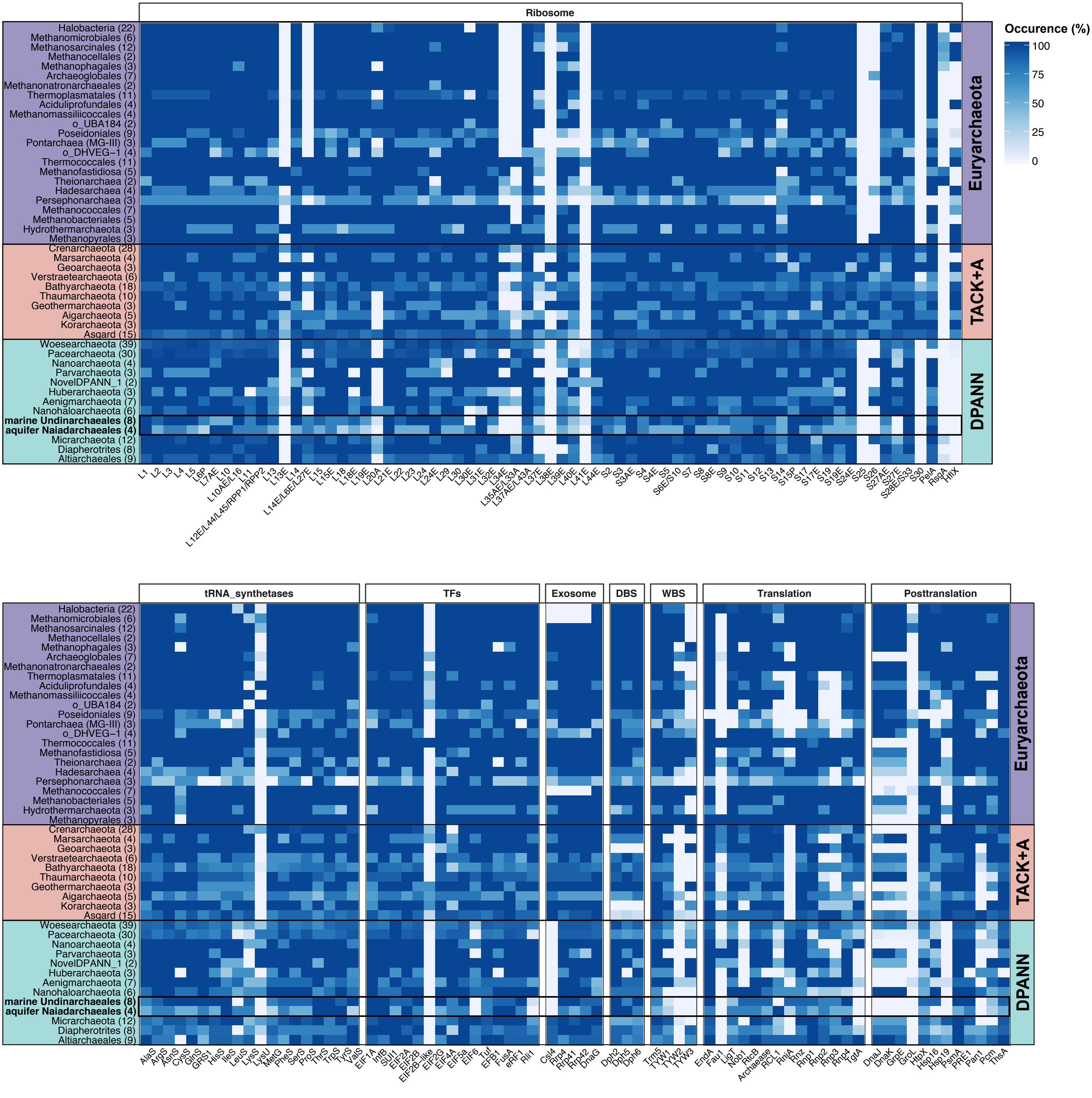


**Supplementary Figure 61 | Presence of key translation-related proteins across major archaeal lineages**. Heatmap of presence/absence patterns of key proteins are summarized across the total number of genomes included in each phylogenetic cluster (shown in percent). TACK + A = TACK + Asgard. DBS = Diphthamide biosynthesis. WBS = Wybutosine biosynthesis. Number in brackets = number of genomes analyzed for each phylogenetic cluster. Supplementary Table 19 lists the proteins used to generate the plot and Supplementary Table 9 lists the raw values.

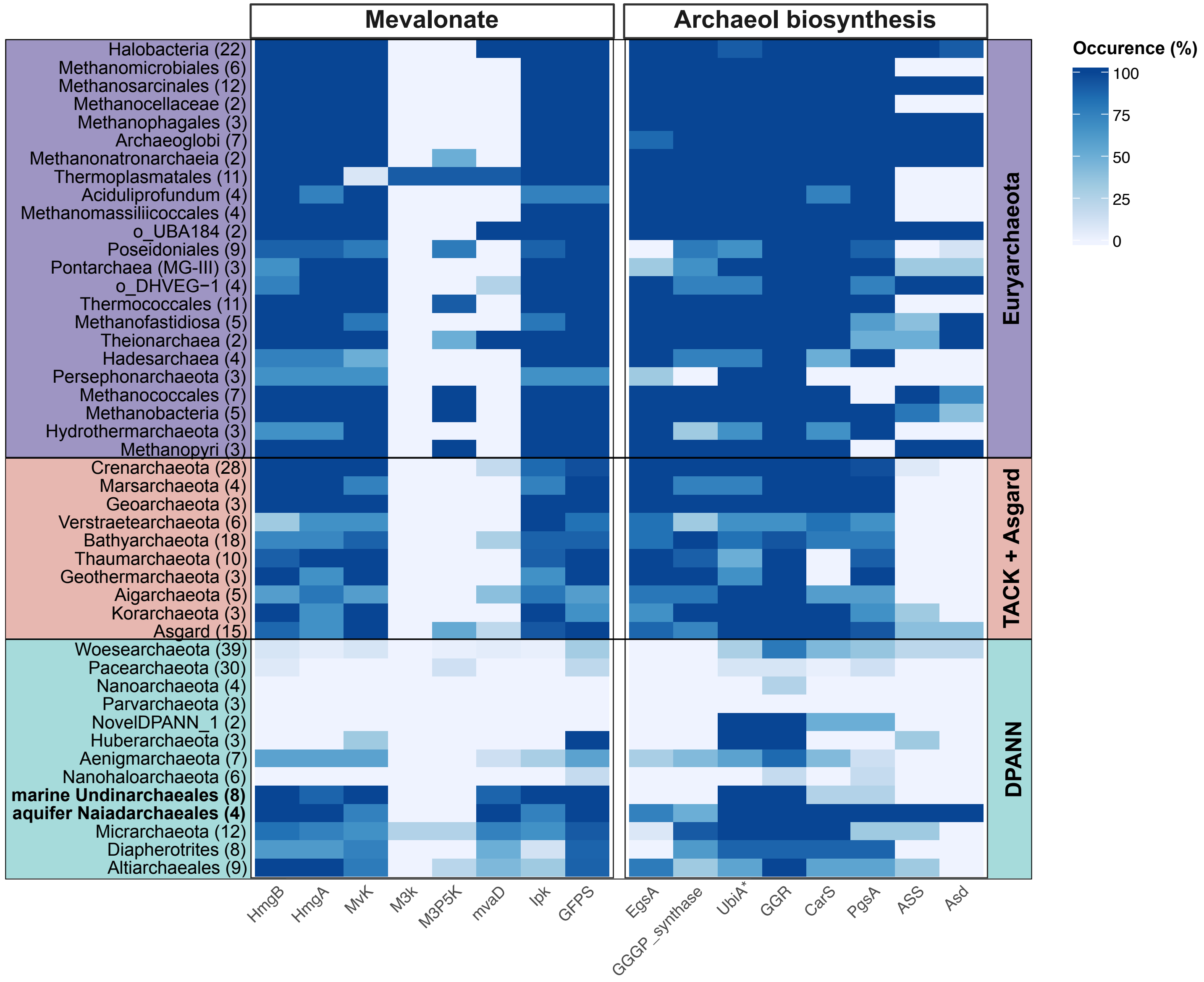


**Supplementary Figure 62 | Presence of key lipid-related proteins across major archaeal lineages.** Heatmap of presence/absence patterns of key proteins are summarized across the total number of genomes included in each phylogenetic cluster (shown in percent). TACK + A = TACK + Asgard. Number in brackets = number of genomes analyzed for each phylogenetic cluster. Supplementary Table 19 lists the proteins used to generate the plot and Supplementary Table 9 lists the raw values. *In Undinarchaeota only the Naidarchaeales UbiA contains a DGGGP synthase domain required for this enzyme to function in lipid biosynthesis.

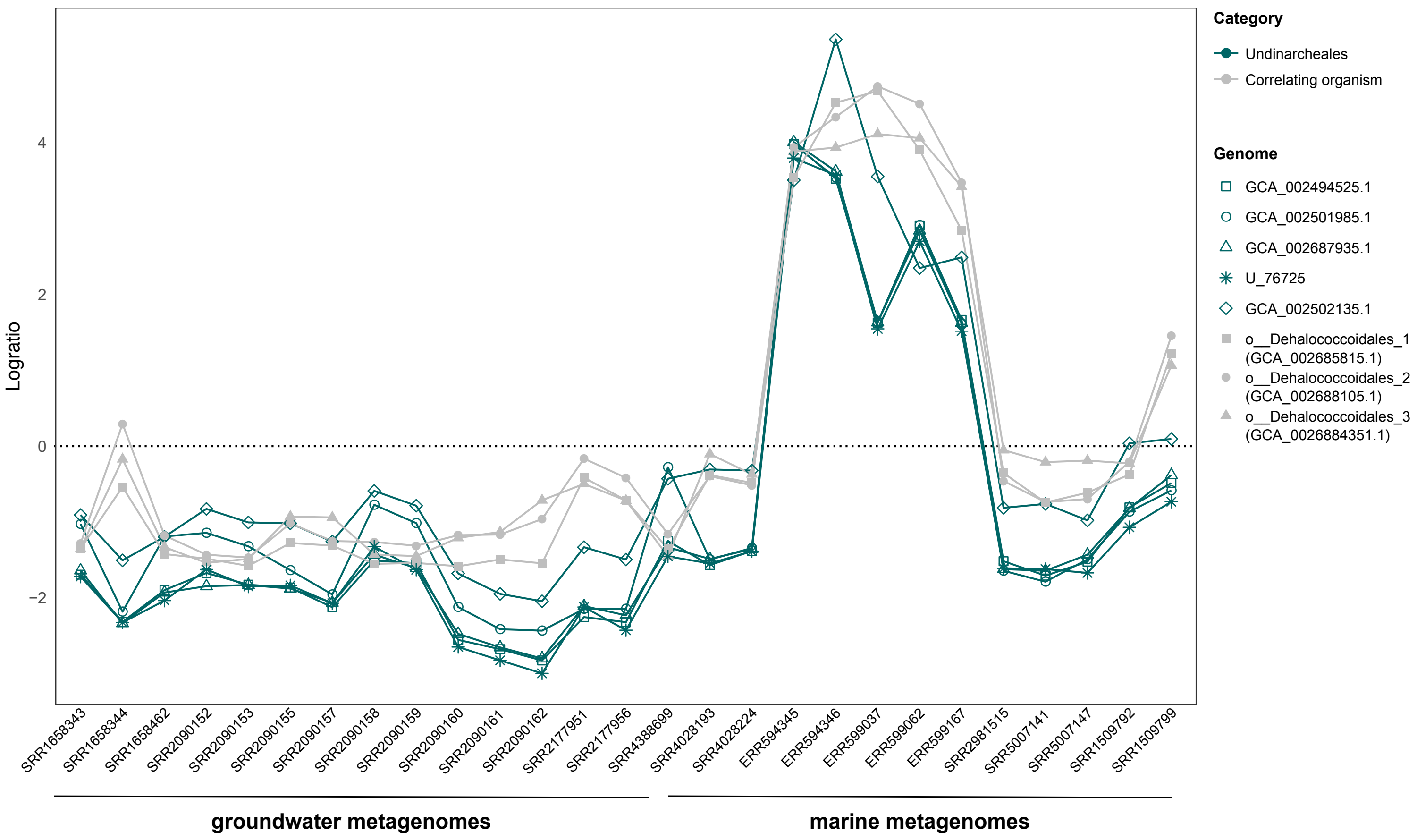


**Supplementary Figure 63 | Determining co-correlations signals of Undinarchaeota with a potential host.** 37 metagenomes containing reads assigned to Undinarchaeota (Supplementary Table 1) were aligned to a reference database of 6,890 archaeal and bacterial genomes. Proportionality was calculated based on normalized relative abundances and centered log-ratio transformation. Genomes shown in this graph were proportional ($\rho \geq 0.9$) to more than one Undinarchaeota MAG and thus were inferred as Undinarchaeota co-correlated.
